# Blind spots in traditional approaches to conservation prioritization in a climate change context

**DOI:** 10.1101/2025.10.01.679788

**Authors:** Luiz Bondi, Beatriz Prado-Monteiro, Luiza FA de Paula, Bruno HP Rosado, Stefan Porembski

## Abstract

Climate change affects biodiversity faster than conservation assessments can be conducted. This issue calls for approaches that support decisions on conservation prioritization, such as species phylogenetic relationships or diversity and endemism metrics. However, these traditional approaches often neglect some important aspects, limiting their effectiveness. We used desiccation-tolerant vascular plants (DT plants) to investigate the effectiveness of such approaches to prioritize species and areas for conservation. We used climate data and modeled the distribution of all DT plants recognized to date to evaluate if species phylogenetic relationships can depict similarities in species sensitivity and exposure to climate change, and to evaluate if centers of diversity and endemism for DT plants can indicate regions prone to climate change. We found that the species phylogenetic relationships weakly explains species sensitivity to climate change, although it can, to some extent, describe trends in species exposure to climate change. We also found that centers of diversity and endemism for DT plants are not necessarily the most prone ones to climate change. We suggest a limited effectiveness of phylogenetic relationships and of diversity and endemism metrics for conservation prioritization, once these approaches might overlook vulnerable species and regions exposed to climate change. We discuss that a better understanding of the mechanisms of diversity would help to identify situations in which closely related species show lower ecological differences than distantly related species, when phylogenetic relationships is a more relevant approach in a conservation context. We also suggest that more efficient conservation strategies in centers of diversity and endemism of DT plants should focus on species sensitivity and adaptive capacity to climate change, rather than the magnitude of climate change.

## 1. Introduction

Climate change is considered one of the main threats to global biodiversity (Thomas *et al*., 2004; Pimm, 2008; Pecl *et al*., 2017). That is because shifts in climate conditions can exceed species’ capacity to cope with them (Dawson *et al*., 2011; Anderson, 2016), promoting species extinctions and declines in species diversity worldwide (Thomas *et al*., 2004; AguirrelGutiérrez *et al*., 2022). To understand the effects of climate change on biodiversity is not a simple task, once the magnitude of changes varies throughout locations and species have different sensitivities and adaptive capacities to climate change (Dawson *et al*., 2011). The problem is that biodiversity declines are occurring before conservation assessments on most species can be conducted (Guénard et al., 2025), motivating efforts for the prioritization of species and areas for conservation (e.g., IUCN, 2012; Myers et al., 2000).

Among the different traditional approaches to identify conservation priorities, we highlight two. The first approach uses species phylogenetic relationships to designate priority species for conservation. The second approach focus on diversity and endemism metrics for the identification of locations with the highest or most unique diversity to appoint priority areas for conservation (by unique we refer to species from to distinctive phylogenetic clades and/or do not occur elsewhere). These approaches are valuable tools for biodiversity conservation because they are based on consistent premises to draw conservation strategies. The first approach relies on the expectation that phylogenetically closely related species tend to share similar requirements and tolerances for environmental conditions due to some degree of phylogenetic inertia (Webb *et al*., 2002; Wiens, 2004). The second approach relies on the objective of protecting the largest number of species and the ecosystem functions supported by them (Purvis et al., 2005; Kier et al., 2009; Rosauer et al., 2009; Winter et al., 2013; Harrison & Noss, 2017).

However, these approaches are not necessarily effective. For example, Hähn et al. (2025) show that species phylogenetic relationships do not always describe species ecological similarities. Similarly, most species-rich regions identified by Barthlott et al. (2005) are not always the one in which the magnitude of shifts in climate is higher according to Bowler et al. (2016). As a consequence, the blind spots of traditional approaches could mislead conservation efforts to species and areas with lower conservation needs. To better understand the effectiveness of those approaches to identify conservation priorities would be particularly important for species that remain largely overlooked in conservation, where the knowledge gap about species vulnerability to climate change is more pronounced.

We argue that desiccation-tolerant vascular plants (DT plants) are good models to discuss the effectiveness of such approaches for species overlooked in conservation. DT plants form a polyphyletic group of plants that can overcome the desiccation of their photosynthetic tissues without losing biomass (i.e., less than 13-20% of protoplasmic water; Oliver *et al*., 2000; Porembski & Barthlott, 2000). Although exhibiting an uncommon response among vascular plants, DT plants have a wide phylogenetic and geographic distribution (Oliver *et al*., 2000; Porembski & Barthlott, 2000). Because of their ability to tolerate desiccation, the impacts of climate change on these species have been poorly discussed (Bondi et al. 2024; Seidou et al., 2025). However, empirical studies have shown that DT plants are in fact affected by changes in environmental constraints, such as temperature and drought regimes (Marks *et al*., 2021). Thus, the lack of studies addressing the conservation aspects of DT is not justified, since these species can in fact be threatened with extinction by climate change. To the best of our knowledge, little is known about to which extent species phylogenetic relationships can explain their ecological similarities or to which extent diversity and endemism metrics can indicate regions most prone to climate change. The conservation of these species becomes particularly relevant when considering that they are regarded as sources of genetic information for many applied purposes, such as increasing drought tolerance of crop species (Farrant et al., 2020).

In this study, we investigate to which extent traditional approaches to identify conservation priorities effectively address the conservation needs of species overlooked in conservation in a climate change context. For that, we used DT plants and evaluated how well their phylogenetic relationships and diversity and endemism metrics can help to identify of priority species and areas for conservation. First, to examine the relevance of species phylogenetic relationships for conservation prioritization, we test the hypothesis that (i) phylogenetically closely related species are more similarly sensitive and exposed to climate change than phylogenetically distantly related species. Then, to examine the relevance of diversity and endemism metrics for conservation prioritization, we test the hypothesis that (ii) centers of diversity and endemism coincide with regions most prone to climate change. We believe that our findings would not only contribute to our understanding about strategies for conservation prioritization of DT plants, but also to different sets of species neglected in conservation, which is very much desired in a climate-changing world.

## 2. Materials and methods

### 2.1. Model system: Desiccation Tolerant Vascular Plants

The desiccation tolerance of vegetative organs has independently re-evolved multiple times in tracheophytes (from fern allies to eudicots), and in each time involving different responses (e.g., some constitutive responses in ferns species contrasting with induced responses in monocots; Oliver *et al*., 2000). In general, their desiccation tolerance capacity is influenced by changes in the desiccation rate, light and temperature during their desiccation and rehydration processes, besides the frequency, intensity, and duration of drought events (Farrant *et al*., 1999; Farrant & Kruger, 2001; Farrant *et al*., 2003; Georgieva *et al*., 2008; Marks *et al*., 2021). Rock outcrops from eastern South America, south/southeastern Africa, Madagascar, and Western Australia are expected to be the centers of diversity for DT plants (Porembski & Barthlott, 2000). But differences in diversity patterns of DT plants are also noticeable (e.g., DT Poaceae are almost absent in South America, while DT Velloziaceae are not found in Australia; Porembski *et al*., 2021). The fact that those regions hold distinct climates might imply that the magnitude of climate change will be different across them.

### 2.2. Climate data

We used climate data to all our analyses and we chose the ones describing for the six main constraints to DT plants’ diversity and distribution, as listed by Marks *et al*. (2021): desiccation rate, light and temperature during their desiccation and rehydration processes, besides the frequency, intensity, and duration of drought events. For that we estimated the (i) vapor pressure deficit (VPD), (ii) solar radiation (SRad), (iii) mean annual temperature (MAT), (iv) drought frequency (DRF), (v) drought intensity (DRI), and (vi) drought length (DRL) as a proxy for desiccation rate, light and temperature during species desiccation and rehydration, besides the frequency, intensity, and duration of drought events, respectively. Here, higher values of VPD, SRad, MAT, DRI, DRL, and DRF describe climates in which the desiccation rate of DT plants is greater, light and temperature is higher during desiccation-rehydration processes, and droughts are more intense, extensive, and frequent.

Firstly, to generate the distribution hypotheses for species and estimate species sensitivity to climate change, we calculated the VPD, SRad, and MAT using historical datasets from the Worldclim v2.1 database (Fick & Hijmans 2017) and the DRF, DRI, and DRL using the one-month dataset from the Standardized Precipitation Evapotranspiration Index (SPEI) database (Vicente-Serrano et al., 2010). While the mean monthly values for the period from January 1970 to December 2000 were considered for Worldclim datasets, the period from January 1901 to December 2018 was considered for the SPEI dataset. The VPD was estimated as the annual mean value of the monthly differences between saturated vapor pressure and actual vapor pressure (Grossiord *et al*., 2020; please see the supporting information for more details). For the actual vapor pressure we used the *vapr* dataset, while the saturated vapor pressure was estimated using the *tmin* and *tmax* datasets and the equation provided by Fick & Hijmans (2017). The SRad and MAT were set by the annual mean of the *srad* and *bio1* datasets, respectively. For DRF, DRI, and DRL, we considered a drought event as composed of a given set of consecutive dry months (i.e. SPEI < 0). The DRF was estimated as the mean count of drought events per year. The DRI was estimated as the whole-period average of drought event intensity, given by the cumulative SPEI for each month within a drought event. At last, DRL was estimated as the whole-period average of the number of consecutive dry months within a drought event. While the Worldclim datasets were selected for providing an accurate estimation of solar radiation and temperature (Fick & Hijmans 2017), the SPEI dataset was chosen for providing a robust estimate of drought (while considering the hydraulic balance between water demand via evapotranspiration and water supply via precipitation; Vicente-Serrano et al., 2010).

Then, to estimate historical climatic variability for VPD, SRad, MAT, DRI, DRL, and DRF, we calculated differences between the average climatic conditions registered for the period from 1958 to 1989 and for the period from 1990 to 2021 (i.e., ΔVPD, ΔSRad, ΔMAT, ΔDRI, ΔDRL, and ΔDRF, respectively). Here we still used the one-month SPEI dataset to calculate ΔDRF, ΔDRI, and ΔDRL, as described above. However, due to our need to access climate data for the abovementioned periods we used the individual-years datasets *vpd, srad, tmin*, and *tmax* from the Terra Climate project (Abatzoglou *et al*., 2018) to estimate the ΔVPD, ΔSRad, and ΔMAT. While the *vpd* and *srad* datasets from Terra Climate were used to estimate ΔVPD and ΔSRad, respectively, ΔMAT was calculated by using the mean value between the maximum and minimum temperature (i.e. from the *tmax* and *tmin* datasets). A higher historical climatic variability (i.e., a higher value of ΔVPD, ΔSRad, ΔMAT, ΔDRI, ΔDRL, and ΔDRF) describes a higher magnitude of shifts in climate a location is likely to undergo.

The average of a minimum 30-year period to estimate average climatic conditions, as recommended by the World Meteorological Organization (https://public.wmo.int/). We used raster grid resolution of 2°30’’ x 2°30’’ for all our analyses.

### 2.3. Species distribution hypotheses

We generated distribution hypotheses for species in order to test both research hypotheses. For that, we first conducted a bibliographic search using the Web of Science search engine (apps.webofknowledge.com) and the key-words combination *“desiccation tolerant” OR “resurrection”) AND (angiosperm* OR pteridophyte* OR lycophyte* OR vascular OR plant*\* to compile a list of DT plants. To improve the species list, we also included additional studies which were not present in the bibliographic search (please see references for Table S1 in Supporting Information). All DT plants reported at the “species” taxonomic level by those scientific studies were considered, and the accepted names followed the Tropicos database (Tropicos.org, 2022). We found 337 DT plants (80 genera and 21 families) reported in 1145 scientific studies (please see Table S1 in Supporting Information).

Then, we obtained the occurrence records for every DT plant using preserved herbarium specimens with available geographic information from the databases (i) GBIF - Global Biodiversity Information Facility (GBIF.org, 2022; please see Table S2 in Supporting Information for more details), (ii) Tropicos, and (iii) Species Link (speciesLink network, 2022). From those records, we removed duplicated, erroneous and uncertain data according to the databases (i) POWO - Plants of the World Online (POWO, 2022), (ii) World Plants (Hassler, 2022), and (iii) Flora e Funga do Brasil (Flora e Funga do Brasil, 2022). We also accepted records in which the precise description of locality or municipality was provided, by using the centroid of the municipality, but removed country and province centroids from our occurrences dataset with the aid of the R package CoordinateCleaner (Zizka et al., 2019). Finally, to avoid the effects of the uneven sampling bias, we reduced multiple records for the species within an area of 1 km-radius to only one occurrence. The 1 km-radius was chosen by considering the premise that most species’ populations occur on isolated rock outcrops.

For the distribution hypotheses, we considered the consensus areas for the species occurrence between the two modeling approaches, whenever it was possible to assess. We used the Maximum Entropy technique as the first modeling approach (MaxEnt; Phillips *et al*., 2004), which was conducted based on a climatic niche perspective. All MaxEnt models were calibrated with the same six variables mentioned above to describe the six main environmental constraints to DT plants (i.e. VPD, SRad, MAT, DRF, DRI, DRL). Since this study makes use of a climatic perspective, we did not consider other important factors for the diversity and distribution of DT plants, such as topo-edaphic conditions. Then, we conducted the Inverse-distance weighted model approach (IDW) to predict every species distribution by a presence-absence interpolation model. We evaluated the predictive power of models from both techniques by the area under the receiver operating characteristic (AUC) after cross-validation using the method of k-means (k=5), in which 10000 random background points were generated. Each MaxEnt and IDW models were produced by at least 50% of consensus between five different random cross-validation routines for the same approach. At last, we generated binary distribution hypotheses for each species, in which individual model thresholds were estimated using the minimum omission rates for true positives and true negatives (i.e., best sensitivity and specificity). For every species, we used raster grids of 2°30’’ x 2°30’’ resolution, expanding the extent of possible occurrence to 5° of latitude and longitude beyond the species’ most external occurrence points.

The MaxEnt technique was chosen due to its ability to identify suitable areas of occurrence for species (Elith *et al*., 2011) with good predictive power, being little affected by the sample size effects and the sort of data which is required (Hernandez *et al*., 2006; Wisz *et al*., 2008; Elith *et al*., 2011; Gogol-Prokurat, 2011; Yackulic *et al*., 2013; Feng *et al*., 2019). The IDW technique was chosen due to its capacity to predict species distribution under a strong spatial autocorrelation and because it provides an alternative hypothesis for the niche-based species distribution models, which can overestimate species distribution measures to areas beyond species dispersal capacity (Diniz-Filho *et al*., 2003; Pearson & Dawson, 2003; Roberts *et al*., 2004; Bahn & Mcgill, 2007; Elith *et al*., 2011). Putting aside biotic interactions, through the combination of the mentioned species distribution model techniques, the assessment of species’ actual distribution is supposed to be more realistic once it depends on both abiotic suitability and habitat accessibility (Soberón & Peterson, 2005; Peterson, 2009). However, those modeling techniques could not be performed for species with less than five observation points after rarefying occurrences. To get around this problem, we estimated the species distribution by applying the method of the Circular Area with a radius of 50 km (Ca_50_) for species with less than five observation points, as proposed by Hijmans & Spooner (2001). The species distribution models were performed for 316 species, and 20 species had less than 5 valid occurrences, so only the Ca_50_ was used to assess their distribution hypotheses. We did not produce a distribution hypothesis for *Tripogon polyanthus* due to the lack of valid occurrences for these species.

### 2.4. Phylogenetic relationships depicting priority species for conservation

To test the hypothesis that phylogenetically closely related species are more similarly sensitive and exposed to climate change, we considered the phylogenetic hypothesis provided by Jin & Qian (2019; *Scenario 3*) to measure the phylogenetic signal (i.e., the degree of phylogenetic constraint in species resemblance; Molina-Venegas et al., 2017) on species relationship with environmental constraints. The phylogenetic signal on species ecological patterns was estimated using Pagel’s λ (Harvey & Pagel, 1991) with aid of *phylosignal* R package (Keck et al., 2016). This metric was chosen for its low error rate at detecting variation as expected by Brownian motion, and due to its robustness under high polytomy, which is found in the used phylogenetic hypothesis (Münkemüller et al., 2012).

To evaluate if species are similarly sensitive to climate change we calculated the relative importance of VPD, SRad, MAT, DRF, DRI, and DRL in explaining the species distribution. That is because the relative importance of a given environmental variables in explaining a species distribution increases as the species sensitivity to changes in this environmental variable increases. Those values we obtained from a Jackknife test on MaxEnt models. Here, the species distribution was repeatedly modelled leaving out one variable at a time, using the difference between the models performance (i.e., AUC values with and without each variable) to describe their importance for the distribution of a given species. A higher relative importance of a given environmental constraint (i.e., higher difference between models performance) would describe the higher sensitivity of the species to changes in this factor. Then, if environmental constraints show similar relative importance to a pair of species, they can be expected to be similarly sensitive to climate change.

To evaluate if species are similarly exposed to climate change, we calculated mean values of historical climatic variability within their distribution areas. We used ΔVPD, ΔSRad, ΔMAT, ΔDRF, ΔDRI, and ΔDRL to describe historical climatic variability. Higher historical climatic variability would describe greater magnitude of shifts in climate species might encounter within their distribution ranges. If a pair of species is subjected to similar historical climatic variability, in every used dimension, they can be expected to be similarly exposed to climate change.

### 2.5. Centers of diversity and endemism depicting priority areas for conservation

To test the hypothesis that centers of diversity and endemism coincide with regions more prone to climate change, we firstly identified the centers of diversity and endemism using different diversity and endemism metrics. While centers of diversity were determined by combining regions of highest species richness and phylogenetic diversity, centers of endemism were highlighted by combining regions with highest endemism richness and phylogenetic endemism. Although we recognize that estimations of diversity are highly dependent on the spatial scale, we only used a 2°30’’ x 2°30’’ resolution for simplification.

To estimate the centers of diversity for DT plants, we first overlapped the distribution hypotheses of every DT plant and used the cumulative species count in each spatial unit (i.e., grid cell) to measure species richness. A high species richness is encountered in locations where many species overlap their distribution. Then, we estimated the phylogenetic diversity for the same spatial units. Here, we calculated the Rao index for α-diversity (Rao, 1982) of the phylogenetic trees (using the phylogenetic hypothesis provided by Jin & Qian, 2019) constructed for the species that share their occurrence in the same spatial unit. The Rao index was chosen for providing robust measurement of the species redundancy (de Bello et al., 2010), from a phylogenetic perspective in the case of our study. By last, we grouped spatial units into 16 interval classes according to scores of species richness and phylogenetic richness. Those scores represented values from 0-25%, 25-50%, 50-75%, and 75-100% in relation to the maximum score of species richness and phylogenetic richness registered worldwide. Spatial units that scored 75-100% of maximum species richness and phylogenetic diversity were accounted as centers of diversity.

To estimate the centers of endemism for DT plants, we followed the same logic. We first estimated the endemism richness by weighting the value of each species presence in a spatial unit by its inverse range size (e.g., presence in a spatial unit equals to 1 over the number of spatial units the species is found) and summing up the presence values of every species that co-occur in the same spatial unit, as performed by Kier *et al*., (2009). Locations with higher endemism richness accumulated a higher number of species with restricted geographical ranges. Then, we estimated the phylogenetic endemism in the same spatial units, following Rosauer *et al*. (2009). This metric considers the species’ endemism, the species phylogenetic distance to the closely related taxa, and the range of the specie’s closely related taxa. Regions with higher phylogenetic endemism gathered more species in which they are spatially and phylogenetically restricted. Spatial units that scored 75-100% of maximum endemism richness and phylogenetic endemism were accounted as centers of endemism.

At last, we correlated the diversity and endemism scores in each spatial unit with the historical climatic variability for the same spatial unit. ΔVPD, ΔSRad, ΔMAT, ΔDRF, ΔDRI, and ΔDRL were used as a proxy for the magnitude of shifts in climate a location will undergo owing to climate change. To allow an comparison between the magnitude of shifts in climate with centers of diversity and endemism, we standardized and summed the values for every climatic variable using the function *decostand* from the package vegan (Oksanen et al., 2022) All geographic information system routines, descriptive, and statistical analyses, besides all graphical representations, were conducted in R software 4.2.0 (R Core Team, 2022; Table S5).

## 3. Results

### 3.1. Phylogenetic relationships depicting priority species for conservation

In general, we found a weak phylogenetic signal on the extent to which environmental constraints influence species distribution (i.e., our proxy for species sensitivity to climate change), as the relative importance of environmental constraints was highly variable across and within clades (Fig.1). While the strongest (yet weak) signal was registered to VPD (Pagel’s λ = 0.37, *p*-value = 0.001), no significant phylogenetic signal was found for the influence of SRad on species distribution. Regarding the mean historical climatic variability recorded for species (i.e., our proxy for species exposure to climate change), the strength of the phylogenetic signal on the magnitude of shifts in climate species might encounter within their distribution ranges was variable (Fig.2). While a strong phylogenetic signal was registered to ΔMAT (Pagel’s λ = 0.92, *p*-value = 0.001) and ΔVPD (Pagel’s λ = 0.78, *p*-value = 0.001), a weaker phylogenetic signal was registered to ΔSRad (Pagel’s λ = 0.55, *p*-value = 0.001) and ΔDRF (Pagel’s λ = 0.2, *p*-value = 0.003). No significant phylogenetic signal was found for ΔDRL and ΔDRI.

**Fig. 1.**
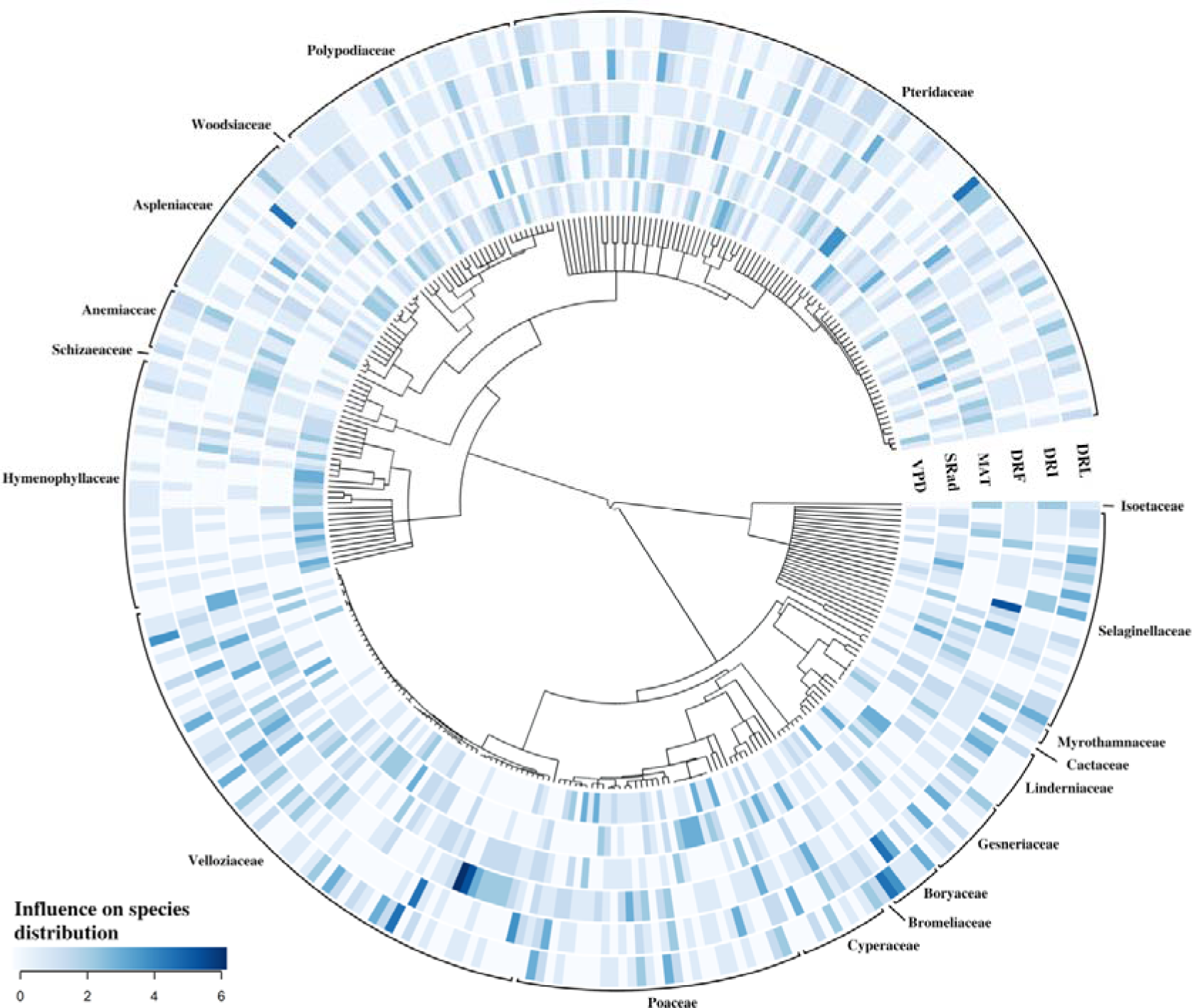
Species sensitivity to climate change (described by the environmental constraints influence species distribution) under a phylogenetic perspective. VPD – vapor pressure deficit; SRad – solar radiation; MAT – mean annual temperature; DRF – drought frequency; DRI – drought intensity; DRL – drought length.

**Fig. 2.**
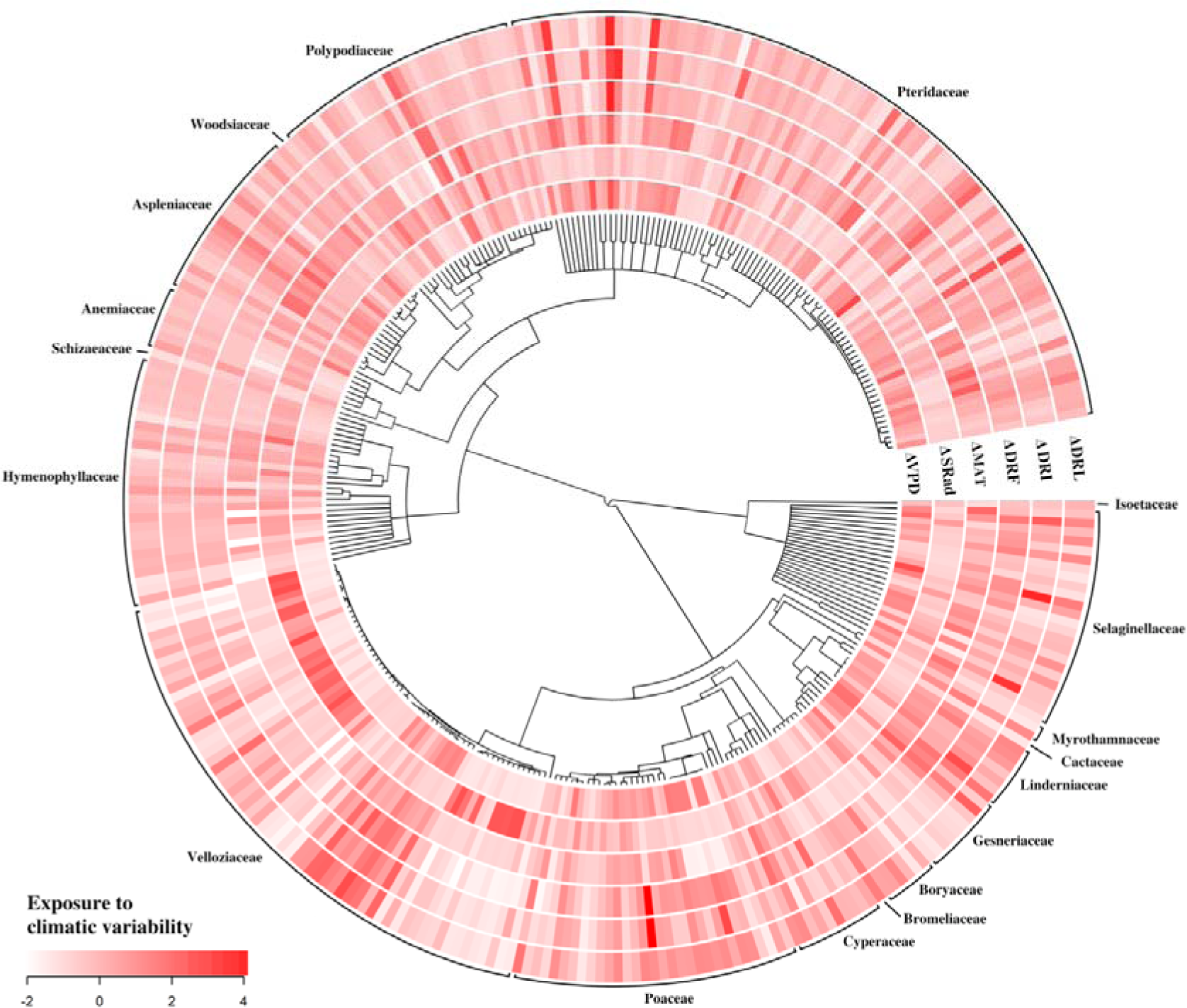
Species exposure to climate change (described by the mean historical climatic variability recorded for within species distribution range) under a phylogenetic perspective. ΔVPD – shifts in vapor pressure deficit; ΔSRad – shifts in solar radiation; ΔMAT – shifts in mean annual temperature; ΔDRF – shifts in drought frequency; ΔDRI – shifts in drought intensity; ΔDRL – shifts in drought length.

### 3.2. Centers of diversity and endemism depicting priority areas for conservation

We found centers of diversity for DT plants worldwide in the Americas, Africa and Madagascar (Fig.3; please see Fig.S1 in Supporting Information), while centers of endemism for DT plants were found in all continents, except Antarctica (please see Fig.S2 in Supporting Information).

**Fig. 3.**
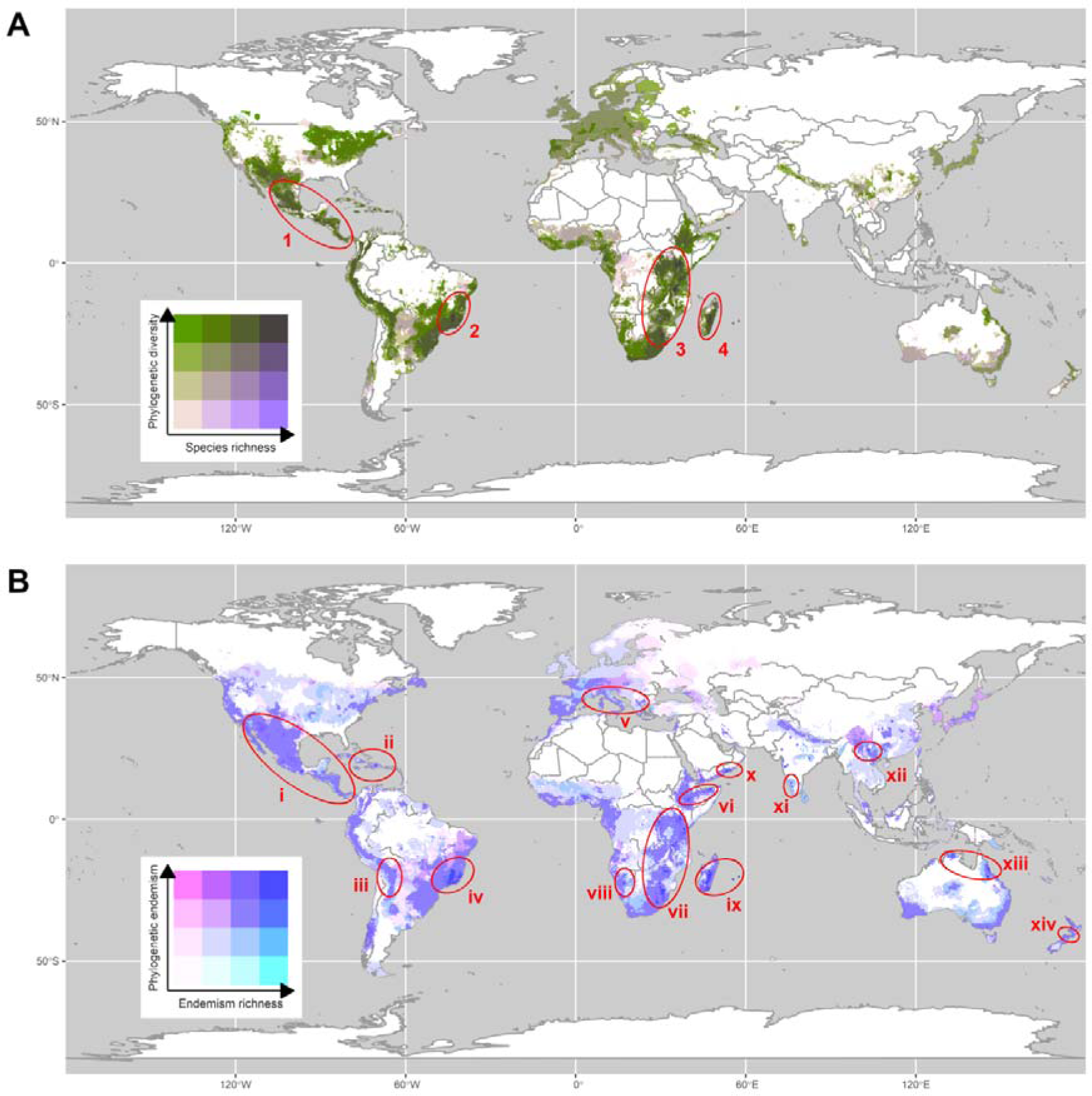
Centers of diversity and endemism for desiccation-tolerant vascular plants. A - Centers of diversity including (1) the Central American Cordillera, (2) Cadeia do Espinhaço – Sugarloaf Land, (3) East African Rift – Eastern Highlands – Drakensberg, and (4) Malagasy Central High Plateau. Centers of endemism including locations in the (i) Mexican – Central American Cordilleras, (ii) Caribbean Islands, (iii) Bolivia’s Cordillera Oriental – Quebrada de Humahuaca, (iv) Cadeia do Espinhaço – Sugarloaf Land, (v) Provence – Ionian – Belasica Range, (vi) Ogo and Bale Mountains, (vii) East African Rift – Eastern Highlands – Drakensberg, (viii) Khomas Hochland, (ix) Madagascar – Mascarene Islands, (x) Center-southern Arabian Mountains, (xi) Western Ghats, (xii) Yunnan province, (xiii) Northern Territory – Wet Tropics, and (xiv) Richmond Range.

High species diversity and endemism was not always registered in locations in which historical climatic variability was high (Fig.4). Instead, we could observe that species richness tend to increase with lower historical climatic variability, but only when considering VPD, DRF, DRI, and DRL. No clear trend in endemism richness could be observed in function of historical climatic variability, although regions with high endemism richness are less prone to shifts in VPD, DRF, DRI, and DRL. When phylogeny was taken into account, we could not identify increase or decrease trends in diversity and endemism alongside with historical climatic variability.

**Fig. 4.**
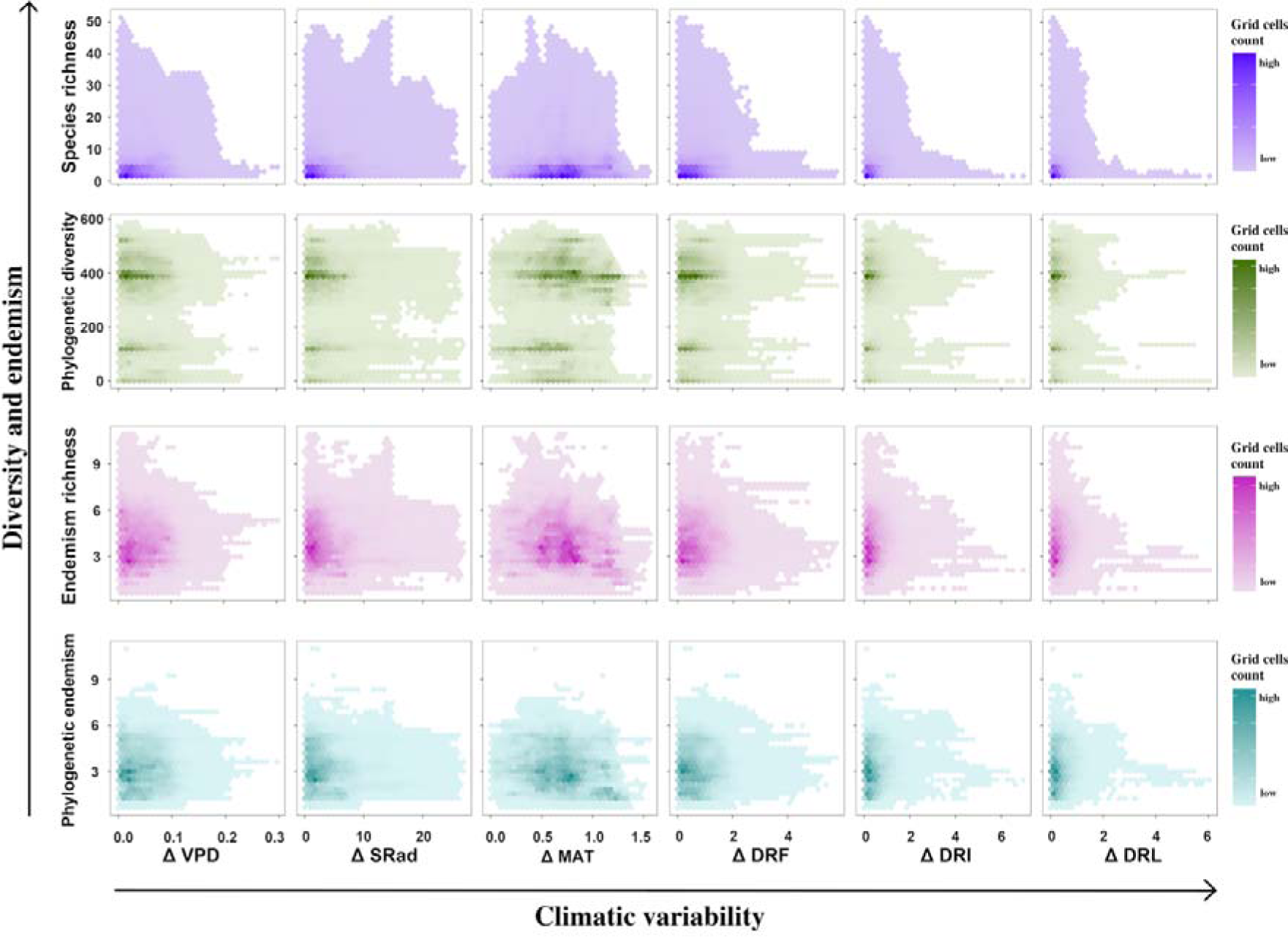
Trends in diversity and endemism of desiccation-tolerant vascular plants alongside with historical climatic variability. The diversity scores were described by species richness and phylogenetic diversity, while the endemism scores were described by endemism richness and phylogenetic endemism. ΔVPD – shifts in vapor pressure deficit; ΔSRad – shifts in solar radiation; ΔMAT – shifts in mean annual temperature; ΔDRF – shifts in drought frequency; ΔDRI – shifts in drought intensity; ΔDRL – shifts in drought length.

## 4. Discussion

### 4.1. Conservation strategies should not rely on species phylogenetic relationships

We could not completely confirm the hypothesis that phylogenetically closely related species are more similarly sensitive and exposed to climate change than phylogenetically distantly related species. In fact, species phylogenetic relationships does not always predict species-environment relationships (Hähn et al., 2025), as it did not predict DT plants sensitivity to climate change but the species exposure to it. Thus, conservation actions that rely on species phylogenetic relationships might correctly identify exposure trends among closely related species that cluster in space, but might neglect that these species may have different sensitivities to climate change.

One factor that could explain our findings is the ecological similarity among species that share little evolutionary history. This similarity might be promoted by the environment shaping advantageous plant responses to cope with common constraints (Weiher & Keddy, 1995; Hille Ris Lambers et al., 2012; de Bello et al., 2015; Kraft *et al*., 2015). Hence, coexisting distantly related species would be more similarly sensitive to climate change than closely related species due to their convergent responses to the environment and co-occurrence. This can be illustrated by *Vellozia variabilis* (Velloziaceae) and *Selaginella convoluta* (Selaginellaceae) which were found ecologically similar within their overlapping distribution. It is likely that climate change affects those species in a similar way if they exhibit equivalent responses to changes.

Alternatively, our results might have identified the ecological dissimilarity among closely related taxa, which can be explained by a limiting similarity (sensu MacArthur & Levins, 1967) required for species in sympatry. Closely related species might exhibit important niche differences due to stabilizing mechanisms that enable their coexistence (MacArthur & Levins, 1967; Chesson, 2000). The logic is, due to competitive exclusion principle, two closely related species must exhibit enough niche differences to achieve a stable coexistence (Hardin 1960; Tilman, 1982; Chesson, 2000; Cadotte et al., 2017). As a consequence, they show different sensitivities to climate change even when exposed to the same shifts in climate. This could be exemplified by Velloziaceae species, which showed a high intra-generic divergence on the environmental constraints that influence their distribution, but are exposed to similar climatic variability due to high geographical overlap (Fig.1 and 2).

Yet, we should not disregard the fact that closely related sympatric species can still be similarly sensitive to climate change. That would happen in two situations. First, when coexisting species have small fitness differences, that is, when they have equivalent ability to survive, grow, and reproduce under specific resource availability (Tilman, 1982; Chesson, 2000; Mayfield & Levine, 2010). Alternatively, closely related species can coexist when intraspecific competition exceeds interspecific competition, that is, when the negative effects of competition are greater among individuals of a same species than between individuals of different species (Chesson, 2000; Mayfield & Levine, 2010). Thus, coexistence mechanisms can also cause the ecological similarities between among closely related species (Kraft *et al*., 2015).

We believe that conservationists and decision-makers would greatly benefit from a sharpened awareness of the mechanisms of diversity. Due to the nature of our data, we could not evaluate the species coexistence. We believe that future studies explicitly designed to investigate these mechanisms could not only contribute to identify priority species to conservation, but also provide important insights for further conservation efforts. For instance, many DT species occur on inselbergs, which are rock outcrops shaped by similar ecological processes and that function as terrestrial island-like ecosystems (Porembski & Barthlott, 2000). Because of that, it can be observed a high rate of species turnover across inselbergs alongside with the cluster of these species in a functional space (Paula et al., 2020). That is, closely related species replace each other across different inselbergs, but their niche differences are small. It suggests that, if fitness differences of closely related species are big and intraspecific competition is weak, species capacity to track suitable environments in climate change scenarios is limited by both dispersal limitation and competitive exclusion of species. That would not only increase the vulnerability of species with phylogenetic equivalents, but also hinder intensive intervention actions on these species, such as assisted migration. We believe that the use of species phylogenetic relationships is helpful in such situations.

It is also noteworthy to mention that the time scale in which climate change operates is different from the time scale in which evolutionary processes operate (de Bello *et al*., 2015; Anderson, 2016). Processes important to generate spatial patterns under larger time scales, are less important under shorter time scales (Blois et al., 2013; Lovell et al., 2023). As a consequence, species phylogenetic relationships have a reduced importance in identifying priority species for conservation in climate change context. We believe that functional traits could bridge evolutionary and ecological process, and produce relevant insights in this respect. For instance, the cuticle is absent in leaves of Hymenophyllaceae species unfolding a consisten high sensitivity of Hymenophyllaceae species to changes in desiccation rate (VPD scored a relative importance of 57 % ± 4.1 to explain their distribution). Since this trait is conserved within Hymenophyllaceae and can describe a lower capacity of plants to cope with changes in VPD, species phylogenetic relationships becomes a powerful tool to identify priority species for conservation when great shifts in VPD are observed. However, traits are not always conserved within phylogenetic lineages and the degree of their “functionality” varies as they do not always determine plants’ fitness (Losos, 2008; Shipley et al., 2016). More studies are still needed to better understand in which situations functional approaches can mechanistically explain the relevance of species phylogenetic relationships in a conservation context.

### 4.2. Centers of diversity and endemism for desiccation-tolerant vascular plants do not always coincide with regions more prone to climate change

Our results did not support the hypothesis that centers of diversity and endemism coincide with regions most prone to climate change. The general trend is that regions with high species and endemism richness are less prone to climate change. This might indicate that the evolutionary processes that drove the diversity of DT plants in those regions are related to long-term climatic evenness. The problem is that species that evolved under long-term climatic evenness are expected to be less tolerant to environmental changes when compared to species from regions with higher historical climatic variability (Willig *et al*., 2003; Fine, 2015). It means that despite being less exposed to climate change, species found in centers of diversity and endemism for DT plants might be more sensitive to changes and exhibit a lower adaptive capacity to changes. Thus, for those regions, we need conservation assessments with a clearer focus on the species sensitivity and adaptive capacity, because even little shifts in climate might exceed their capacity to cope with change.

The inverse reasoning might be applied for regions poorer in species and more prone to climate change. Species from regions with higher historical climatic variability might better cope with changes when compared with species from regions of long term climatic evenness. Yet, shifts in climate can be big enough to still exceed their capacity to cope with changes. Losing such species could mean losing unique evolutionary solutions to deal with drought. This is more special when we consider that functional redundancy is expected to decrease in less species-rich locations, and the lower functional redundancy might reduce the ecological stability of plant communities (Biggs *et al*., 2020). For example, the local extinction of one species might represent the loss of its function if not properly compensated by functionally redundant species (Suding *et al*., 2008). Approaches that rely on diversity and endemism metrics tend to fail to recognize such important aspects of plant communities and consequently overlook the relevance of biodiversity “coldspots” (Kareiva & Marvier, 2003; Allan *et al*., 2022). This can be illustrated by northwestern parts of Brazilian cerrado and eastern parts of Angolan Miombo woodlands, which are prone to climate change but included in a biome historically neglected in conservation (i.e., Tropical and subtropical grasslands, savannas, and shrublands; Dobson et al., 2022). Future conservation initiatives should not neglect less fashionable geographical areas, and monitor how the magnitude of climate change can lead to population declines.

It is noteworthy to mention some limitations of our study. For instance, low species richness in some regions might reflect the geographical bias in desiccation tolerance studies. For example, there is a geographical bias in desiccation tolerance studies (Tebele et al., 2021) and this bias might reflect the low species richness of DT plants in regions rich in vascular plants (e.g., from Ecuador to Costa Rica or in Borneo and Papua New Guinea as identified by Barthlott *et al*., 2005; Kier *et al.,* 2005; Kreft *et al.,* 2008). Besides, our study did not account for the accuracy of space-for-time substitutions, population declines crossing a threshold in which species cannot maintain viable populations, and the existence of stochastic events that are omitted in 30-years climatic means. We believe that future studies that address these knowledge gaps could significantly improve our capacity to identify priority areas for the conservation of these species.

## 5. Conclusion

We suggest a limited effectiveness of traditional approaches to identify conservation priorities in addressing the conservation needs of DT plants in a climate change context. It is possible that such approaches show the same limitations for the conservation prioritization of other species overlooked in conservation. We believe that phylogeny is less relevant in a conservation context when closely related species show higher ecological differences than distantly related species, which was the case of DT plants. This can be caused by the environment or coexistence mechanisms promoting the ecological similarity among distantly related species or the ecological dissimilarity among closely related species. In our view, investigating the mechanisms of diversity could help us to identify such situations and improve our understanding about the role of species phylogenetic relationships for conservation prioritization.

Diversity and endemism metrics might also have a limited effectiveness for determining priority areas for conservation. We believe that identifying centers of diversity and endemism loses relevance in a conservation context when the magnitude of shifts in climate is taken into account. That is possibly because species found in the centers of diversity and endemism evolved in contexts of historical climatic evenness, becoming more sensitive and exhibiting lower adaptive capacity to climate change. We argue that conservation efforts in these regions focus on intrinsic factors to species (i.e., species sensitivity and adaptive capacity), while we do not neglect the extrinsic factors to species (i.e., species exposure to climate change) in regions with fewer species but with unique evolutionary histories.

## DATA ACCESSIBILITY STATEMENT

All datasets generated during and/or analyzed during the current study are available in the Zenodo repository (https://doi.org/10.5281/zenodo.7607611)

## AUTHOR CONTRIBUTIONS

L.B., B.P-M., L.F.A.P., B.H.P.R., and S.P. conceived the idea; L.B. and B.P-M. collected and analyzed the data, and L.B. wrote the manuscript. All authors contributed critically to the drafts and gave final approval for publication.

## ACKNOWLEDGMENTS

The authors thank Aboubacar Zon, Fernando Palma, Konrad Schultz, Luiza Castro, Michael Konietzka, Miguel Inácio, and Smrithy Vijayan for valuable discussions. The authors are grateful for the support from the Deutscher Akademischer Austauschdienst (DAAD; grant number 57440921), Fundação Carlos Chagas Filho de Amparo à Pesquisa do Estado do Rio de Janeiro (FAPERJ; grant number E-26/200.401/2019), and Coordenação de Aperfeiçoamento de Pessoal de Nível Superior (CAPES; grant number 88887.569558/2020-00).

