## Supporting Information for "Blind spots in traditional approaches to conservation prioritization in a climate change context"

The following Supporting Information is available for this article:

**Information about how climate data were calculated**

We used datasets from the Worldclim v2.1 database (Fick & Hijmans 2017) to estimate VPD, SRad and MAT. For the VPD, we calculated the annual mean values of the monthly differences between saturated vapor pressure and actual vapor pressure (Grossiord et al., 2020) using the equation provided by Fick & Hijmans (2017; Eq. 1).

(Eq. 1)

$$VPD={12}^{-1}\times\sum_{i=1}^{i=12} {es}_{i}-{ea}_{i}$$

$${es}_{i}=2^{-1}\times\left( {es}_{i}^{max}-{es}_{i}^{min} \right)$$

$${es}_{i}^{min}=0.611\times{10}^{\frac{7.5\cdot T_{i}^{min}}{\left( 237.7+T_{i}^{min} \right)}}$$

$${es}_{i}^{max}=0.611\times{10}^{\frac{7.5\cdot T_{i}^{max}}{\left( 237.7+T_{i}^{max} \right)}}$$

where, *i* is a given month of the year, *es* is the saturated vapour pressure, *ea* is the actual vapor pressure (*vapr* dataset), and *T^min^* and *T^max^* are minimum and maximum temperatures (*tmin* and *tmax* datasets), respectively. For SRad and MAT, we calculated the annual mean values of *srad* and *bio1* datasets, respectively.

We used the dataset from Standardized Precipitation Evapotranspiration Index (SPEI) database (*spei01*; Vicente-Serrano et al., 2010) to estimate DRF, DRI, and DRL. In this study, a drought event is composed of a given set of consecutive dry months (i.e. SPEI < 0). We estimated the DRF as the mean count of drought events per year (Eq. 2); while DRI was calculated as the whole-period average of drought event intensity (i.e., the cumulative SPEI for each month within a drought event; Eq. 3) and DRL as the whole-period average of the number of consecutive dry months within a drought event (Eq. 4).

(Eq. 2)

$$DRF=n$$

(Eq. 3)

$$DRI=n^{-1}\times\sum_{j=1}^{j=n} {DRI}_{j}$$

$${DRI}_{j}=\sum_{i=1}^{i=n} {SPEI}_{i}$$

(Eq. 4)

$$DRL=n^{-1}\times\sum_{i=1}^{i=m} {dm}_{i}$$

where, *i* is a given month, *j* is a given drought event, *n* is the number of drought events during the period considered, *m* is the number of consecutive dry months within a drought event, and *dm* is a dry month.

**Table S1**Desiccation-tolerant vascular plants and their botanical families.

| Species | Botanical family |
| --- | --- |
| *Anemia ferruginea* Kunth | Anemiaceae |
| *Anemia flexuosa* (Savigny) Sw. | Anemiaceae |
| *Anemia mexicana* Klotzsch | Anemiaceae |
| *Anemia rotundifolia* Schrad. | Anemiaceae |
| *Anemia tomentosa* (Savigny) Sw. | Anemiaceae |
| *Anemia villosa* Humb. & Bonpl. ex Willd. | Anemiaceae |
| *Mohria caffrorum* (L.) Desv. | Anemiaceae |
| *Asplenium adiantum-nigrum* L. | Aspleniaceae |
| *Asplenium aethiopicum* (Burm. f.) Bech. | Aspleniaceae |
| *Asplenium ceterach* L. | Aspleniaceae |
| *Asplenium cordatum* (Thunb.) Sw. | Aspleniaceae |
| *Asplenium dalhousiae* Hook. | Aspleniaceae |
| *Asplenium friesiorum* C. Chr. | Aspleniaceae |
| *Asplenium megalura* Hieron. | Aspleniaceae |
| *Asplenium monanthes* L. | Aspleniaceae |
| *Asplenium obovatum* Viv. | Aspleniaceae |
| *Asplenium praegracile* Rosenst. | Aspleniaceae |
| *Asplenium pringlei* Davenp. | Aspleniaceae |
| *Asplenium ruta-muraria* L. | Aspleniaceae |
| *Asplenium rutifolium* (P.J. Bergius) Kunze | Aspleniaceae |
| *Asplenium sandersonii* Hook. | Aspleniaceae |
| *Asplenium septentrionale* (L.) Hoffm. | Aspleniaceae |
| *Asplenium theciferum* (Kunth) Mett. | Aspleniaceae |
| *Asplenium trichomanes* L. | Aspleniaceae |
| *Asplenium uhligii* Hieron. | Aspleniaceae |
| *Pleurosorus rutifolius* Fée | Aspleniaceae |
| *Borya constricta* Churchill | Boryaceae |
| *Borya inopinata* P.I. Forst. & E.J. Thomps. | Boryaceae |
| *Borya mirabilis* Churchill | Boryaceae |
| *Borya nitida* Labill. | Boryaceae |
| *Borya scirpoidea* Lindl. | Boryaceae |
| *Borya septentrionalis* F. Muell. | Boryaceae |
| *Borya sphaerocephala* R. Br. | Boryaceae |
| *Pitcairnia lanuginosa* Ruiz & Pav. | Bromeliaceae |
| *Blossfeldia liliputana* Werderm. | Cactaceae |
| *Afrotrilepis pilosa* (Boeckeler) J. Raynal | Cyperaceae |
| *Coleochloa abyssinica* (Hochst. ex A. Rich.) Gilly | Cyperaceae |
| *Coleochloa microcephala* Nelmes | Cyperaceae |
| *Coleochloa pallidior* Nelmes | Cyperaceae |
| *Coleochloa setifera* (Ridl.) Gilly | Cyperaceae |
| *Microdracoides squamosus* Hua | Cyperaceae |
| *Trilepis ciliatifolia* T. Koyama | Cyperaceae |

**Table S1**(continued)

| *Trilepis lhotzkiana* Nees ex Arn. | Cyperaceae |
| --- | --- |
| *Trilepis microstachya* (C.B. Clarke) H. Pfeiff. | Cyperaceae |
| *Davallia angustata* Wall. ex Hook. & Grev. | Davalliaceae |
| *Elaphoglossum acrostichoides* (Hook. & Grev.) Schelpe | Dryopteridaceae |
| *Elaphoglossum petiolatum* (Sw.) Urb. | Dryopteridaceae |
| *Elaphoglossum piloselloides* (C. Presl) T. Moore | Dryopteridaceae |
| *Boea hygrometrica* (Bunge) R. Br. | Gesneriaceae |
| *Boea hygroscopica* F. Muell. | Gesneriaceae |
| *Damrongia clarkeana* (Hemsl.) C. Puglisi | Gesneriaceae |
| *Haberlea rhodopensis* Friv. | Gesneriaceae |
| *Oreocharis mileensis* (W.T. Wang) Mich. Möller & A. Weber | Gesneriaceae |
| *Paraboea crassifolia* (Hemsl.) B.L. Burtt | Gesneriaceae |
| *Paraboea rufescens* (Franch.) B.L. Burtt | Gesneriaceae |
| *Ramonda myconi* (L.) Rchb. | Gesneriaceae |
| *Ramonda nathaliae* Pančić & Petrovič | Gesneriaceae |
| *Ramonda serbica* Pančić | Gesneriaceae |
| *Cardiomanes reniforme* (G. Forst.) C. Presl | Hymenophyllaceae |
| *Crepidomanes chevalieri* (Christ) Ebihara & Dubuisson | Hymenophyllaceae |
| *Crepidomanes frappieri* (Cordem.) J.P. Roux | Hymenophyllaceae |
| *Crepidomanes inopinatum* (Pic. Serm.) J.P. Roux | Hymenophyllaceae |
| *Crepidomanes melanotrichum* (Schltdl.) J.P. Roux | Hymenophyllaceae |
| *Didymoglossum erosum* (Willd.) J.P. Roux | Hymenophyllaceae |
| *Hymenoglossum cruentum* (Cav.) C. Presl | Hymenophyllaceae |
| *Hymenophyllum capillare* Desv. | Hymenophyllaceae |
| *Hymenophyllum caudiculatum* Mart. | Hymenophyllaceae |
| *Hymenophyllum dentatum* Cav. | Hymenophyllaceae |
| *Hymenophyllum fucoides* (Sw.) Sw. | Hymenophyllaceae |
| *Hymenophyllum hirsutum* (L.) Sw. | Hymenophyllaceae |
| *Hymenophyllum kuhnii* C. Chr. | Hymenophyllaceae |
| *Hymenophyllum peltatum* (Poir.) Desv. | Hymenophyllaceae |
| *Hymenophyllum plicatum* Kaulf. | Hymenophyllaceae |
| *Hymenophyllum polyanthos* (Sw.) Sw. | Hymenophyllaceae |
| *Hymenophyllum sanguinolentum* (G. Forst.) Sw. | Hymenophyllaceae |
| *Hymenophyllum splendidum* Bosch | Hymenophyllaceae |
| *Hymenophyllum tunbrigense* (L.) Sm. | Hymenophyllaceae |
| *Polyphlebium borbonicum* (Bosch) Ebihara & Dubuisson | Hymenophyllaceae |
| *Trichomanes bucinatum* Mickel & Beitel | Hymenophyllaceae |
| *Trichomanes capillaceum* L. | Hymenophyllaceae |
| *Trichomanes diaphanum* Kunth | Hymenophyllaceae |
| *Trichomanes polypodioides* L. | Hymenophyllaceae |
| *Trichomanes pyxidiferum* L. | Hymenophyllaceae |
| *Trichomanes radicans* Sw. | Hymenophyllaceae |

**Table S1**(continued)

| *Trichomanes rigidum Sw.* | Hymenophyllaceae |
| --- | --- |
| *Isoetes australis R.O. Williams* | Isoetaceae |
| *Craterostigma hirsutum S. Moore* | Linderniaceae |
| *Craterostigma lanceolatum (Engl.) Skan* | Linderniaceae |
| *Craterostigma plantagineum Hochst.* | Linderniaceae |
| *Craterostigma pumilum Hochst.* | Linderniaceae |
| *Craterostigma wilmsii Engl. ex Diels* | Linderniaceae |
| *Lindernia brevidens Skan* | Linderniaceae |
| *Lindernia intrepidus (Dinter) Oberm.* | Linderniaceae |
| *Lindernia monroi* (S. Moore) Eb. Fisch. | Linderniaceae |
| *Lindernia purpurea* (Lebrun & Touss.) R. Germ. | Linderniaceae |
| *Linderniella pulchella* (Skan) Eb. Fisch., Schäferh. & Kai Müll. | Linderniaceae |
| *Linderniella wilmsii* (Engl. ex Diels) Eb. Fisch., Schäferh. & Kai Müll. | Linderniaceae |
| *Myrothamnus flabellifolius* Welw. | Myrothamnaceae |
| *Myrothamnus moschatus* (Baill.) Baill. ex Nied. | Myrothamnaceae |
| *Eragrostiella bifaria* (Vahl) Bor | Poaceae |
| *Eragrostiella brachyphylla* (Stapf) Bor | Poaceae |
| *Eragrostiella nardoides* (Trin.) Bor | Poaceae |
| *Eragrostis nindensis* Ficalho & Hiern | Poaceae |
| *Eragrostis paradoxa* Launert | Poaceae |
| *Micrachne patentiflora* (Stent & J.M. Rattray) P.M. Peterson | Poaceae |
| *Micraira adamsii* Lazarides | Poaceae |
| *Micraira lazaridis* L.G. Clark, Wendel & Craven | Poaceae |
| *Micraira multinervia* Lazarides | Poaceae |
| *Micraira spinifera* Lazarides | Poaceae |
| *Micraira subulifolia* F. Muell. | Poaceae |
| *Micraira tenuis* Lazarides | Poaceae |
| *Micraira viscidula* Lazarides | Poaceae |
| *Microchloa caffra* Nees | Poaceae |
| *Microchloa indica* (L. f.) P. Beauv. | Poaceae |
| *Microchloa kunthii* Desv. | Poaceae |
| *Oropetium aristatum* (Stapf) Pilg. | Poaceae |
| *Oropetium capense* Stapf | Poaceae |
| *Oropetium roxburghianum* S.M. Phillips | Poaceae |
| *Oropetium thomaeum* (L. f.) Trin. | Poaceae |
| *Sporobolus atrovirens* (Kunth) Kunth | Poaceae |
| *Sporobolus elongatus* R. Br. | Poaceae |
| *Sporobolus festivus* Hochst. ex A. Rich. | Poaceae |
| *Sporobolus fimbriatus* (Trin.) Nees | Poaceae |
| *Sporobolus pellucidus* Hochst. | Poaceae |
| *Sporobolus ruspolianus* Chiov. | Poaceae |
| *Sporobolus stapfianus* Gand. | Poaceae |
| *Styppeiochloa hitchcockii* (A. Camus) Cope | Poaceae |

**Table S1**(continued)

| *Tripogon capillatus* Jaub. & Spach | Poaceae |
| --- | --- |
| *Tripogon curvatus* S.M. Phillips & Launert | Poaceae |
| *Tripogon filiformis* Nees | Poaceae |
| *Tripogon jacquemontii* Stapf | Poaceae |
| *Tripogon lisboae* Stapf | Poaceae |
| *Tripogon major* Hook. f. | Poaceae |
| *Tripogon polyanthus* Naik & Patunkar | Poaceae |
| *Tripogon curvatus* (F. Muell.) P.M. Peterson & Romasch. | Poaceae |
| *Tripogonella minima* (A. Rich.) P.M. Peterson & Romasch. | Poaceae |
| *Tripogonella spicata* (Nees) P.M. Peterson & Romasch. | Poaceae |
| *Ctenopteris heterophylla* Tindale | Polypodiaceae |
| *Goniophlebium furfuraceum* (Schltdl. & Cham.) T. Moore | Polypodiaceae |
| *Pleopeltis minima* (Bory) J. Prado & R.Y. Hirai | Polypodiaceae |
| *Pleopeltis plebeia* (Schltdl. & Cham.) A.R. Sm. & Tejero | Polypodiaceae |
| *Pleopeltis pleopeltifolia* (Raddi) Alston | Polypodiaceae |
| *Pleopeltis polypodioides* (L.) E.G. Andrews & Windham | Polypodiaceae |
| *Polypodium cambricum* L. | Polypodiaceae |
| *Polypodium interjectum* Shivas | Polypodiaceae |
| *Polypodium remotum* Desv. | Polypodiaceae |
| *Polypodium virginianum* L. | Polypodiaceae |
| *Polypodium vulgare* L. | Polypodiaceae |
| *Actiniopteris australis* (L. f.) Link | Pteridaceae |
| *Actiniopteris dimorpha* Pic. Serm. | Pteridaceae |
| *Actiniopteris radiata* (Sw.) Link | Pteridaceae |
| *Actiniopteris semiflabellata* Pic. Serm. | Pteridaceae |
| *Adiantum hispidulum* Sw. | Pteridaceae |
| *Adiantum incisum* Forssk. | Pteridaceae |
| *Adiantum latifolium* Lam. | Pteridaceae |
| *Adiantum raddianum* C. Presl | Pteridaceae |
| *Aleuritopteris albomarginata* (C.B. Clarke) Ching | Pteridaceae |
| *Aleuritopteris farinosa* (Forssk.) Fée | Pteridaceae |
| *Allosorus coriaceus* (Decne.) Christenh. | Pteridaceae |
| *Allosorus pteridioides* (Reichard) Christenh. | Pteridaceae |
| *Argyrochosma fendleri* (Kunze) Windham | Pteridaceae |
| *Astrolepis cochisensis* (Goodd.) D.M. Benham & Windham | Pteridaceae |
| *Astrolepis integerrima* (Hook.) D.M. Benham & Windham | Pteridaceae |
| *Astrolepis sinuata* (Lag. ex Sw.) D.M. Benham & Windham | Pteridaceae |
| *Bommeria hispida* (Mett. ex Kuhn) Underw. | Pteridaceae |
| *Cheilanthes bonariensis* (Willd.) Proctor | Pteridaceae |
| *Cheilanthes buchtienii* (Rosenst.) R.M. Tryon | Pteridaceae |
| *Cheilanthes capensis* (Thunb.) Sw. | Pteridaceae |
| *Cheilanthes catanensis* (Cosent.) H.P. Fuchs | Pteridaceae |
| *Cheilanthes depauperata* Baker | Pteridaceae |

**Table S1**(continued)

| *Cheilanthes dinteri* Brause | Pteridaceae |
| --- | --- |
| *Cheilanthes distans* (R. Br.) Mett. | Pteridaceae |
| *Cheilanthes eckloniana* Mett. | Pteridaceae |
| *Cheilanthes hirta* Sw. | Pteridaceae |
| *Cheilanthes inaequalis* (Kunze) Mett. | Pteridaceae |
| *Cheilanthes lasiophylla* Pic. Serm. | Pteridaceae |
| *Cheilanthes lendigera* (Cav.) Sw. | Pteridaceae |
| *Cheilanthes marginata* Kunth | Pteridaceae |
| *Cheilanthes marlothii* (Hieron.) Domin | Pteridaceae |
| *Cheilanthes multifida* (Sw.) Sw. | Pteridaceae |
| *Cheilanthes myriophylla* Desv. | Pteridaceae |
| *Cheilanthes nitidula* Wall. ex Hook. | Pteridaceae |
| *Cheilanthes notholaenoides* (Desv.) Maxon ex Weath. | Pteridaceae |
| *Cheilanthes parryi* (D.C. Eaton) Domin | Pteridaceae |
| *Cheilanthes parviloba* Sw. | Pteridaceae |
| *Cheilanthes pringlei* Davenp. | Pteridaceae |
| *Cheilanthes quadripinnata* (Forssk.) Kuhn | Pteridaceae |
| *Cheilanthes sieberi* Kunze | Pteridaceae |
| *Cheilanthes tenuifolia* (Burm. f.) Sw. | Pteridaceae |
| *Cheilanthes tomentosa* Link | Pteridaceae |
| *Cheilanthes viridis* (Forssk.) Sw. | Pteridaceae |
| *Cheilanthes wrightii* Hook. | Pteridaceae |
| *Cosentinia vellea* (Aiton) Tod. | Pteridaceae |
| *Doryopteris collina* (Raddi) J. Sm. | Pteridaceae |
| *Doryopteris concolor* (Langsd. & Fisch.) Kuhn | Pteridaceae |
| *Doryopteris kitchingii* (Baker) Bonap. | Pteridaceae |
| *Doryopteris pedata* (L.) Fée | Pteridaceae |
| *Doryopteris triphylla* (Lam.) Christ | Pteridaceae |
| *Doryopteris varians* (Raddi) Sm. | Pteridaceae |
| *Haplopteris volkensii* (Hieron.) E.H. Crane | Pteridaceae |
| *Hemionitis palmata* L. | Pteridaceae |
| *Hemionitis tomentosa* (Lam.) Raddi | Pteridaceae |
| *Myriopteris rufa* Fée | Pteridaceae |
| *Negripteris scioana* (Chiov.) Pic. Serm. | Pteridaceae |
| *Notholaena dipinnata* Fraser-Jenk. | Pteridaceae |
| *Notholaena lanuginosa* Desv. ex Poir. | Pteridaceae |
| *Notholaena muelleri* (Hook.) Fraser-Jenk. | Pteridaceae |
| *Onychium divaricatum* (Poir.) Alston | Pteridaceae |
| *Paragymnopteris marantae* (L.) K.H. Shing | Pteridaceae |
| *Pellaea andromedifolia* (Kaulf.) Fée | Pteridaceae |
| *Pellaea atropurpurea* (L.) Link | Pteridaceae |
| *Pellaea boivinii* Hook. | Pteridaceae |
| *Pellaea brachyptera* (T. Moore) Baker | Pteridaceae |

**Table S1**(continued)

| *Pellaea bridgesii Hook.* | Pteridaceae |
| --- | --- |
| *Pellaea calomelanos (Sw.) Link* | Pteridaceae |
| *Pellaea dura (Willd.) Hook.* | Pteridaceae |
| *Pellaea falcata* Fée | Pteridaceae |
| *Pellaea glabella* Mett. ex Kuhn | Pteridaceae |
| *Pellaea longipilosa* Bonap. | Pteridaceae |
| *Pellaea mucronata* (D.C. Eaton) D.C. Eaton | Pteridaceae |
| *Pellaea ovata* (Desv.) Weath. | Pteridaceae |
| *Pellaea pectiniformis* Baker | Pteridaceae |
| *Pellaea rotundifolia* (G. Forst.) Hook. | Pteridaceae |
| *Pellaea sagittata* (Cav.) Link | Pteridaceae |
| *Pellaea ternifolia* (Cav.) Link | Pteridaceae |
| *Pellaea truncata* Goodd. | Pteridaceae |
| *Pellaea wrightiana* Hook. | Pteridaceae |
| *Pentagramma triangularis* (Kaulf.) Yatsk., Windham & E. Wollenw. | Pteridaceae |
| *Vittaria guineensis* Desv. | Pteridaceae |
| *Vittaria isoetifolia* Bory | Pteridaceae |
| *Schizaea pusilla* Pursh | Schizaeaceae |
| *Selaginella arizonica* Maxon | Selaginellaceae |
| *Selaginella bryopteris* Baker | Selaginellaceae |
| *Selaginella caffrorum* (Milde) Hieron. | Selaginellaceae |
| *Selaginella convoluta* (Arn.) Spring | Selaginellaceae |
| *Selaginella densa* Rydb. | Selaginellaceae |
| *Selaginella digitata* Spring | Selaginellaceae |
| *Selaginella dregei* (C. Presl) Hieron. | Selaginellaceae |
| *Selaginella echinata* Baker | Selaginellaceae |
| *Selaginella eremophila* Maxon | Selaginellaceae |
| *Selaginella helicoclada* Alston | Selaginellaceae |
| *Selaginella helvetica* (L.) Spring | Selaginellaceae |
| *Selaginella imbricata* (Forssk.) Spring ex Decne. | Selaginellaceae |
| *Selaginella lepidophylla* (Hook. & Grev.) Spring | Selaginellaceae |
| *Selaginella nivea* Alston | Selaginellaceae |
| *Selaginella njamnjamensis* Hieron. | Selaginellaceae |
| *Selaginella peruviana* (Milde) Hieron. | Selaginellaceae |
| *Selaginella phillipsiana* (Hieron.) Alston | Selaginellaceae |
| *Selaginella pilifera* A. Braun | Selaginellaceae |
| *Selaginella rupincola* Underw. | Selaginellaceae |
| *Selaginella sartorii* Hieron. | Selaginellaceae |
| *Selaginella sellowii* Hieron. | Selaginellaceae |
| *Selaginella tamariscina* (P. Beauv.) Spring | Selaginellaceae |
| *Selaginella trisulcata* Aspl. | Selaginellaceae |
| *Selaginella yemensis* (Sw.) Spring | Selaginellaceae |
| *Arthropteris orientalis* (J.F. Gmel.) Posth. | Tectariaceae |

**Table S1**(continued)

| *Acanthochlamys bracteata P.C. Kao* | Velloziaceae |
| --- | --- |
| *Barbacenia blackii L.B. Sm.* | Velloziaceae |
| *Barbacenia fanniae (N.L. Menezes) Mello-Silva* | Velloziaceae |
| *Barbacenia flava Mart. ex Schult. f.* | Velloziaceae |
| *Barbacenia fragrans Goethart & Henrard* | Velloziaceae |
| *Barbacenia gentianoides Goethart & Henrard* | Velloziaceae |
| *Barbacenia gounelleana* Beauverd | Velloziaceae |
| *Barbacenia graminifolia* L.B. Sm. | Velloziaceae |
| *Barbacenia longiflora* Mart. | Velloziaceae |
| *Barbacenia longiscapa* Goethart & Henrard | Velloziaceae |
| *Barbacenia macrantha* Lem. | Velloziaceae |
| *Barbacenia purpurea* Hook. | Velloziaceae |
| *Barbacenia riedeliana* Goethart & Henrard | Velloziaceae |
| *Barbacenia seubertiana* Goethart & Henrard | Velloziaceae |
| *Barbacenia spectabilis* L.B. Sm. & Ayensu | Velloziaceae |
| *Barbacenia tomentosa* Mart. | Velloziaceae |
| *Barbaceniopsis boliviensis* (Baker) L.B. Sm. | Velloziaceae |
| *Barbaceniopsis humahuaquensis* Noher | Velloziaceae |
| *Vellozia albiflora* Pohl | Velloziaceae |
| *Vellozia andina* Ibisch, R. Vásquez & Nowicki | Velloziaceae |
| *Vellozia angustifolia* Goethart & Henrard | Velloziaceae |
| *Vellozia candida* J.C. Mikan | Velloziaceae |
| *Vellozia caput-ardeae* L.B. Sm. & Ayensu | Velloziaceae |
| *Vellozia caruncularis* Mart. ex Seub. | Velloziaceae |
| *Vellozia ciliata* L.B. Sm. | Velloziaceae |
| *Vellozia compacta* Mart. ex Schult. f. | Velloziaceae |
| *Vellozia declinans* Goethart & Henrard | Velloziaceae |
| *Vellozia epidendroides* Mart. ex Schult. f. | Velloziaceae |
| *Vellozia flavicans* Mart. ex Schult. f. | Velloziaceae |
| *Vellozia glochidea* Pohl | Velloziaceae |
| *Vellozia hatschbachii* L.B. Sm. & Ayensu | Velloziaceae |
| *Vellozia hirsuta* Goethart & Henrard | Velloziaceae |
| *Vellozia nanuzae* L.B. Sm. & Ayensu | Velloziaceae |
| *Vellozia nivea* L.B. Sm. & Ayensu | Velloziaceae |
| *Vellozia plicata* Mart. | Velloziaceae |
| *Vellozia pulchra* L.B. Sm. | Velloziaceae |
| *Vellozia resinosa* Mart. ex Schult. f. | Velloziaceae |
| *Vellozia sellowii* Seub. | Velloziaceae |
| *Vellozia semirii* Mello-Silva & N.L. Menezes | Velloziaceae |
| *Vellozia squalida* Mart. ex Schult. f. | Velloziaceae |
| *Vellozia streptophylla* L.B. Sm. | Velloziaceae |
| *Vellozia subscabra* J.C. Mikan | Velloziaceae |
| *Vellozia taxifolia* (Mart. ex Schult. f.) Mart. ex Seub. | Velloziaceae |

**Table S1**(continued)

| *Vellozia tubiflora (A. Rich.) Kunth* | Velloziaceae |
| --- | --- |
| *Vellozia variabilis Mart. ex Schult. f.* | Velloziaceae |
| *Vellozia variegata Goethart & Henrard* | Velloziaceae |
| *Vellozia verruculosa Mart. ex Schult. f.* | Velloziaceae |
| *Xerophyta dasylirioides Baker* | Velloziaceae |
| *Xerophyta eglandulosa H. Perrier* | Velloziaceae |
| *Xerophyta elegans (Balf.) Baker* | Velloziaceae |
| *Xerophyta equisetoides Baker* | Velloziaceae |
| *Xerophyta humilis (Baker) T. Durand & Schinz* | Velloziaceae |
| *Xerophyta nandrasanae* Phillipson & Lowry | Velloziaceae |
| *Xerophyta pectinata* Baker | Velloziaceae |
| *Xerophyta pinifolia* Lam. | Velloziaceae |
| *Xerophyta retinervis* Baker | Velloziaceae |
| *Xerophyta rippsteinii* L.B. Sm., J.-P. Lebrun & Stork | Velloziaceae |
| *Xerophyta scabrida* (Pax) T. Durand & Schinz | Velloziaceae |
| *Xerophyta schlechteri* (Baker) N.L. Menezes | Velloziaceae |
| *Xerophyta schnizleinia* (L.B. Sm. & Ayensu) Baker | Velloziaceae |
| *Xerophyta spekei* Baker | Velloziaceae |
| *Xerophyta splendens* (Rendle) N.L. Menezes | Velloziaceae |
| *Xerophyta squarrosa* Baker | Velloziaceae |
| *Xerophyta villosa* (Baker) L.B. Sm. & Ayensu | Velloziaceae |
| *Xerophyta viscosa* Baker | Velloziaceae |
| *Woodsia ilvensis* (L.) R. Br. | Woodsiaceae |

**Table S2**References to GBIF datasets used to estimate the species distribution.

| Species | References to used GBIF datasets |
| --- | --- |
| *Anemia ferruginea* | GBIF.org (22 March 2023) GBIF Occurrence Download https://doi.org/10.15468/dl.sa9zab |
| *Anemia flexuosa* | GBIF.org (22 March 2023) GBIF Occurrence Download https://doi.org/10.15468/dl.vde2cc |
| *Anemia mexicana* | GBIF.org (22 March 2023) GBIF Occurrence Download https://doi.org/10.15468/dl.k9mbkq |
| *Anemia rotundifolia* | GBIF.org (22 March 2023) GBIF Occurrence Download https://doi.org/10.15468/dl.epbqcq |
| *Anemia tomentosa* | GBIF.org (22 March 2023) GBIF Occurrence Download https://doi.org/10.15468/dl.3tmkpy |
| *Anemia villosa* | GBIF.org (22 March 2023) GBIF Occurrence Download https://doi.org/10.15468/dl.j43tqx |
| *Mohria caffrorum* | GBIF.org (22 March 2023) GBIF Occurrence Download https://doi.org/10.15468/dl.npnhhm |
| *Asplenium adiantum-nigrum* | GBIF.org (22 March 2023) GBIF Occurrence Download https://doi.org/10.15468/dl.7pppyw |
| *Asplenium aethiopicum* | GBIF.org (22 March 2023) GBIF Occurrence Download https://doi.org/10.15468/dl.hjqzcy |
| *Asplenium ceterach* | GBIF.org (22 March 2023) GBIF Occurrence Download https://doi.org/10.15468/dl.k7vbgr |
| *Asplenium cordatum* | GBIF.org (22 March 2023) GBIF Occurrence Download https://doi.org/10.15468/dl.v6deag |
| *Asplenium dalhousiae* | GBIF.org (22 March 2023) GBIF Occurrence Download https://doi.org/10.15468/dl.upva47 |
| *Asplenium friesiorum* | GBIF.org (22 March 2023) GBIF Occurrence Download https://doi.org/10.15468/dl.gen6n3 |
| *Asplenium megalura* | GBIF.org (22 March 2023) GBIF Occurrence Download https://doi.org/10.15468/dl.uweaw5 |
| *Asplenium monanthes* | GBIF.org (22 March 2023) GBIF Occurrence Download https://doi.org/10.15468/dl.6c9ztx |
| *Asplenium obovatum* | GBIF.org (22 March 2023) GBIF Occurrence Download https://doi.org/10.15468/dl.a9tgsp |
| *Asplenium praegracile* | GBIF.org (22 March 2023) GBIF Occurrence Download https://doi.org/10.15468/dl.vet62p |
| *Asplenium pringlei* | GBIF.org (22 March 2023) GBIF Occurrence Download https://doi.org/10.15468/dl.53g72r |
| *Asplenium ruta-muraria* | GBIF.org (22 March 2023) GBIF Occurrence Download https://doi.org/10.15468/dl.x2x8ej |
| *Asplenium rutifolium* | GBIF.org (22 March 2023) GBIF Occurrence Download https://doi.org/10.15468/dl.4e9y4h |

**Table S2**(continued)

| *Asplenium sandersonii* | GBIF.org (22 March 2023) GBIF Occurrence Download https://doi.org/10.15468/dl.k5hxv2 |
| --- | --- |
| *Asplenium septentrionale* | GBIF.org (22 March 2023) GBIF Occurrence Download https://doi.org/10.15468/dl.eseasq |
| *Asplenium theciferum* | GBIF.org (22 March 2023) GBIF Occurrence Download https://doi.org/10.15468/dl.5mdju7 |
| *Asplenium trichomanes* | GBIF.org (22 March 2023) GBIF Occurrence Download https://doi.org/10.15468/dl.5m7wdb |
| *Asplenium uhligii* | GBIF.org (22 March 2023) GBIF Occurrence Download https://doi.org/10.15468/dl.5u77u2 |
| *Pleurosorus rutifolius* | GBIF.org (22 March 2023) GBIF Occurrence Download https://doi.org/10.15468/dl.pfykca |
| *Borya constricta* | GBIF.org (22 March 2023) GBIF Occurrence Download https://doi.org/10.15468/dl.azjcre |
| *Borya inopinata* | GBIF.org (22 March 2023) GBIF Occurrence Download https://doi.org/10.15468/dl.5vyu6h |
| *Borya mirabilis* | GBIF.org (22 March 2023) GBIF Occurrence Download https://doi.org/10.15468/dl.4u7yvh |
| *Borya nitida* | GBIF.org (22 March 2023) GBIF Occurrence Download https://doi.org/10.15468/dl.6f4hvs |
| *Borya scirpoidea* | GBIF.org (22 March 2023) GBIF Occurrence Download https://doi.org/10.15468/dl.mc6zqm |
| *Borya septentrionalis* | GBIF.org (22 March 2023) GBIF Occurrence Download https://doi.org/10.15468/dl.pznhwz |
| *Borya sphaerocephala* | GBIF.org (22 March 2023) GBIF Occurrence Download https://doi.org/10.15468/dl.4hfbhc |
| *Pitcairnia lanuginosa* | GBIF.org (22 March 2023) GBIF Occurrence Download https://doi.org/10.15468/dl.p4szxw |
| *Blossfeldia liliputana* | GBIF.org (22 March 2023) GBIF Occurrence Download https://doi.org/10.15468/dl.69baj9 |
| *Afrotrilepis pilosa* | GBIF.org (22 March 2023) GBIF Occurrence Download https://doi.org/10.15468/dl.n52wrd |
| *Coleochloa abyssinica* | GBIF.org (22 March 2023) GBIF Occurrence Download https://doi.org/10.15468/dl.japukw |
| *Coleochloa microcephala* | GBIF.org (22 March 2023) GBIF Occurrence Download https://doi.org/10.15468/dl.jycdrx |
| *Coleochloa pallidior* | GBIF.org (22 March 2023) GBIF Occurrence Download https://doi.org/10.15468/dl.ps6mdw |
| *Coleochloa setifera* | GBIF.org (22 March 2023) GBIF Occurrence Download https://doi.org/10.15468/dl.jxfrsy |

**Table S2**(continued)

| *Microdracoides squamosus* | GBIF.org (22 March 2023) GBIF Occurrence Download https://doi.org/10.15468/dl.6pwvgd |
| --- | --- |
| *Trilepis ciliatifolia* | GBIF.org (22 March 2023) GBIF Occurrence Download https://doi.org/10.15468/dl.td7jpp |
| *Trilepis lhotzkiana* | GBIF.org (22 March 2023) GBIF Occurrence Download https://doi.org/10.15468/dl.w5mmwh |
| *Trilepis microstachya* | GBIF.org (22 March 2023) GBIF Occurrence Download https://doi.org/10.15468/dl.x6cyth |
| *Davallia angustata* | GBIF.org (22 March 2023) GBIF Occurrence Download https://doi.org/10.15468/dl.2298zs |
| *Elaphoglossum acrostichoides* | GBIF.org (22 March 2023) GBIF Occurrence Download https://doi.org/10.15468/dl.698qkw |
| *Elaphoglossum petiolatum* | GBIF.org (22 March 2023) GBIF Occurrence Download https://doi.org/10.15468/dl.8q88sf |
| *Elaphoglossum piloselloides* | GBIF.org (22 March 2023) GBIF Occurrence Download https://doi.org/10.15468/dl.z282q4 |
| *Boea hygrometrica* | GBIF.org (22 March 2023) GBIF Occurrence Download https://doi.org/10.15468/dl.uedr97 |
| *Boea hygroscopica* | GBIF.org (22 March 2023) GBIF Occurrence Download https://doi.org/10.15468/dl.4smyuc |
| *Damrongia clarkeana* | GBIF.org (22 March 2023) GBIF Occurrence Download https://doi.org/10.15468/dl.3wg7vn |
| *Haberlea rhodopensis* | GBIF.org (22 March 2023) GBIF Occurrence Download https://doi.org/10.15468/dl.tsuzdp |
| *Oreocharis mileensis* | GBIF.org (22 March 2023) GBIF Occurrence Download https://doi.org/10.15468/dl.euea3q |
| *Paraboea crassifolia* | GBIF.org (22 March 2023) GBIF Occurrence Download https://doi.org/10.15468/dl.qp5hf8 |
| *Paraboea rufescens* | GBIF.org (22 March 2023) GBIF Occurrence Download https://doi.org/10.15468/dl.fgf6s4 |
| *Ramonda myconi* | GBIF.org (22 March 2023) GBIF Occurrence Download https://doi.org/10.15468/dl.petm8e |
| *Ramonda nathaliae* | GBIF.org (22 March 2023) GBIF Occurrence Download https://doi.org/10.15468/dl.7aeqcg |
| *Ramonda serbica* | GBIF.org (22 March 2023) GBIF Occurrence Download https://doi.org/10.15468/dl.fw7ngm |
| *Cardiomanes reniforme* | GBIF.org (22 March 2023) GBIF Occurrence Download https://doi.org/10.15468/dl.27wr9f |
| *Crepidomanes chevalieri* | GBIF.org (22 March 2023) GBIF Occurrence Download https://doi.org/10.15468/dl.we3am5 |
| *Crepidomanes frappieri* | GBIF.org (22 March 2023) GBIF Occurrence Download https://doi.org/10.15468/dl.2m7evw |

**Table S2**(continued)

| *Crepidomanes inopinatum* | GBIF.org (22 March 2023) GBIF Occurrence Download https://doi.org/10.15468/dl.ahznv9 |
| --- | --- |
| *Crepidomanes melanotrichum* | GBIF.org (22 March 2023) GBIF Occurrence Download https://doi.org/10.15468/dl.r2sdq5 |
| *Didymoglossum erosum* | GBIF.org (22 March 2023) GBIF Occurrence Download https://doi.org/10.15468/dl.9avcdb |
| *Hymenoglossum cruentum* | GBIF.org (22 March 2023) GBIF Occurrence Download https://doi.org/10.15468/dl.thmyfs |
| *Hymenophyllum capillare* | GBIF.org (22 March 2023) GBIF Occurrence Download https://doi.org/10.15468/dl.wqpewp |
| *Hymenophyllum caudiculatum* | GBIF.org (22 March 2023) GBIF Occurrence Download https://doi.org/10.15468/dl.tekt8z |
| *Hymenophyllum dentatum* | GBIF.org (22 March 2023) GBIF Occurrence Download https://doi.org/10.15468/dl.98f432 |
| *Hymenophyllum fucoides* | GBIF.org (22 March 2023) GBIF Occurrence Download https://doi.org/10.15468/dl.vqx8mx |
| *Hymenophyllum hirsutum* | GBIF.org (22 March 2023) GBIF Occurrence Download https://doi.org/10.15468/dl.rr3cxv |
| *Hymenophyllum kuhnii* | GBIF.org (22 March 2023) GBIF Occurrence Download https://doi.org/10.15468/dl.f529nm |
| *Hymenophyllum peltatum* | GBIF.org (22 March 2023) GBIF Occurrence Download https://doi.org/10.15468/dl.6vh9hg |
| *Hymenophyllum plicatum* | GBIF.org (22 March 2023) GBIF Occurrence Download https://doi.org/10.15468/dl.bq2wu8 |
| *Hymenophyllum polyanthos* | GBIF.org (22 March 2023) GBIF Occurrence Download https://doi.org/10.15468/dl.qywftz |
| *Hymenophyllum sanguinolentum* | GBIF.org (22 March 2023) GBIF Occurrence Download https://doi.org/10.15468/dl.yeta3s |
| *Hymenophyllum splendidum* | GBIF.org (22 March 2023) GBIF Occurrence Download https://doi.org/10.15468/dl.7j4fhq |
| *Hymenophyllum tunbrigense* | GBIF.org (22 March 2023) GBIF Occurrence Download https://doi.org/10.15468/dl.z9k99v |
| *Polyphlebium borbonicum* | GBIF.org (22 March 2023) GBIF Occurrence Download https://doi.org/10.15468/dl.zvdfg9 |
| *Trichomanes bucinatum* | GBIF.org (22 March 2023) GBIF Occurrence Download https://doi.org/10.15468/dl.8r8yub |
| *Trichomanes capillaceum* | GBIF.org (22 March 2023) GBIF Occurrence Download https://doi.org/10.15468/dl.tumgu7 |
| *Trichomanes diaphanum* | GBIF.org (22 March 2023) GBIF Occurrence Download https://doi.org/10.15468/dl.qz2n6y |
| *Trichomanes polypodioides* | GBIF.org (22 March 2023) GBIF Occurrence Download https://doi.org/10.15468/dl.s4h6jx |

**Table S2**(continued)

| *Trichomanes pyxidiferum* | GBIF.org (22 March 2023) GBIF Occurrence Download https://doi.org/10.15468/dl.hgn9q4 |
| --- | --- |
| *Trichomanes radicans* | GBIF.org (22 March 2023) GBIF Occurrence Download https://doi.org/10.15468/dl.28ycp2 |
| *Trichomanes rigidum* | GBIF.org (22 March 2023) GBIF Occurrence Download https://doi.org/10.15468/dl.6y7be5 |
| *Isoetes australis* | GBIF.org (22 March 2023) GBIF Occurrence Download https://doi.org/10.15468/dl.a3hgeg |
| *Craterostigma hirsutum* | GBIF.org (22 March 2023) GBIF Occurrence Download https://doi.org/10.15468/dl.cmwwns |
| *Craterostigma lanceolatum* | GBIF.org (22 March 2023) GBIF Occurrence Download https://doi.org/10.15468/dl.k8qdxz |
| *Craterostigma plantagineum* | GBIF.org (22 March 2023) GBIF Occurrence Download https://doi.org/10.15468/dl.pnxv3a |
| *Craterostigma pumilum* | GBIF.org (22 March 2023) GBIF Occurrence Download https://doi.org/10.15468/dl.5hhk74 |
| *Craterostigma wilmsii* | GBIF.org (22 March 2023) GBIF Occurrence Download https://doi.org/10.15468/dl.26h4xy |
| *Lindernia brevidens* | GBIF.org (22 March 2023) GBIF Occurrence Download https://doi.org/10.15468/dl.3grqxn |
| *Lindernia intrepidus* | GBIF.org (22 March 2023) GBIF Occurrence Download https://doi.org/10.15468/dl.7bamba |
| *Lindernia monroi* | GBIF.org (22 March 2023) GBIF Occurrence Download https://doi.org/10.15468/dl.hmpnq7 |
| *Lindernia purpurea* | GBIF.org (22 March 2023) GBIF Occurrence Download https://doi.org/10.15468/dl.93cpxk |
| *Linderniella pulchella* | GBIF.org (22 March 2023) GBIF Occurrence Download https://doi.org/10.15468/dl.fns8vw |
| *Linderniella wilmsii* | GBIF.org (22 March 2023) GBIF Occurrence Download https://doi.org/10.15468/dl.et4dwe |
| *Myrothamnus flabellifolius* | GBIF.org (22 March 2023) GBIF Occurrence Download https://doi.org/10.15468/dl.cndpau |
| *Myrothamnus moschatus* | GBIF.org (22 March 2023) GBIF Occurrence Download https://doi.org/10.15468/dl.rjsf4g |
| *Eragrostiella bifaria* | GBIF.org (22 March 2023) GBIF Occurrence Download https://doi.org/10.15468/dl.gdhn3z |
| *Eragrostiella brachyphylla* | GBIF.org (22 March 2023) GBIF Occurrence Download https://doi.org/10.15468/dl.2du7jn |
| *Eragrostiella nardoides* | GBIF.org (22 March 2023) GBIF Occurrence Download https://doi.org/10.15468/dl.xqes2y |
| *Eragrostis nindensis* | GBIF.org (22 March 2023) GBIF Occurrence Download https://doi.org/10.15468/dl.t6ppej |

**Table S2**(continued)

| *Eragrostis paradoxa* | GBIF.org (22 March 2023) GBIF Occurrence Download https://doi.org/10.15468/dl.ehea4s |
| --- | --- |
| *Micrachne patentiflora* | GBIF.org (22 March 2023) GBIF Occurrence Download https://doi.org/10.15468/dl.cgs4jd |
| *Micraira adamsii* | GBIF.org (22 March 2023) GBIF Occurrence Download https://doi.org/10.15468/dl.9gwnej |
| *Micraira lazaridis* | GBIF.org (22 March 2023) GBIF Occurrence Download https://doi.org/10.15468/dl.bfzt9p |
| *Micraira multinervia* | GBIF.org (22 March 2023) GBIF Occurrence Download https://doi.org/10.15468/dl.phbv6k |
| *Micraira spinifera* | GBIF.org (22 March 2023) GBIF Occurrence Download https://doi.org/10.15468/dl.d6jhr9 |
| *Micraira subulifolia* | GBIF.org (22 March 2023) GBIF Occurrence Download https://doi.org/10.15468/dl.jda3g5 |
| *Micraira tenuis* | GBIF.org (22 March 2023) GBIF Occurrence Download https://doi.org/10.15468/dl.fy5f76 |
| *Micraira viscidula* | GBIF.org (22 March 2023) GBIF Occurrence Download https://doi.org/10.15468/dl.4qkycx |
| *Microchloa caffra* | GBIF.org (22 March 2023) GBIF Occurrence Download https://doi.org/10.15468/dl.uu9ttf |
| *Microchloa indica* | GBIF.org (22 March 2023) GBIF Occurrence Download https://doi.org/10.15468/dl.4pek7h |
| *Microchloa kunthii* | GBIF.org (22 March 2023) GBIF Occurrence Download https://doi.org/10.15468/dl.sym7kq |
| *Oropetium aristatum* | GBIF.org (22 March 2023) GBIF Occurrence Download https://doi.org/10.15468/dl.uuvqb7 |
| *Oropetium capense* | GBIF.org (22 March 2023) GBIF Occurrence Download https://doi.org/10.15468/dl.r6247a |
| *Oropetium roxburghianum* | GBIF.org (22 March 2023) GBIF Occurrence Download https://doi.org/10.15468/dl.z8m9zb |
| *Oropetium thomaeum* | GBIF.org (22 March 2023) GBIF Occurrence Download https://doi.org/10.15468/dl.x6jrdk |
| *Sporobolus atrovirens* | GBIF.org (22 March 2023) GBIF Occurrence Download https://doi.org/10.15468/dl.4r6ekh |
| *Sporobolus elongatus* | GBIF.org (22 March 2023) GBIF Occurrence Download https://doi.org/10.15468/dl.jb8ken |
| *Sporobolus festivus* | GBIF.org (22 March 2023) GBIF Occurrence Download https://doi.org/10.15468/dl.7k7ff4 |
| *Sporobolus fimbriatus* | GBIF.org (22 March 2023) GBIF Occurrence Download https://doi.org/10.15468/dl.ba8gp7 |
| *Sporobolus pellucidus* | GBIF.org (22 March 2023) GBIF Occurrence Download https://doi.org/10.15468/dl.7khsy2 |

**Table S2**(continued)

| *Sporobolus ruspolianus* | GBIF.org (22 March 2023) GBIF Occurrence Download https://doi.org/10.15468/dl.6mf4vv |
| --- | --- |
| *Sporobolus stapfianus* | GBIF.org (22 March 2023) GBIF Occurrence Download https://doi.org/10.15468/dl.nx4pse |
| *Styppeiochloa hitchcockii* | GBIF.org (22 March 2023) GBIF Occurrence Download https://doi.org/10.15468/dl.ncvmhc |
| *Tripogon capillatus* | GBIF.org (22 March 2023) GBIF Occurrence Download https://doi.org/10.15468/dl.kr7gbe |
| *Tripogon curvatus* | GBIF.org (22 March 2023) GBIF Occurrence Download https://doi.org/10.15468/dl.cke9zn |
| *Tripogon filiformis* | GBIF.org (22 March 2023) GBIF Occurrence Download https://doi.org/10.15468/dl.xk46dh |
| *Tripogon jacquemontii* | GBIF.org (22 March 2023) GBIF Occurrence Download https://doi.org/10.15468/dl.bf6pxj |
| *Tripogon lisboae* | GBIF.org (22 March 2023) GBIF Occurrence Download https://doi.org/10.15468/dl.xw9qxx |
| *Tripogon major* | GBIF.org (22 March 2023) GBIF Occurrence Download https://doi.org/10.15468/dl.pmnuqm |
| *Tripogon polyanthus* | GBIF.org (22 March 2023) GBIF Occurrence Download https://doi.org/10.15468/dl.eqmadm |
| *Tripogonella loliiformis* | GBIF.org (22 March 2023) GBIF Occurrence Download https://doi.org/10.15468/dl.gvfkbd |
| *Tripogonella minima* | GBIF.org (22 March 2023) GBIF Occurrence Download https://doi.org/10.15468/dl.gzfvm3 |
| *Tripogonella spicata* | GBIF.org (22 March 2023) GBIF Occurrence Download https://doi.org/10.15468/dl.cyn2mv |
| *Ctenopteris heterophylla* | GBIF.org (22 March 2023) GBIF Occurrence Download https://doi.org/10.15468/dl.zejbzk |
| *Goniophlebium furfuraceum* | GBIF.org (22 March 2023) GBIF Occurrence Download https://doi.org/10.15468/dl.2w7hjj |
| *Loxogramme abyssinica* | GBIF.org (22 March 2023) GBIF Occurrence Download https://doi.org/10.15468/dl.tdve7b |
| *Loxogramme lanceolata* | GBIF.org (22 March 2023) GBIF Occurrence Download https://doi.org/10.15468/dl.d7bq8c |
| *Melpomene flabelliformis* | GBIF.org (22 March 2023) GBIF Occurrence Download https://doi.org/10.15468/dl.82wkxt |
| *Melpomene peruviana* | GBIF.org (22 March 2023) GBIF Occurrence Download https://doi.org/10.15468/dl.bswgb4 |
| *Microgramma piloselloides* | GBIF.org (22 March 2023) GBIF Occurrence Download https://doi.org/10.15468/dl.xuwydq |
| *Pecluma eurybasis* | GBIF.org (22 March 2023) GBIF Occurrence Download https://doi.org/10.15468/dl.4pjhjr |

**Table S2**(continued)

| *Platycerium stemaria* | GBIF.org (22 March 2023) GBIF Occurrence Download https://doi.org/10.15468/dl.ffzdam |
| --- | --- |
| *Pleopeltis angusta* | GBIF.org (22 March 2023) GBIF Occurrence Download https://doi.org/10.15468/dl.4ms43y |
| *Pleopeltis crassinervata* | GBIF.org (22 March 2023) GBIF Occurrence Download https://doi.org/10.15468/dl.kvxbuh |
| *Pleopeltis hirsutissima* | GBIF.org (22 March 2023) GBIF Occurrence Download https://doi.org/10.15468/dl.grv3hs |
| *Pleopeltis macrocarpa* | GBIF.org (22 March 2023) GBIF Occurrence Download https://doi.org/10.15468/dl.ekk9fu |
| *Pleopeltis mexicana* | GBIF.org (22 March 2023) GBIF Occurrence Download https://doi.org/10.15468/dl.xfep7a |
| *Pleopeltis minima* | GBIF.org (22 March 2023) GBIF Occurrence Download https://doi.org/10.15468/dl.avrds9 |
| *Pleopeltis plebeia* | GBIF.org (22 March 2023) GBIF Occurrence Download https://doi.org/10.15468/dl.6q76fn |
| *Pleopeltis pleopeltifolia* | GBIF.org (22 March 2023) GBIF Occurrence Download https://doi.org/10.15468/dl.9aygvn |
| *Pleopeltis polypodioides* | GBIF.org (22 March 2023) GBIF Occurrence Download https://doi.org/10.15468/dl.x8kzf4 |
| *Polypodium cambricum* | GBIF.org (22 March 2023) GBIF Occurrence Download https://doi.org/10.15468/dl.6hmwga |
| *Polypodium interjectum* | GBIF.org (22 March 2023) GBIF Occurrence Download https://doi.org/10.15468/dl.3eyvcr |
| *Polypodium remotum* | GBIF.org (22 March 2023) GBIF Occurrence Download https://doi.org/10.15468/dl.dgvmch |
| *Polypodium virginianum* | GBIF.org (22 March 2023) GBIF Occurrence Download https://doi.org/10.15468/dl.v7bte4 |
| *Polypodium vulgare* | GBIF.org (22 March 2023) GBIF Occurrence Download https://doi.org/10.15468/dl.d6sc3j |
| *Actiniopteris australis* | GBIF.org (22 March 2023) GBIF Occurrence Download https://doi.org/10.15468/dl.dvctg3 |
| *Actiniopteris dimorpha* | GBIF.org (22 March 2023) GBIF Occurrence Download https://doi.org/10.15468/dl.ku38h4 |
| *Actiniopteris radiata* | GBIF.org (22 March 2023) GBIF Occurrence Download https://doi.org/10.15468/dl.w7xj3c |
| *Actiniopteris semiflabellata* | GBIF.org (22 March 2023) GBIF Occurrence Download https://doi.org/10.15468/dl.u46p8v |
| *Adiantum hispidulum* | GBIF.org (22 March 2023) GBIF Occurrence Download https://doi.org/10.15468/dl.mpnexj |
| *Adiantum incisum* | GBIF.org (22 March 2023) GBIF Occurrence Download https://doi.org/10.15468/dl.m4pmmc |

**Table S2**(continued)

| *Adiantum latifolium* | GBIF.org (22 March 2023) GBIF Occurrence Download https://doi.org/10.15468/dl.e5nqq9 |
| --- | --- |
| *Adiantum raddianum* | GBIF.org (22 March 2023) GBIF Occurrence Download https://doi.org/10.15468/dl.y96d4u |
| *Aleuritopteris albomarginata* | GBIF.org (22 March 2023) GBIF Occurrence Download https://doi.org/10.15468/dl.q2brh4 |
| *Aleuritopteris farinosa* | GBIF.org (22 March 2023) GBIF Occurrence Download https://doi.org/10.15468/dl.vzzhtk |
| *Allosorus coriaceus* | GBIF.org (22 March 2023) GBIF Occurrence Download https://doi.org/10.15468/dl.5sremd |
| *Allosorus pteridioides* | GBIF.org (22 March 2023) GBIF Occurrence Download https://doi.org/10.15468/dl.2j4ya8 |
| *Argyrochosma fendleri* | GBIF.org (22 March 2023) GBIF Occurrence Download https://doi.org/10.15468/dl.nysvzz |
| *Astrolepis cochisensis* | GBIF.org (22 March 2023) GBIF Occurrence Download https://doi.org/10.15468/dl.tewn6z |
| *Astrolepis integerrima* | GBIF.org (22 March 2023) GBIF Occurrence Download https://doi.org/10.15468/dl.9uawx5 |
| *Astrolepis sinuata* | GBIF.org (22 March 2023) GBIF Occurrence Download https://doi.org/10.15468/dl.ts2vtx |
| *Bommeria hispida* | GBIF.org (22 March 2023) GBIF Occurrence Download https://doi.org/10.15468/dl.php9yn |
| *Cheilanthes bonariensis* | GBIF.org (22 March 2023) GBIF Occurrence Download https://doi.org/10.15468/dl.33k6bd |
| *Cheilanthes buchtienii* | GBIF.org (22 March 2023) GBIF Occurrence Download https://doi.org/10.15468/dl.ktn4qp |
| *Cheilanthes capensis* | GBIF.org (22 March 2023) GBIF Occurrence Download https://doi.org/10.15468/dl.becmgd |
| *Cheilanthes catanensis* | GBIF.org (22 March 2023) GBIF Occurrence Download https://doi.org/10.15468/dl.dafh4z |
| *Cheilanthes depauperata* | GBIF.org (22 March 2023) GBIF Occurrence Download https://doi.org/10.15468/dl.4z7rsv |
| *Cheilanthes dinteri* | GBIF.org (22 March 2023) GBIF Occurrence Download https://doi.org/10.15468/dl.j7e6tv |
| *Cheilanthes distans* | GBIF.org (22 March 2023) GBIF Occurrence Download https://doi.org/10.15468/dl.nnvwym |
| *Cheilanthes eckloniana* | GBIF.org (22 March 2023) GBIF Occurrence Download https://doi.org/10.15468/dl.d3ytby |
| *Cheilanthes fragillima* | GBIF.org (22 March 2023) GBIF Occurrence Download https://doi.org/10.15468/dl.c9v9rh |
| *Cheilanthes glauca* | GBIF.org (22 March 2023) GBIF Occurrence Download https://doi.org/10.15468/dl.5quw65 |

**Table S2**(continued)

| *Cheilanthes gracillima* | GBIF.org (22 March 2023) GBIF Occurrence Download https://doi.org/10.15468/dl.usveyj |
| --- | --- |
| *Cheilanthes hirta* | GBIF.org (22 March 2023) GBIF Occurrence Download https://doi.org/10.15468/dl.d59eff |
| *Cheilanthes inaequalis* | GBIF.org (22 March 2023) GBIF Occurrence Download https://doi.org/10.15468/dl.xba4pw |
| *Cheilanthes lasiophylla* | GBIF.org (22 March 2023) GBIF Occurrence Download https://doi.org/10.15468/dl.m9rq8d |
| *Cheilanthes lendigera* | GBIF.org (22 March 2023) GBIF Occurrence Download https://doi.org/10.15468/dl.uvqmw6 |
| *Cheilanthes marginata* | GBIF.org (22 March 2023) GBIF Occurrence Download https://doi.org/10.15468/dl.hxccfx |
| *Cheilanthes marlothii* | GBIF.org (22 March 2023) GBIF Occurrence Download https://doi.org/10.15468/dl.ke2gxv |
| *Cheilanthes multifida* | GBIF.org (22 March 2023) GBIF Occurrence Download https://doi.org/10.15468/dl.9csppx |
| *Cheilanthes myriophylla* | GBIF.org (22 March 2023) GBIF Occurrence Download https://doi.org/10.15468/dl.93r2t5 |
| *Cheilanthes nitidula* | GBIF.org (22 March 2023) GBIF Occurrence Download https://doi.org/10.15468/dl.auv9qf |
| *Cheilanthes notholaenoides* | GBIF.org (22 March 2023) GBIF Occurrence Download https://doi.org/10.15468/dl.k6jspn |
| *Cheilanthes parryi* | GBIF.org (22 March 2023) GBIF Occurrence Download https://doi.org/10.15468/dl.xzt4gf |
| *Cheilanthes parviloba* | GBIF.org (22 March 2023) GBIF Occurrence Download https://doi.org/10.15468/dl.kaxfbp |
| *Cheilanthes pringlei* | GBIF.org (22 March 2023) GBIF Occurrence Download https://doi.org/10.15468/dl.evnxrg |
| *Cheilanthes quadripinnata* | GBIF.org (22 March 2023) GBIF Occurrence Download https://doi.org/10.15468/dl.m8jywm |
| *Cheilanthes sieberi* | GBIF.org (22 March 2023) GBIF Occurrence Download https://doi.org/10.15468/dl.qn5f4q |
| *Cheilanthes tenuifolia* | GBIF.org (22 March 2023) GBIF Occurrence Download https://doi.org/10.15468/dl.62mnf6 |
| *Cheilanthes tomentosa* | GBIF.org (22 March 2023) GBIF Occurrence Download https://doi.org/10.15468/dl.snkpaq |
| *Cheilanthes viridis* | GBIF.org (22 March 2023) GBIF Occurrence Download https://doi.org/10.15468/dl.76qb8a |
| *Cheilanthes wrightii* | GBIF.org (22 March 2023) GBIF Occurrence Download https://doi.org/10.15468/dl.nxy7m6 |
| *Cosentinia vellea* | GBIF.org (22 March 2023) GBIF Occurrence Download https://doi.org/10.15468/dl.dafh4z |

**Table S2**(continued)

| *Doryopteris collina* | GBIF.org (22 March 2023) GBIF Occurrence Download https://doi.org/10.15468/dl.mpefs7 |
| --- | --- |
| *Doryopteris concolor* | GBIF.org (22 March 2023) GBIF Occurrence Download https://doi.org/10.15468/dl.g6dnze |
| *Doryopteris kitchingii* | GBIF.org (22 March 2023) GBIF Occurrence Download https://doi.org/10.15468/dl.jnuegy |
| *Doryopteris pedata* | GBIF.org (22 March 2023) GBIF Occurrence Download https://doi.org/10.15468/dl.2bac3d |
| *Doryopteris triphylla* | GBIF.org (22 March 2023) GBIF Occurrence Download https://doi.org/10.15468/dl.4xrf5t |
| *Doryopteris varians* | GBIF.org (22 March 2023) GBIF Occurrence Download https://doi.org/10.15468/dl.dx9hp4 |
| *Haplopteris volkensii* | GBIF.org (22 March 2023) GBIF Occurrence Download https://doi.org/10.15468/dl.sp37at |
| *Hemionitis palmata* | GBIF.org (22 March 2023) GBIF Occurrence Download https://doi.org/10.15468/dl.c66s7c |
| *Hemionitis tomentosa* | GBIF.org (22 March 2023) GBIF Occurrence Download https://doi.org/10.15468/dl.6nrsvt |
| *Myriopteris rufa* | GBIF.org (22 March 2023) GBIF Occurrence Download https://doi.org/10.15468/dl.xrecnp |
| *Negripteris scioana* | GBIF.org (22 March 2023) GBIF Occurrence Download https://doi.org/10.15468/dl.hv5594 |
| *Notholaena dipinnata* | GBIF.org (22 March 2023) GBIF Occurrence Download https://doi.org/10.15468/dl.n63ua8 |
| *Notholaena lanuginosa* | GBIF.org (22 March 2023) GBIF Occurrence Download https://doi.org/10.15468/dl.dafh4z |
| *Notholaena muelleri* | GBIF.org (22 March 2023) GBIF Occurrence Download https://doi.org/10.15468/dl.thgc6z |
| *Onychium divaricatum* | GBIF.org (22 March 2023) GBIF Occurrence Download https://doi.org/10.15468/dl.y35mxa |
| *Paragymnopteris marantae* | GBIF.org (22 March 2023) GBIF Occurrence Download https://doi.org/10.15468/dl.tgqkfw |
| *Pellaea andromedifolia* | GBIF.org (22 March 2023) GBIF Occurrence Download https://doi.org/10.15468/dl.brw9km |
| *Pellaea atropurpurea* | GBIF.org (22 March 2023) GBIF Occurrence Download https://doi.org/10.15468/dl.r4kujm |
| *Pellaea boivinii* | GBIF.org (22 March 2023) GBIF Occurrence Download https://doi.org/10.15468/dl.a7as3z |
| *Pellaea brachyptera* | GBIF.org (22 March 2023) GBIF Occurrence Download https://doi.org/10.15468/dl.rr7h9b |
| *Pellaea bridgesii* | GBIF.org (22 March 2023) GBIF Occurrence Download https://doi.org/10.15468/dl.t9bqts |

**Table S2**(continued)

| *Pellaea calomelanos* | GBIF.org (22 March 2023) GBIF Occurrence Download https://doi.org/10.15468/dl.nnpkuu |
| --- | --- |
| *Pellaea dura* | GBIF.org (22 March 2023) GBIF Occurrence Download https://doi.org/10.15468/dl.zsgpw2 |
| *Pellaea falcata* | GBIF.org (22 March 2023) GBIF Occurrence Download https://doi.org/10.15468/dl.v34edm |
| *Pellaea glabella* | GBIF.org (22 March 2023) GBIF Occurrence Download https://doi.org/10.15468/dl.6k7jc6 |
| *Pellaea longipilosa* | GBIF.org (22 March 2023) GBIF Occurrence Download https://doi.org/10.15468/dl.h38cvb |
| *Pellaea mucronata* | GBIF.org (22 March 2023) GBIF Occurrence Download https://doi.org/10.15468/dl.7yjd8k |
| *Pellaea ovata* | GBIF.org (22 March 2023) GBIF Occurrence Download https://doi.org/10.15468/dl.vdajbu |
| *Pellaea pectiniformis* | GBIF.org (22 March 2023) GBIF Occurrence Download https://doi.org/10.15468/dl.7xkaek |
| *Pellaea rotundifolia* | GBIF.org (22 March 2023) GBIF Occurrence Download https://doi.org/10.15468/dl.8hy8j9 |
| *Pellaea sagittata* | GBIF.org (22 March 2023) GBIF Occurrence Download https://doi.org/10.15468/dl.bx8thv |
| *Pellaea ternifolia* | GBIF.org (22 March 2023) GBIF Occurrence Download https://doi.org/10.15468/dl.cbmtcn |
| *Pellaea truncata* | GBIF.org (22 March 2023) GBIF Occurrence Download https://doi.org/10.15468/dl.dn3w7t |
| *Pellaea wrightiana* | GBIF.org (22 March 2023) GBIF Occurrence Download https://doi.org/10.15468/dl.9rd92u |
| *Pentagramma triangularis* | GBIF.org (22 March 2023) GBIF Occurrence Download https://doi.org/10.15468/dl.qk7txw |
| *Vittaria guineensis* | GBIF.org (22 March 2023) GBIF Occurrence Download https://doi.org/10.15468/dl.8ng3dd |
| *Vittaria isoetifolia* | GBIF.org (22 March 2023) GBIF Occurrence Download https://doi.org/10.15468/dl.ynkgvh |
| *Schizaea pusilla* | GBIF.org (22 March 2023) GBIF Occurrence Download https://doi.org/10.15468/dl.4hhv6k |
| *Selaginella arizonica* | GBIF.org (22 March 2023) GBIF Occurrence Download https://doi.org/10.15468/dl.sgt2tu |
| *Selaginella bryopteris* | GBIF.org (22 March 2023) GBIF Occurrence Download https://doi.org/10.15468/dl.gkjxpw |
| *Selaginella caffrorum* | GBIF.org (22 March 2023) GBIF Occurrence Download https://doi.org/10.15468/dl.frchjk |
| *Selaginella convoluta* | GBIF.org (22 March 2023) GBIF Occurrence Download https://doi.org/10.15468/dl.xd6ake |

**Table S2**(continued)

| *Selaginella densa* | GBIF.org (22 March 2023) GBIF Occurrence Download https://doi.org/10.15468/dl.2krpnn |
| --- | --- |
| *Selaginella digitata* | GBIF.org (22 March 2023) GBIF Occurrence Download https://doi.org/10.15468/dl.k6eka7 |
| *Selaginella dregei* | GBIF.org (22 March 2023) GBIF Occurrence Download https://doi.org/10.15468/dl.g2dbs3 |
| *Selaginella echinata* | GBIF.org (22 March 2023) GBIF Occurrence Download https://doi.org/10.15468/dl.3rrvn2 |
| *Selaginella eremophila* | GBIF.org (22 March 2023) GBIF Occurrence Download https://doi.org/10.15468/dl.yjqekh |
| *Selaginella helicoclada* | GBIF.org (22 March 2023) GBIF Occurrence Download https://doi.org/10.15468/dl.m4w6rh |
| *Selaginella helvetica* | GBIF.org (22 March 2023) GBIF Occurrence Download https://doi.org/10.15468/dl.z4srkk |
| *Selaginella imbricata* | GBIF.org (22 March 2023) GBIF Occurrence Download https://doi.org/10.15468/dl.gkjxpw |
| *Selaginella lepidophylla* | GBIF.org (22 March 2023) GBIF Occurrence Download https://doi.org/10.15468/dl.hpdjsj |
| *Selaginella nivea* | GBIF.org (22 March 2023) GBIF Occurrence Download https://doi.org/10.15468/dl.4n449x |
| *Selaginella njamnjamensis* | GBIF.org (22 March 2023) GBIF Occurrence Download https://doi.org/10.15468/dl.5uud4g |
| *Selaginella peruviana* | GBIF.org (22 March 2023) GBIF Occurrence Download https://doi.org/10.15468/dl.7zywft |
| *Selaginella phillipsiana* | GBIF.org (22 March 2023) GBIF Occurrence Download https://doi.org/10.15468/dl.j568ks |
| *Selaginella pilifera* | GBIF.org (22 March 2023) GBIF Occurrence Download https://doi.org/10.15468/dl.nfqtwa |
| *Selaginella rupincola* | GBIF.org (22 March 2023) GBIF Occurrence Download https://doi.org/10.15468/dl.p34w5f |
| *Selaginella sartorii* | GBIF.org (22 March 2023) GBIF Occurrence Download https://doi.org/10.15468/dl.t5vsy8 |
| *Selaginella sellowii* | GBIF.org (22 March 2023) GBIF Occurrence Download https://doi.org/10.15468/dl.4zt397 |
| *Selaginella tamariscina* | GBIF.org (22 March 2023) GBIF Occurrence Download https://doi.org/10.15468/dl.dpm3g2 |
| *Selaginella trisulcata* | GBIF.org (22 March 2023) GBIF Occurrence Download https://doi.org/10.15468/dl.m49rbn |
| *Selaginella yemensis* | GBIF.org (22 March 2023) GBIF Occurrence Download https://doi.org/10.15468/dl.b73j9x |
| *Arthropteris orientalis* | GBIF.org (22 March 2023) GBIF Occurrence Download https://doi.org/10.15468/dl.yfu9nx |

**Table S2**(continued)

| *Acanthochlamys bracteata* | GBIF.org (22 March 2023) GBIF Occurrence Download https://doi.org/10.15468/dl.tc6vaj |
| --- | --- |
| *Barbacenia blackii* | GBIF.org (22 March 2023) GBIF Occurrence Download https://doi.org/10.15468/dl.dea6tj |
| *Barbacenia fanniae* | GBIF.org (22 March 2023) GBIF Occurrence Download https://doi.org/10.15468/dl.42336d |
| *Barbacenia flava* | GBIF.org (22 March 2023) GBIF Occurrence Download https://doi.org/10.15468/dl.d7yq3k |
| *Barbacenia fragrans* | GBIF.org (22 March 2023) GBIF Occurrence Download https://doi.org/10.15468/dl.u7jbpr |
| *Barbacenia gentianoides* | GBIF.org (22 March 2023) GBIF Occurrence Download https://doi.org/10.15468/dl.scjhe7 |
| *Barbacenia gounelleana* | GBIF.org (22 March 2023) GBIF Occurrence Download https://doi.org/10.15468/dl.ntkmb3 |
| *Barbacenia graminifolia* | GBIF.org (22 March 2023) GBIF Occurrence Download https://doi.org/10.15468/dl.fu9dsg |
| *Barbacenia longiflora* | GBIF.org (22 March 2023) GBIF Occurrence Download https://doi.org/10.15468/dl.6xsqsg |
| *Barbacenia longiscapa* | GBIF.org (22 March 2023) GBIF Occurrence Download https://doi.org/10.15468/dl.xb5sv7 |
| *Barbacenia macrantha* | GBIF.org (22 March 2023) GBIF Occurrence Download https://doi.org/10.15468/dl.ra3s87 |
| *Barbacenia purpurea* | GBIF.org (22 March 2023) GBIF Occurrence Download https://doi.org/10.15468/dl.strqx6 |
| *Barbacenia riedeliana* | GBIF.org (22 March 2023) GBIF Occurrence Download https://doi.org/10.15468/dl.wnqdnc |
| *Barbacenia seubertiana* | GBIF.org (22 March 2023) GBIF Occurrence Download https://doi.org/10.15468/dl.h885my |
| *Barbacenia spectabilis* | GBIF.org (22 March 2023) GBIF Occurrence Download https://doi.org/10.15468/dl.646as8 |
| *Barbacenia tomentosa* | GBIF.org (22 March 2023) GBIF Occurrence Download https://doi.org/10.15468/dl.qmu23t |
| *Barbaceniopsis boliviensis* | GBIF.org (22 March 2023) GBIF Occurrence Download https://doi.org/10.15468/dl.yfy33v |
| *Barbaceniopsis humahuaquensis* | GBIF.org (22 March 2023) GBIF Occurrence Download https://doi.org/10.15468/dl.w5nw3n |
| *Vellozia albiflora* | GBIF.org (22 March 2023) GBIF Occurrence Download https://doi.org/10.15468/dl.pv5vq2 |
| *Vellozia andina* | GBIF.org (22 March 2023) GBIF Occurrence Download https://doi.org/10.15468/dl.4rf6ce |
| *Vellozia angustifolia* | GBIF.org (22 March 2023) GBIF Occurrence Download https://doi.org/10.15468/dl.3trncs |

**Table S2**(continued)

| *Vellozia candida* | GBIF.org (22 March 2023) GBIF Occurrence Download https://doi.org/10.15468/dl.56bfsv |
| --- | --- |
| *Vellozia caput-ardeae* | GBIF.org (22 March 2023) GBIF Occurrence Download https://doi.org/10.15468/dl.apnykz |
| *Vellozia caruncularis* | GBIF.org (22 March 2023) GBIF Occurrence Download https://doi.org/10.15468/dl.t8vs2q |
| *Vellozia ciliata* | GBIF.org (22 March 2023) GBIF Occurrence Download https://doi.org/10.15468/dl.q6487f |
| *Vellozia compacta* | GBIF.org (22 March 2023) GBIF Occurrence Download https://doi.org/10.15468/dl.ceebkb |
| *Vellozia declinans* | GBIF.org (22 March 2023) GBIF Occurrence Download https://doi.org/10.15468/dl.yjspn5 |
| *Vellozia epidendroides* | GBIF.org (22 March 2023) GBIF Occurrence Download https://doi.org/10.15468/dl.kb97qr |
| *Vellozia flavicans* | GBIF.org (22 March 2023) GBIF Occurrence Download https://doi.org/10.15468/dl.qwzbxg |
| *Vellozia glochidea* | GBIF.org (22 March 2023) GBIF Occurrence Download https://doi.org/10.15468/dl.8trmju |
| *Vellozia hatschbachii* | GBIF.org (22 March 2023) GBIF Occurrence Download https://doi.org/10.15468/dl.sccdse |
| *Vellozia hirsuta* | GBIF.org (22 March 2023) GBIF Occurrence Download https://doi.org/10.15468/dl.4a35fn |
| *Vellozia nanuzae* | GBIF.org (22 March 2023) GBIF Occurrence Download https://doi.org/10.15468/dl.emp5nx |
| *Vellozia nivea* | GBIF.org (22 March 2023) GBIF Occurrence Download https://doi.org/10.15468/dl.wjab47 |
| *Vellozia plicata* | GBIF.org (22 March 2023) GBIF Occurrence Download https://doi.org/10.15468/dl.kjrpja |
| *Vellozia pulchra* | GBIF.org (22 March 2023) GBIF Occurrence Download https://doi.org/10.15468/dl.2j85fg |
| *Vellozia resinosa* | GBIF.org (22 March 2023) GBIF Occurrence Download https://doi.org/10.15468/dl.tt5kbz |
| *Vellozia sellowii* | GBIF.org (22 March 2023) GBIF Occurrence Download https://doi.org/10.15468/dl.atkrnf |
| *Vellozia semirii* | GBIF.org (22 March 2023) GBIF Occurrence Download https://doi.org/10.15468/dl.yn5g4f |
| *Vellozia squalida* | GBIF.org (22 March 2023) GBIF Occurrence Download https://doi.org/10.15468/dl.qmfuxz |
| *Vellozia streptophylla* | GBIF.org (22 March 2023) GBIF Occurrence Download https://doi.org/10.15468/dl.2uk6z3 |
| *Vellozia subscabra* | GBIF.org (22 March 2023) GBIF Occurrence Download https://doi.org/10.15468/dl.v7y6ux |

**Table S2**(continued)

| *Vellozia taxifolia* | GBIF.org (22 March 2023) GBIF Occurrence Download https://doi.org/10.15468/dl.z6cdrs |
| --- | --- |
| *Vellozia tubiflora* | GBIF.org (22 March 2023) GBIF Occurrence Download https://doi.org/10.15468/dl.xsh9uu |
| *Vellozia variabilis* | GBIF.org (22 March 2023) GBIF Occurrence Download https://doi.org/10.15468/dl.h78xms |
| *Vellozia variegata* | GBIF.org (22 March 2023) GBIF Occurrence Download https://doi.org/10.15468/dl.dzhvby |
| *Vellozia verruculosa* | GBIF.org (22 March 2023) GBIF Occurrence Download https://doi.org/10.15468/dl.nbsddt |
| *Xerophyta dasylirioides* | GBIF.org (22 March 2023) GBIF Occurrence Download https://doi.org/10.15468/dl.at9m53 |
| *Xerophyta eglandulosa* | GBIF.org (22 March 2023) GBIF Occurrence Download https://doi.org/10.15468/dl.cfgra3 |
| *Xerophyta elegans* | GBIF.org (22 March 2023) GBIF Occurrence Download https://doi.org/10.15468/dl.pupthf |
| *Xerophyta equisetoides* | GBIF.org (22 March 2023) GBIF Occurrence Download https://doi.org/10.15468/dl.eczrnm |
| *Xerophyta humilis* | GBIF.org (22 March 2023) GBIF Occurrence Download https://doi.org/10.15468/dl.rr4nrq |
| *Xerophyta nandrasanae* | GBIF.org (22 March 2023) GBIF Occurrence Download https://doi.org/10.15468/dl.h42xye |
| *Xerophyta pectinata* | GBIF.org (22 March 2023) GBIF Occurrence Download https://doi.org/10.15468/dl.rfsbps |
| *Xerophyta pinifolia* | GBIF.org (22 March 2023) GBIF Occurrence Download https://doi.org/10.15468/dl.6677cg |
| *Xerophyta retinervis* | GBIF.org (22 March 2023) GBIF Occurrence Download https://doi.org/10.15468/dl.84bsa6 |
| *Xerophyta rippsteinii* | GBIF.org (22 March 2023) GBIF Occurrence Download https://doi.org/10.15468/dl.pdy6fj |
| *Xerophyta scabrida* | GBIF.org (22 March 2023) GBIF Occurrence Download https://doi.org/10.15468/dl.6r72gy |
| *Xerophyta schlechteri* | GBIF.org (22 March 2023) GBIF Occurrence Download https://doi.org/10.15468/dl.2uz55d |
| *Xerophyta schnizleinia* | GBIF.org (22 March 2023) GBIF Occurrence Download https://doi.org/10.15468/dl.suyyuq |
| *Xerophyta spekei* | GBIF.org (22 March 2023) GBIF Occurrence Download https://doi.org/10.15468/dl.e8sbpb |
| *Xerophyta splendens* | GBIF.org (22 March 2023) GBIF Occurrence Download https://doi.org/10.15468/dl.uywgqw |
| *Xerophyta squarrosa* | GBIF.org (22 March 2023) GBIF Occurrence Download https://doi.org/10.15468/dl.rb76jm |

**Table S2**(continued)

| *Xerophyta villosa* | GBIF.org (22 March 2023) GBIF Occurrence Download https://doi.org/10.15468/dl.7vmwrt |
| --- | --- |
| *Xerophyta viscosa* | GBIF.org (22 March 2023) GBIF Occurrence Download https://doi.org/10.15468/dl.23smj6 |
| *Woodsia ilvensis* | GBIF.org (22 March 2023) GBIF Occurrence Download https://doi.org/10.15468/dl.mev4ch |

**Table S3**The relative importance of environmental variables for the desiccation-tolerant vascular plants distribution. VPD – vapor pressure deficit; SRad – solar radiation; MAT – mean annual temperature; DRF – drought frequency; DRI – drought intensity; DRL – drought length.

| Species | MAT | SRAD | VPD | DRF | DRI | DRL |
| --- | --- | --- | --- | --- | --- | --- |
| *Anemia ferruginea* | 63.97 | 13.68 | 6.64 | 5.53 | 4.63 | 5.56 |
| *Anemia flexuosa* | 62.08 | 20.30 | 12.23 | 0.36 | 3.87 | 1.16 |
| *Anemia rotundifolia* | 29.49 | 7.79 | 53.29 | 2.54 | 0.14 | 6.75 |
| *Anemia tomentosa* | 76.74 | 4.00 | 8.34 | 3.87 | 0.57 | 6.49 |
| *Anemia villosa* | 34.20 | 6.02 | 29.04 | 2.03 | 12.98 | 15.72 |
| *Anemia mexicana* | 38.99 | 9.37 | 8.08 | 6.63 | 19.80 | 17.13 |
| *Mohria caffrorum* | 78.08 | 2.50 | 6.43 | 0.13 | 0.20 | 12.66 |
| *Asplenium aethiopicum* | 34.26 | 4.74 | 57.28 | 1.39 | 1.74 | 0.60 |
| *Asplenium adiantum-nigrum* | 52.62 | 2.58 | 36.36 | 1.70 | 3.86 | 2.88 |
| *Asplenium cordatum* | 80.58 | 13.86 | 0.41 | 0.84 | 0.78 | 3.52 |
| *Asplenium dalhousiae* | 36.51 | 8.16 | 4.40 | 18.98 | 5.58 | 26.37 |
| *Asplenium friesiorum* | 27.61 | 1.25 | 61.95 | 0.43 | 4.76 | 4.00 |
| *Asplenium megalura* | 17.17 | 31.81 | 19.69 | 23.05 | 2.49 | 5.79 |
| *Asplenium monanthes* | 30.63 | 11.17 | 47.84 | 2.52 | 1.96 | 5.87 |
| *Asplenium obovatum* | 32.45 | 19.17 | 45.12 | 1.60 | 0.68 | 0.97 |
| *Asplenium pringlei* | 2.93 | 21.48 | 2.03 | 49.10 | 3.85 | 0.61 |
| *Asplenium praegracile* | 1.01 | 12.40 | 38.56 | 4.27 | 43.76 | 0.00 |
| *Asplenium ruta-muraria* | 63.75 | 6.51 | 19.16 | 4.44 | 1.59 | 4.55 |
| *Asplenium rutifolium* | 32.26 | 1.08 | 51.24 | 4.24 | 3.64 | 7.54 |
| *Asplenium sandersonii* | 11.74 | 3.98 | 69.06 | 0.29 | 8.52 | 6.42 |
| *Asplenium septentrionale* | 50.05 | 9.19 | 13.57 | 14.58 | 0.86 | 11.75 |
| *Asplenium trichomanes* | 55.79 | 6.74 | 23.48 | 1.80 | 1.14 | 11.05 |
| *Asplenium theciferum* | 37.84 | 21.52 | 26.51 | 2.26 | 1.42 | 10.46 |
| *Asplenium uhligii* | 60.50 | 16.27 | 1.68 | 18.69 | 1.11 | 1.75 |
| *Asplenium ceterach* | 18.33 | 7.58 | 64.40 | 3.43 | 2.74 | 3.52 |
| *Pleurosorus rutifolius* | 83.53 | 7.40 | 4.05 | 1.19 | 1.64 | 2.19 |
| *Borya constricta* | 58.63 | 3.85 | 2.80 | 1.95 | 28.53 | 4.23 |
| *Borya mirabilis* | 1.23 | 0.10 | 36.85 | 10.87 | 47.14 | 3.81 |
| *Borya nitida* | 0.49 | 25.69 | 31.93 | 0.00 | 0.00 | 41.89 |
| *Borya septentrionalis* | 0.87 | 0.38 | 90.36 | 1.22 | 0.98 | 6.20 |
| *Borya scirpoidea* | 0.45 | 3.52 | 36.14 | 0.46 | 2.04 | 57.39 |
| *Borya sphaerocephala* | 8.64 | 36.76 | 5.57 | 2.29 | 10.06 | 36.68 |
| *Pitcairnia lanuginosa* | 26.33 | 20.19 | 15.38 | 6.27 | 13.81 | 18.02 |
| *Blossfeldia liliputana* | 61.09 | 0.48 | 1.62 | 9.52 | 27.28 | 0.00 |
| *Afrotrilepis pilosa* | 32.82 | 23.54 | 6.29 | 0.35 | 7.13 | 29.88 |
| *Coleochloa abyssinica* | 49.34 | 5.65 | 0.00 | 23.01 | 0.21 | 21.79 |
| *Coleochloa microcephala* | 7.01 | 62.19 | 0.00 | 9.51 | 0.35 | 20.94 |
| *Coleochloa pallidior* | 14.83 | 22.53 | 34.49 | 0.08 | 28.07 | 0.00 |

**Table S3**(continued)

| *Coleochloa setifera* | 12.18 | 33.10 | 49.12 | 3.13 | 1.51 | 0.95 |
| --- | --- | --- | --- | --- | --- | --- |
| *Microdracoides squamosus* | 5.42 | 25.72 | 6.16 | 42.65 | 20.06 | 0.00 |
| *Trilepis ciliatifolia* | 15.41 | 26.06 | 31.27 | 11.80 | 12.80 | 2.66 |
| *Trilepis lhotzkiana* | 50.22 | 5.38 | 32.51 | 1.04 | 4.32 | 6.54 |
| *Trilepis microstachya* | 4.25 | 70.85 | 4.65 | 10.74 | 3.51 | 6.01 |
| *Davallia angustata* | 20.00 | 15.86 | 37.44 | 8.95 | 13.29 | 4.46 |
| *Elaphoglossum acrostichoides* | 22.48 | 2.26 | 67.82 | 0.68 | 2.21 | 4.55 |
| *Elaphoglossum petiolatum* | 62.67 | 14.14 | 15.32 | 1.10 | 2.54 | 4.22 |
| *Elaphoglossum piloselloides* | 81.75 | 5.12 | 8.15 | 1.23 | 2.78 | 0.96 |
| *Boea hygroscopica* | 0.03 | 1.98 | 89.65 | 7.86 | 0.04 | 0.44 |
| *Boea hygrometrica* | 31.55 | 15.78 | 12.30 | 0.22 | 27.01 | 13.15 |
| *Damrongia clarkeana* | 3.65 | 51.60 | 28.62 | 0.12 | 0.00 | 16.01 |
| *Haberlea rhodopensis* | 40.83 | 12.93 | 9.06 | 7.62 | 29.52 | 0.04 |
| *Paraboea crassifolia* | 80.17 | 6.31 | 5.49 | 0.09 | 3.87 | 4.06 |
| *Paraboea rufescens* | 51.04 | 28.81 | 8.84 | 9.54 | 1.05 | 0.73 |
| *Ramonda myconi* | 12.36 | 65.55 | 3.70 | 0.59 | 11.42 | 6.38 |
| *Ramonda nathaliae* | 7.56 | 62.17 | 0.00 | 5.27 | 0.00 | 25.00 |
| *Ramonda serbica* | 3.41 | 51.29 | 0.00 | 0.00 | 3.19 | 2.12 |
| *Crepidomanes frappieri* | 11.31 | 1.45 | 78.01 | 7.75 | 0.33 | 1.14 |
| *Crepidomanes inopinatum* | 44.91 | 11.37 | 41.30 | 1.90 | 0.11 | 0.41 |
| *Hymenophyllum caudiculatum* | 31.99 | 22.92 | 44.23 | 0.44 | 0.09 | 0.33 |
| *Hymenoglossum cruentum* | 20.33 | 7.08 | 71.44 | 0.36 | 0.19 | 0.60 |
| *Hymenophyllum capillare* | 9.41 | 2.53 | 75.80 | 3.28 | 4.48 | 4.49 |
| *Hymenophyllum dentatum* | 5.40 | 1.76 | 89.66 | 1.03 | 0.13 | 2.02 |
| *Hymenophyllum fucoides* | 22.53 | 15.52 | 52.41 | 0.20 | 5.58 | 3.76 |
| *Hymenophyllum hirsutum* | 6.40 | 3.73 | 82.73 | 0.64 | 3.74 | 2.75 |
| *Hymenophyllum kuhnii* | 4.49 | 8.36 | 72.39 | 6.65 | 4.03 | 4.09 |
| *Cardiomanes reniforme* | 1.31 | 2.47 | 92.21 | 1.14 | 2.86 | 0.01 |
| *Hymenophyllum peltatum* | 32.51 | 5.16 | 51.76 | 1.76 | 4.45 | 4.36 |
| *Hymenophyllum polyanthos* | 19.33 | 7.05 | 68.64 | 0.38 | 1.64 | 2.96 |
| *Hymenophyllum plicatum* | 8.64 | 13.83 | 74.90 | 0.62 | 2.00 | 0.01 |
| *Hymenophyllum sanguinolentum* | 35.82 | 11.35 | 29.98 | 11.06 | 4.56 | 7.22 |
| *Hymenophyllum splendidum* | 16.57 | 30.34 | 28.30 | 12.94 | 6.18 | 5.68 |
| *Hymenophyllum tunbrigense* | 35.35 | 2.29 | 58.46 | 0.27 | 0.26 | 3.36 |
| *Polyphlebium borbonicum* | 16.77 | 4.67 | 74.33 | 1.23 | 1.18 | 1.83 |
| *Trichomanes bucinatum* | 17.31 | 34.59 | 20.76 | 14.84 | 12.50 | 0.00 |
| *Crepidomanes chevalieri* | 28.17 | 1.23 | 33.88 | 32.14 | 4.38 | 0.20 |
| *Trichomanes capillaceum* | 44.38 | 17.55 | 26.68 | 0.44 | 1.39 | 9.56 |
| *Trichomanes diaphanum* | 21.22 | 10.46 | 63.99 | 2.62 | 0.85 | 0.85 |
| *Didymoglossum erosum* | 2.40 | 7.71 | 82.02 | 0.22 | 0.13 | 7.51 |
| *Crepidomanes melanotrichum* | 18.57 | 1.19 | 64.39 | 4.61 | 4.91 | 6.33 |

**Table S3**(continued)

| *Trichomanes pyxidiferum* | 54.50 | 6.54 | 35.76 | 0.83 | 1.47 | 0.89 |
| --- | --- | --- | --- | --- | --- | --- |
| *Trichomanes polypodioides* | 34.01 | 6.92 | 42.05 | 1.85 | 2.57 | 12.61 |
| *Trichomanes rigidum* | 31.82 | 12.23 | 48.67 | 1.15 | 2.33 | 3.80 |
| *Trichomanes radicans* | 33.69 | 24.99 | 33.63 | 0.84 | 0.91 | 5.94 |
| *Isoetes australis* | 65.36 | 1.29 | 3.87 | 2.93 | 23.56 | 2.99 |
| *Craterostigma hirsutum* | 65.43 | 2.79 | 0.00 | 0.00 | 31.78 | 0.00 |
| *Craterostigma lanceolatum* | 10.29 | 0.46 | 75.59 | 13.57 | 0.03 | 0.07 |
| *Craterostigma plantagineum* | 32.21 | 6.78 | 45.73 | 4.93 | 9.78 | 0.58 |
| *Craterostigma pumilum* | 58.19 | 0.02 | 18.67 | 0.00 | 21.28 | 1.84 |
| *Craterostigma wilmsii* | 61.43 | 0.00 | 18.46 | 3.18 | 16.92 | 0.00 |
| *Linderniella pulchella* | 33.49 | 30.71 | 14.29 | 3.07 | 18.44 | 0.00 |
| *Linderniella wilmsii* | 34.43 | 35.20 | 10.95 | 5.84 | 0.00 | 13.59 |
| *Myrothamnus flabellifolius* | 42.62 | 42.02 | 4.44 | 2.43 | 2.18 | 6.32 |
| *Myrothamnus moschatus* | 10.19 | 28.16 | 35.68 | 6.02 | 0.44 | 19.51 |
| *Micraira multinervia* | 0.74 | 48.67 | 21.10 | 0.00 | 25.40 | 4.09 |
| *Eragrostiella bifaria* | 10.57 | 50.52 | 18.14 | 20.73 | 0.04 | 0.00 |
| *Eragrostiella brachyphylla* | 0.00 | 1.65 | 89.57 | 7.90 | 0.87 | 0.00 |
| *Eragrostiella nardoides* | 67.08 | 6.76 | 18.30 | 7.86 | 0.00 | 0.00 |
| *Eragrostis nindensis* | 39.98 | 55.75 | 0.05 | 0.71 | 1.41 | 2.10 |
| *Eragrostis paradoxa* | 83.56 | 0.15 | 0.01 | 0.00 | 7.77 | 8.51 |
| *Micraira adamsii* | 31.00 | 36.47 | 12.53 | 9.47 | 8.54 | 2.00 |
| *Microchloa caffra* | 23.10 | 25.20 | 24.81 | 21.20 | 3.37 | 2.31 |
| *Microchloa indica* | 23.40 | 31.96 | 29.66 | 7.85 | 2.97 | 4.14 |
| *Microchloa kunthii* | 40.44 | 9.44 | 38.90 | 1.37 | 0.91 | 8.94 |
| *Micraira lazaridis* | 0.81 | 14.06 | 80.00 | 1.65 | 2.29 | 1.19 |
| *Micrachne patentiflora* | 28.26 | 50.36 | 10.45 | 10.35 | 0.59 | 0.00 |
| *Micraira subulifolia* | 8.84 | 1.06 | 85.93 | 3.30 | 0.08 | 0.79 |
| *Micraira spinifera* | 4.99 | 65.72 | 8.67 | 11.61 | 0.10 | 8.92 |
| *Micraira tenuis* | 0.22 | 67.02 | 13.39 | 7.35 | 5.26 | 6.77 |
| *Micraira viscidula* | 0.04 | 70.76 | 13.29 | 8.77 | 7.13 | 0.00 |
| *Oropetium aristatum* | 4.42 | 5.28 | 80.15 | 4.42 | 3.04 | 2.69 |
| *Oropetium capense* | 59.80 | 3.86 | 6.86 | 18.06 | 6.44 | 4.97 |
| *Oropetium thomaeum* | 6.50 | 1.77 | 64.25 | 24.98 | 0.00 | 2.50 |
| *Sporobolus atrovirens* | 23.03 | 3.39 | 17.89 | 48.99 | 0.27 | 6.44 |
| *Sporobolus elongatus* | 17.26 | 31.20 | 45.53 | 1.47 | 1.84 | 2.70 |
| *Sporobolus festivus* | 10.40 | 2.54 | 29.34 | 16.15 | 12.61 | 28.96 |
| *Sporobolus fimbriatus* | 21.87 | 3.83 | 2.74 | 34.73 | 20.07 | 16.76 |
| *Styppeiochloa hitchcockii* | 73.16 | 3.72 | 5.24 | 2.51 | 1.15 | 14.22 |
| *Sporobolus pellucidus* | 3.14 | 12.74 | 30.92 | 16.54 | 5.16 | 31.50 |
| *Sporobolus ruspolianus* | 10.20 | 34.13 | 55.32 | 0.16 | 0.19 | 0.00 |
| *Sporobolus stapfianus* | 23.32 | 9.18 | 32.07 | 28.45 | 3.61 | 3.38 |

**Table S3**(continued)

| *Tripogon curvatus* | 32.11 | 4.89 | 10.70 | 3.41 | 13.10 | 35.79 |
| --- | --- | --- | --- | --- | --- | --- |
| *Tripogon jacquemontii* | 15.73 | 0.00 | 34.26 | 10.20 | 26.88 | 12.92 |
| *Tripogon major* | 44.32 | 28.99 | 5.04 | 15.61 | 5.80 | 0.24 |
| *Tripogonella minima* | 6.51 | 2.86 | 5.42 | 42.03 | 14.29 | 28.88 |
| *Tripogonella spicata* | 46.13 | 13.16 | 20.24 | 2.80 | 15.41 | 2.26 |
| *Polypodium cambricum* | 30.60 | 15.12 | 48.18 | 4.97 | 0.44 | 0.69 |
| *Ctenopteris heterophylla* | 25.11 | 40.49 | 0.00 | 0.71 | 9.48 | 24.22 |
| *Melpomene flabelliformis* | 8.71 | 6.19 | 72.47 | 4.08 | 0.52 | 8.04 |
| *Loxogramme abyssinica* | 26.88 | 9.86 | 52.00 | 3.26 | 2.85 | 5.14 |
| *Microgramma piloselloides* | 38.10 | 24.82 | 14.26 | 8.17 | 3.10 | 11.55 |
| *Melpomene peruviana* | 71.58 | 14.10 | 10.06 | 1.63 | 0.96 | 1.67 |
| *Pleopeltis angusta* | 25.93 | 2.55 | 21.90 | 22.12 | 17.17 | 10.34 |
| *Pleopeltis crassinervata* | 33.18 | 3.76 | 48.59 | 4.99 | 8.08 | 1.40 |
| *Pecluma eurybasis* | 87.89 | 6.51 | 3.52 | 0.67 | 0.73 | 0.67 |
| *Goniophlebium furfuraceum* | 36.43 | 24.51 | 12.38 | 26.68 | 0.00 | 0.00 |
| *Pleopeltis hirsutissima* | 45.73 | 44.88 | 8.75 | 0.23 | 0.08 | 0.33 |
| *Polypodium interjectum* | 36.41 | 6.73 | 44.02 | 0.75 | 4.38 | 7.70 |
| *Pleopeltis macrocarpa* | 21.90 | 1.88 | 69.67 | 0.77 | 0.64 | 5.15 |
| *Pleopeltis mexicana* | 34.79 | 5.43 | 28.82 | 10.78 | 15.42 | 4.75 |
| *Pleopeltis minima* | 60.62 | 26.62 | 3.72 | 3.41 | 3.47 | 2.16 |
| *Pleopeltis polypodioides* | 2.94 | 72.88 | 12.91 | 0.46 | 3.00 | 7.81 |
| *Pleopeltis plebeia* | 45.66 | 3.61 | 28.70 | 2.07 | 14.23 | 5.72 |
| *Pleopeltis pleopeltifolia* | 61.20 | 22.08 | 15.30 | 0.61 | 0.73 | 0.07 |
| *Polypodium remotum* | 82.96 | 3.54 | 8.73 | 1.52 | 2.69 | 0.57 |
| *Polypodium virginianum* | 56.94 | 14.17 | 14.40 | 0.31 | 8.48 | 5.70 |
| *Polypodium vulgare* | 53.64 | 2.40 | 39.29 | 3.23 | 0.42 | 1.02 |
| *Platycerium stemaria* | 15.76 | 4.25 | 72.66 | 0.57 | 2.72 | 4.03 |
| *Actiniopteris dimorpha* | 6.22 | 21.14 | 42.88 | 2.44 | 12.36 | 14.96 |
| *Argyrochosma fendleri* | 5.67 | 40.57 | 25.83 | 1.89 | 7.38 | 18.65 |
| *Adiantum hispidulum* | 5.44 | 9.54 | 40.15 | 15.72 | 13.53 | 15.61 |
| *Adiantum incisum* | 12.09 | 6.59 | 43.38 | 7.96 | 17.48 | 12.50 |
| *Adiantum latifolium* | 30.41 | 23.46 | 31.34 | 1.08 | 1.99 | 11.72 |
| *Actiniopteris radiata* | 13.19 | 5.49 | 32.51 | 10.18 | 25.12 | 13.51 |
| *Adiantum raddianum* | 76.87 | 14.47 | 6.88 | 1.07 | 0.30 | 0.41 |
| *Actiniopteris semiflabellata* | 6.15 | 14.32 | 57.22 | 2.34 | 3.14 | 16.84 |
| *Astrolepis sinuata* | 52.44 | 35.72 | 4.75 | 1.51 | 2.44 | 3.13 |
| *Bommeria hispida* | 30.28 | 55.55 | 9.23 | 0.82 | 0.60 | 3.51 |
| *Cheilanthes capensis* | 5.00 | 30.30 | 0.71 | 3.98 | 2.68 | 57.34 |
| *Cheilanthes distans* | 22.12 | 2.53 | 70.91 | 2.54 | 0.31 | 1.58 |
| *Cheilanthes depauperata* | 5.03 | 70.96 | 0.00 | 1.95 | 0.00 | 22.06 |
| *Cheilanthes dinteri* | 5.86 | 16.67 | 36.09 | 5.33 | 13.44 | 22.61 |

**Table S3**(continued)

| *Cheilanthes buchtienii* | 57.66 | 7.60 | 24.12 | 0.00 | 8.78 | 1.85 |
| --- | --- | --- | --- | --- | --- | --- |
| *Cheilanthes eckloniana* | 83.14 | 4.93 | 0.67 | 8.63 | 2.42 | 0.20 |
| *Aleuritopteris farinosa* | 57.37 | 3.09 | 36.26 | 1.55 | 0.00 | 1.72 |
| *Cheilanthes glauca* | 16.78 | 11.52 | 18.18 | 41.89 | 0.99 | 10.64 |
| *Cheilanthes gracillima* | 15.66 | 31.02 | 24.66 | 20.83 | 4.27 | 3.56 |
| *Cheilanthes hirta* | 44.03 | 33.21 | 7.87 | 6.63 | 7.88 | 0.38 |
| *Astrolepis cochisensis* | 34.90 | 38.70 | 10.67 | 6.82 | 1.15 | 7.76 |
| *Astrolepis integerrima* | 31.06 | 27.63 | 24.72 | 3.64 | 3.83 | 9.12 |
| *Cheilanthes lasiophylla* | 34.56 | 13.73 | 17.80 | 9.85 | 4.92 | 19.15 |
| *Cheilanthes multifida* | 60.37 | 12.92 | 24.45 | 0.00 | 1.46 | 0.80 |
| *Cheilanthes marginata* | 74.49 | 5.98 | 12.65 | 0.82 | 5.62 | 0.44 |
| *Cheilanthes parviloba* | 63.22 | 9.00 | 0.21 | 11.47 | 16.11 | 0.00 |
| *Cheilanthes sieberi* | 67.54 | 16.88 | 7.99 | 1.77 | 0.64 | 5.18 |
| *Cheilanthes tenuifolia* | 40.68 | 1.39 | 33.52 | 2.22 | 2.19 | 20.00 |
| *Cosentinia vellea* | 16.01 | 23.30 | 45.74 | 12.23 | 2.01 | 0.72 |
| *Cheilanthes catanensis* | 9.26 | 13.47 | 68.02 | 8.42 | 0.83 | 0.00 |
| *Doryopteris collina* | 52.25 | 1.19 | 43.09 | 0.44 | 0.63 | 2.41 |
| *Doryopteris concolor* | 80.64 | 13.82 | 1.00 | 2.59 | 0.82 | 1.13 |
| *Cheilanthes notholaenoides* | 64.51 | 21.75 | 9.45 | 0.74 | 2.35 | 1.19 |
| *Doryopteris varians* | 13.45 | 4.61 | 64.96 | 2.90 | 2.18 | 11.90 |
| *Cheilanthes quadripinnata* | 50.39 | 12.52 | 4.90 | 9.23 | 9.65 | 13.30 |
| *Aleuritopteris albomarginata* | 54.75 | 20.62 | 9.47 | 13.93 | 1.04 | 0.20 |
| *Pellaea atropurpurea* | 61.85 | 17.57 | 3.19 | 0.10 | 9.85 | 7.43 |
| *Pellaea andromedifolia* | 47.59 | 17.12 | 7.65 | 6.03 | 2.41 | 19.20 |
| *Cheilanthes bonariensis* | 64.18 | 12.69 | 9.06 | 2.25 | 3.75 | 8.07 |
| *Pellaea sagittata* | 79.73 | 10.83 | 2.12 | 0.58 | 1.92 | 4.82 |
| *Cheilanthes fragillima* | 1.17 | 41.60 | 21.23 | 10.30 | 0.42 | 25.29 |
| *Pellaea glabella* | 40.21 | 29.09 | 23.26 | 0.99 | 4.37 | 2.08 |
| *Cheilanthes inaequalis* | 80.66 | 12.07 | 0.68 | 3.36 | 0.84 | 2.39 |
| *Doryopteris kitchingii* | 96.73 | 0.44 | 0.22 | 2.17 | 0.00 | 0.44 |
| *Cheilanthes lendigera* | 51.94 | 26.96 | 14.86 | 0.50 | 1.77 | 3.97 |
| *Cheilanthes marlothii* | 0.19 | 59.44 | 38.10 | 0.12 | 2.15 | 0.00 |
| *Cheilanthes myriophylla* | 68.56 | 12.12 | 11.97 | 0.78 | 1.87 | 4.69 |
| *Paragymnopteris marantae* | 63.09 | 1.95 | 17.21 | 14.24 | 0.00 | 3.51 |
| *Allosorus coriaceus* | 58.17 | 36.61 | 0.00 | 3.19 | 2.03 | 0.00 |
| *Cheilanthes nitidula* | 52.42 | 20.50 | 0.00 | 0.00 | 0.12 | 26.96 |
| *Pellaea ovata* | 53.58 | 3.62 | 6.71 | 8.73 | 8.45 | 18.91 |
| *Cheilanthes parryi* | 1.45 | 60.02 | 24.76 | 1.05 | 6.07 | 6.65 |
| *Cheilanthes pringlei* | 37.54 | 24.98 | 21.28 | 7.80 | 2.62 | 5.78 |
| *Doryopteris pedata* | 40.45 | 15.02 | 15.11 | 6.71 | 21.88 | 0.82 |
| *Hemionitis palmata* | 16.90 | 57.15 | 10.52 | 2.48 | 7.42 | 5.53 |

**Table S3**(continued)

| *Allosorus pteridioides* | 23.08 | 35.05 | 0.57 | 7.57 | 30.49 | 3.24 |
| --- | --- | --- | --- | --- | --- | --- |
| *Hemionitis tomentosa* | 66.46 | 9.48 | 15.87 | 2.36 | 0.34 | 5.49 |
| *Cheilanthes tomentosa* | 63.92 | 7.93 | 0.82 | 1.22 | 1.63 | 24.48 |
| *Doryopteris triphylla* | 35.42 | 48.89 | 2.97 | 6.95 | 0.34 | 5.43 |
| *Pellaea ternifolia* | 43.16 | 10.50 | 6.79 | 10.75 | 10.44 | 18.36 |
| *Pentagramma triangularis* | 53.88 | 7.80 | 12.88 | 12.28 | 1.18 | 11.98 |
| *Haplopteris volkensii* | 15.43 | 6.51 | 62.55 | 15.02 | 0.49 | 0.00 |
| *Cheilanthes wrightii* | 18.04 | 37.67 | 20.98 | 12.29 | 2.38 | 8.65 |
| *Myriopteris rufa* | 22.67 | 53.77 | 0.87 | 3.00 | 11.90 | 7.79 |
| *Notholaena lanuginosa* | 38.60 | 48.79 | 11.91 | 0.70 | 0.00 | 0.00 |
| *Notholaena muelleri* | 7.39 | 16.70 | 68.06 | 1.19 | 3.91 | 2.75 |
| *Negripteris scioana* | 42.62 | 19.37 | 15.44 | 8.80 | 13.25 | 0.52 |
| *Onychium divaricatum* | 16.06 | 15.81 | 34.72 | 0.00 | 19.93 | 13.47 |
| *Pellaea boivinii* | 14.19 | 32.21 | 45.11 | 7.25 | 0.36 | 0.88 |
| *Pellaea brachyptera* | 17.87 | 30.66 | 6.14 | 28.40 | 3.36 | 13.57 |
| *Pellaea bridgesii* | 7.22 | 57.99 | 18.42 | 13.23 | 0.96 | 2.18 |
| *Pellaea calomelanos* | 34.70 | 20.20 | 41.59 | 1.36 | 1.11 | 1.05 |
| *Pellaea dura* | 19.90 | 13.37 | 62.76 | 2.44 | 1.50 | 0.04 |
| *Pellaea falcata* | 11.07 | 3.88 | 80.76 | 0.35 | 1.01 | 2.94 |
| *Pellaea longipilosa* | 44.96 | 15.29 | 2.31 | 6.47 | 28.08 | 2.89 |
| *Pellaea mucronata* | 20.46 | 28.74 | 6.56 | 22.41 | 3.76 | 18.07 |
| *Pellaea pectiniformis* | 18.61 | 19.76 | 51.07 | 0.78 | 8.03 | 1.76 |
| *Pellaea rotundifolia* | 45.04 | 9.81 | 2.48 | 18.94 | 5.39 | 18.33 |
| *Pellaea truncata* | 8.39 | 81.01 | 2.72 | 0.86 | 4.21 | 2.81 |
| *Cheilanthes viridis* | 24.98 | 3.79 | 40.94 | 7.18 | 8.98 | 14.12 |
| *Pellaea wrightiana* | 16.42 | 76.36 | 1.38 | 0.49 | 1.55 | 3.81 |
| *Vittaria guineensis* | 0.89 | 27.17 | 58.51 | 6.78 | 3.99 | 2.65 |
| *Vittaria isoetifolia* | 5.94 | 11.70 | 78.37 | 1.67 | 0.55 | 1.76 |
| *Schizaea pusilla* | 48.77 | 23.31 | 23.28 | 0.49 | 2.01 | 2.13 |
| *Selaginella arizonica* | 15.21 | 26.00 | 27.41 | 9.85 | 3.65 | 17.89 |
| *Selaginella bryopteris* | 9.81 | 29.07 | 22.89 | 11.26 | 7.14 | 19.82 |
| *Selaginella convoluta* | 48.84 | 26.46 | 10.09 | 2.58 | 2.75 | 9.28 |
| *Selaginella caffrorum* | 70.42 | 3.92 | 19.05 | 6.56 | 0.00 | 0.05 |
| *Selaginella digitata* | 18.73 | 22.94 | 3.07 | 28.69 | 2.16 | 24.41 |
| *Selaginella dregei* | 3.26 | 18.50 | 39.44 | 1.51 | 0.00 | 37.29 |
| *Selaginella densa* | 38.92 | 16.72 | 9.01 | 6.93 | 10.32 | 18.10 |
| *Selaginella echinata* | 13.83 | 31.00 | 8.48 | 13.50 | 9.37 | 23.82 |
| *Selaginella eremophila* | 17.25 | 67.24 | 7.13 | 3.13 | 1.84 | 3.41 |
| *Selaginella helvetica* | 35.02 | 24.50 | 3.32 | 8.50 | 14.81 | 13.85 |
| *Selaginella helicoclada* | 11.06 | 31.33 | 17.17 | 6.01 | 0.54 | 33.89 |
| *Selaginella imbricata* | 7.76 | 13.41 | 54.50 | 0.19 | 21.04 | 3.10 |

**Table S3**(continued)

| *Selaginella lepidophylla* | 11.04 | 12.58 | 19.55 | 1.54 | 17.67 | 37.62 |
| --- | --- | --- | --- | --- | --- | --- |
| *Selaginella nivea* | 1.72 | 8.16 | 6.77 | 81.78 | 0.00 | 1.57 |
| *Selaginella njamnjamensis* | 11.80 | 50.76 | 13.65 | 22.97 | 0.45 | 0.37 |
| *Selaginella peruviana* | 62.97 | 5.39 | 9.53 | 5.44 | 4.17 | 12.49 |
| *Selaginella pilifera* | 11.62 | 19.71 | 9.22 | 48.21 | 2.42 | 8.82 |
| *Selaginella phillipsiana* | 68.14 | 15.90 | 0.84 | 7.27 | 6.51 | 1.35 |
| *Selaginella rupincola* | 30.57 | 44.80 | 4.07 | 6.77 | 10.77 | 3.02 |
| *Selaginella sellowii* | 52.58 | 23.05 | 12.66 | 2.06 | 8.09 | 1.57 |
| *Selaginella sartorii* | 27.24 | 65.62 | 0.31 | 0.88 | 1.82 | 4.13 |
| *Selaginella tamariscina* | 18.24 | 4.02 | 6.37 | 19.17 | 20.62 | 31.58 |
| *Selaginella trisulcata* | 83.86 | 13.37 | 1.61 | 0.18 | 0.01 | 0.97 |
| *Selaginella yemensis* | 58.01 | 12.49 | 15.90 | 0.07 | 0.32 | 13.21 |
| *Arthropteris orientalis* | 17.21 | 2.46 | 72.51 | 2.44 | 2.39 | 3.00 |
| *Acanthochlamys bracteata* | 52.45 | 24.14 | 5.99 | 0.90 | 16.52 | 0.00 |
| *Barbacenia blackii* | 56.46 | 5.99 | 6.14 | 17.09 | 4.24 | 10.09 |
| *Barbacenia flava* | 56.13 | 4.84 | 4.07 | 9.63 | 5.82 | 19.51 |
| *Barbacenia fragrans* | 32.74 | 0.06 | 42.83 | 22.68 | 1.09 | 0.60 |
| *Barbacenia longiflora* | 61.95 | 3.75 | 0.09 | 34.17 | 0.04 | 0.00 |
| *Barbacenia gentianoides* | 59.99 | 2.64 | 3.07 | 29.37 | 0.22 | 4.71 |
| *Barbacenia graminifolia* | 44.93 | 13.49 | 2.40 | 36.86 | 0.73 | 1.58 |
| *Barbaceniopsis humahuaquensis* | 23.83 | 2.13 | 32.65 | 19.06 | 19.71 | 2.62 |
| *Barbacenia longiscapa* | 49.59 | 0.91 | 0.45 | 1.81 | 47.24 | 0.00 |
| *Barbacenia macrantha* | 43.95 | 2.79 | 8.36 | 34.00 | 1.95 | 8.94 |
| *Barbacenia purpurea* | 33.29 | 39.30 | 22.79 | 3.95 | 0.04 | 0.63 |
| *Barbacenia riedeliana* | 49.79 | 2.22 | 0.00 | 38.67 | 0.02 | 9.29 |
| *Barbacenia seubertiana* | 10.84 | 0.02 | 10.93 | 77.62 | 0.55 | 0.04 |
| *Barbacenia spectabilis* | 38.21 | 0.00 | 16.98 | 3.75 | 39.68 | 1.38 |
| *Barbacenia tomentosa* | 33.01 | 4.36 | 7.25 | 5.32 | 0.00 | 50.06 |
| *Barbacenia gounelleana* | 0.84 | 0.64 | 1.41 | 96.62 | 0.00 | 0.50 |
| *Xerophyta elegans* | 13.41 | 49.87 | 31.16 | 0.34 | 1.86 | 3.36 |
| *Vellozia variabilis* | 54.66 | 22.27 | 2.78 | 5.69 | 3.01 | 11.58 |
| *Vellozia albiflora* | 22.62 | 15.79 | 14.41 | 2.93 | 18.23 | 26.02 |
| *Vellozia angustifolia* | 71.72 | 14.00 | 4.06 | 1.19 | 6.67 | 2.36 |
| *Vellozia andina* | 82.03 | 6.06 | 1.11 | 9.77 | 0.77 | 0.26 |
| *Barbaceniopsis boliviensis* | 58.15 | 23.03 | 14.94 | 0.26 | 3.62 | 0.00 |
| *Vellozia candida* | 20.49 | 43.62 | 32.34 | 0.53 | 2.01 | 1.01 |
| *Vellozia ciliata* | 20.35 | 16.81 | 0.00 | 46.85 | 16.00 | 0.00 |
| *Vellozia caput-ardeae* | 56.31 | 0.00 | 0.00 | 42.89 | 0.00 | 0.81 |
| *Vellozia caruncularis* | 34.92 | 2.99 | 6.33 | 7.89 | 13.32 | 34.54 |
| *Vellozia compacta* | 24.49 | 6.11 | 5.19 | 7.93 | 8.70 | 47.57 |
| *Vellozia declinans* | 48.90 | 4.36 | 1.43 | 40.00 | 2.75 | 2.56 |

**Table S3**(continued)

| *Vellozia epidendroides* | 41.29 | 4.60 | 2.26 | 41.81 | 3.73 | 6.31 |
| --- | --- | --- | --- | --- | --- | --- |
| *Vellozia glochidea* | 14.70 | 11.31 | 21.34 | 23.21 | 28.27 | 1.17 |
| *Vellozia hatschbachii* | 10.90 | 52.09 | 13.84 | 2.47 | 11.38 | 9.32 |
| *Vellozia hirsuta* | 52.01 | 2.57 | 3.87 | 11.86 | 21.50 | 8.18 |
| *Vellozia nanuzae* | 43.39 | 1.28 | 0.64 | 41.27 | 8.56 | 4.86 |
| *Vellozia nivea* | 53.57 | 0.81 | 1.35 | 29.00 | 2.76 | 12.52 |
| *Vellozia plicata* | 31.28 | 7.43 | 12.04 | 2.17 | 31.26 | 15.82 |
| *Vellozia pulchra* | 6.18 | 52.23 | 3.79 | 8.78 | 17.28 | 11.73 |
| *Vellozia resinosa* | 51.43 | 4.63 | 1.84 | 34.54 | 1.17 | 6.40 |
| *Vellozia flavicans* | 19.13 | 62.46 | 3.10 | 1.85 | 13.01 | 0.45 |
| *Vellozia sellowii* | 42.47 | 5.38 | 3.29 | 12.28 | 5.62 | 30.95 |
| *Vellozia squalida* | 77.76 | 1.81 | 5.51 | 0.23 | 11.05 | 3.63 |
| *Vellozia subscabra* | 51.94 | 29.62 | 9.02 | 0.21 | 9.21 | 0.00 |
| *Vellozia taxifolia* | 43.29 | 0.19 | 0.03 | 48.53 | 0.68 | 7.28 |
| *Vellozia tubiflora* | 15.39 | 38.98 | 7.65 | 0.83 | 20.33 | 16.82 |
| *Vellozia variegata* | 4.77 | 19.03 | 69.41 | 4.42 | 1.43 | 0.94 |
| *Vellozia verruculosa* | 83.70 | 0.18 | 10.85 | 2.81 | 2.47 | 0.00 |
| *Xerophyta dasylirioides* | 18.36 | 18.19 | 53.05 | 2.39 | 3.30 | 4.71 |
| *Xerophyta eglandulosa* | 62.57 | 25.23 | 0.00 | 2.66 | 6.92 | 2.62 |
| *Xerophyta equisetoides* | 72.60 | 0.00 | 21.77 | 4.39 | 0.00 | 1.24 |
| *Xerophyta humilis* | 41.36 | 46.57 | 7.51 | 4.56 | 0.00 | 0.00 |
| *Xerophyta pinifolia* | 7.40 | 51.36 | 2.28 | 30.14 | 0.00 | 8.83 |
| *Xerophyta pectinata* | 31.30 | 30.42 | 10.20 | 0.76 | 6.72 | 20.59 |
| *Xerophyta retinervis* | 44.52 | 29.63 | 19.66 | 0.00 | 0.02 | 6.17 |
| *Xerophyta squarrosa* | 15.42 | 40.71 | 3.83 | 1.15 | 0.43 | 38.46 |
| *Xerophyta schlechteri* | 15.41 | 51.88 | 3.85 | 1.23 | 2.08 | 25.54 |
| *Xerophyta scabrida* | 5.31 | 70.46 | 0.00 | 9.65 | 0.00 | 14.58 |
| *Xerophyta schnizleinia* | 24.71 | 23.63 | 49.17 | 0.84 | 1.50 | 0.14 |
| *Xerophyta splendens* | 0.22 | 0.00 | 63.07 | 0.03 | 0.11 | 36.57 |
| *Xerophyta villosa* | 0.04 | 32.22 | 20.58 | 25.13 | 1.13 | 20.91 |
| *Xerophyta viscosa* | 25.74 | 20.00 | 54.26 | 0.00 | 0.00 | 0.00 |
| *Woodsia ilvensis* | 18.46 | 53.69 | 14.21 | 0.93 | 9.77 | 2.93 |

Table S4. The mean exposure of desiccation-tolerant vascular plants to shifts in climate. ΔVPD – historical climatic variability in vapor pressure deficit; ΔSRad – historical climatic variability in solar radiation; ΔMAT – historical climatic variability in mean annual temperature; ΔDRF – historical climatic variability in drought frequency; ΔDRI – historical climatic variability in drought intensity; ΔDRL – historical climatic variability in drought length

| *Species* | ΔVPD | ΔSRad | ΔMAT | ΔDRF | ΔDRI | ΔDRL |
| --- | --- | --- | --- | --- | --- | --- |
| *Anemia ferruginea* | 0.02 | 6.4 | 0.48 | 0.53 | 0.5 | 0.33 |
| *Anemia flexuosa* | 0.02 | 4.8 | 0.27 | 0.52 | 0.38 | 0.27 |
| *Anemia mexicana* | 0.08 | 5.94 | 0.72 | 0.58 | 0.35 | 0.28 |
| *Anemia rotundifolia* | 0.01 | 6.4 | 0.4 | 0.54 | 0.47 | 0.34 |
| *Anemia tomentosa* | 0.02 | 4.12 | 0.45 | 0.51 | 0.4 | 0.27 |
| *Anemia villosa* | 0.02 | 5.68 | 0.46 | 0.64 | 0.4 | 0.31 |
| *Mohria caffrorum* | 0.04 | 1.55 | 0.59 | 0.89 | 0.47 | 0.45 |
| *Asplenium adiantum-nigrum* | 0.06 | 3.85 | 0.97 | 0.84 | 0.51 | 0.37 |
| *Asplenium aethiopicum* | 0.04 | 3.83 | 0.74 | 0.66 | 0.57 | 0.38 |
| *Asplenium ceterach* | 0.07 | 4.06 | 0.93 | 1.13 | 0.69 | 0.54 |
| *Asplenium cordatum* | 0.07 | 2.43 | 0.7 | 0.87 | 0.58 | 0.49 |
| *Asplenium dalhousiae* | 0.05 | 1.85 | 0.7 | 0.63 | 0.38 | 0.29 |
| *Asplenium friesiorum* | 0.05 | 3.85 | 0.75 | 0.71 | 0.55 | 0.37 |
| *Asplenium megalura* | 0.05 | 6.55 | 0.76 | 0.82 | 0.54 | 0.42 |
| *Asplenium monanthes* | 0.03 | 4.4 | 0.55 | 0.68 | 0.49 | 0.37 |
| *Asplenium obovatum* | 0.07 | 3.28 | 0.88 | 1.12 | 0.72 | 0.51 |
| *Asplenium praegracile* | 0.02 | 2.65 | 0.66 | 0.26 | 0.38 | 0.2 |
| *Asplenium pringlei* | 0.06 | 4.85 | 0.77 | 0.85 | 0.84 | 0.6 |
| *Asplenium ruta-muraria* | 0.05 | 3.73 | 1.06 | 0.76 | 0.43 | 0.31 |
| *Asplenium rutifolium* | 0.05 | 3.35 | 0.7 | 0.81 | 0.54 | 0.44 |
| *Asplenium sandersonii* | 0.05 | 5.83 | 0.75 | 0.53 | 0.43 | 0.26 |
| *Asplenium septentrionale* | 0.05 | 3.76 | 0.99 | 0.62 | 0.33 | 0.27 |
| *Asplenium theciferum* | 0.02 | 7.31 | 0.45 | 0.53 | 0.41 | 0.25 |
| *Asplenium trichomanes* | 0.04 | 3.02 | 0.9 | 0.71 | 0.37 | 0.28 |
| *Asplenium uhligii* | 0.03 | 3.66 | 0.76 | 0.62 | 0.54 | 0.31 |
| *Pleurosorus rutifolius* | 0.04 | 2.21 | 0.51 | 0.71 | 0.35 | 0.41 |
| *Borya constricta* | 0.04 | 3.3 | 0.4 | 0.59 | 0.39 | 0.35 |
| *Borya inopinata* | 0.03 | 2.72 | 0.48 | 0.74 | 0.08 | 0.29 |
| *Borya mirabilis* | 0.02 | 2.37 | 0.44 | 1.15 | 0.44 | 0.58 |
| *Borya nitida* | 0.01 | 2.87 | 0.34 | 0.71 | 0.48 | 0.44 |
| *Borya scirpoidea* | 0.01 | 2.74 | 0.35 | 0.73 | 0.49 | 0.44 |
| *Borya septentrionalis* | 0 | 1.28 | 0.42 | 0.38 | 0.12 | 0.13 |
| *Borya sphaerocephala* | 0.03 | 2.52 | 0.37 | 0.71 | 0.45 | 0.39 |
| *Pitcairnia lanuginosa* | 0.03 | 5.8 | 0.41 | 0.82 | 0.84 | 0.62 |
| *Blossfeldia liliputana* | 0.02 | 1.64 | 0.42 | 0.33 | 0.15 | 0.13 |

**Table S4**(continued)

| *Afrotrilepis pilosa* | 0.03 | 8.05 | 0.53 | 0.72 | 0.37 | 0.42 |
| --- | --- | --- | --- | --- | --- | --- |
| *Coleochloa abyssinica* | 0.05 | 5.77 | 0.75 | 0.49 | 0.28 | 0.26 |
| *Coleochloa microcephala* | 0.02 | 5.57 | 0.56 | 0.55 | 0.1 | 0.33 |
| *Coleochloa pallidior* | 0.03 | 1.2 | 0.76 | 1.03 | 0.74 | 0.56 |
| *Coleochloa setifera* | 0.04 | 1.64 | 0.65 | 0.86 | 0.6 | 0.43 |
| *Microdracoides squamosus* | 0.04 | 9.88 | 0.51 | 1.35 | 0.25 | 0.68 |
| *Trilepis ciliatifolia* | 0.01 | 2.18 | 0.34 | 0.49 | 0.28 | 0.36 |
| *Trilepis lhotzkiana* | 0.01 | 7.63 | 0.39 | 0.59 | 0.42 | 0.33 |
| *Trilepis microstachya* | 0.01 | 4.28 | 0.42 | 0.49 | 0.45 | 0.36 |
| *Davallia angustata* | 0.01 | 10.24 | 0.71 | 0.35 | 0.21 | 0.13 |
| *Elaphoglossum acrostichoides* | 0.04 | 2.63 | 0.66 | 0.79 | 0.51 | 0.43 |
| *Elaphoglossum petiolatum* | 0.04 | 6.79 | 0.62 | 0.58 | 0.44 | 0.36 |
| *Elaphoglossum piloselloides* | 0.02 | 5.19 | 0.45 | 0.58 | 0.4 | 0.31 |
| *Boea hygrometrica* | 0.08 | 3.57 | 0.77 | 0.5 | 0.31 | 0.24 |
| *Boea hygroscopica* | 0.01 | 1.73 | 0.46 | 0.55 | 0.21 | 0.2 |
| *Damrongia clarkeana* | 0.05 | 2.56 | 0.49 | 0.38 | 0.25 | 0.18 |
| *Haberlea rhodopensis* | 0.07 | 2.75 | 0.83 | 0.57 | 0.3 | 0.16 |
| *Oreocharis mileensis* | 0.07 | 6.95 | 0.69 | 0.6 | 0.04 | 0.18 |
| *Paraboea crassifolia* | 0.04 | 6.84 | 0.38 | 0.33 | 0.18 | 0.13 |
| *Paraboea rufescens* | 0.05 | 7.91 | 0.46 | 0.3 | 0.16 | 0.16 |
| *Ramonda myconi* | 0.08 | 7.66 | 1.08 | 1.28 | 0.84 | 0.76 |
| *Ramonda nathaliae* | 0.08 | 2.4 | 0.83 | 1.11 | 0.45 | 0.25 |
| *Ramonda serbica* | 0.06 | 1.2 | 0.64 | 1.05 | 0.53 | 0.27 |
| *Cardiomanes reniforme* | 0 | 1.77 | 0.24 | 0.36 | 0.39 | 0.16 |
| *Crepidomanes chevalieri* | 0.04 | 5.2 | 0.73 | 0.53 | 0.39 | 0.34 |
| *Crepidomanes frappieri* | 0.03 | 1.98 | 0.59 | 0.44 | 0.4 | 0.13 |
| *Crepidomanes inopinatum* | 0.06 | 5.46 | 0.78 | 0.74 | 0.6 | 0.45 |
| *Crepidomanes melanotrichum* | 0.05 | 3.27 | 0.69 | 0.78 | 0.53 | 0.43 |
| *Didymoglossum erosum* | 0.05 | 8.91 | 0.65 | 0.49 | 0.34 | 0.26 |
| *Hymenoglossum cruentum* | 0.01 | 3.11 | 0.18 | 0.76 | 0.51 | 0.4 |
| *Hymenophyllum capillare* | 0.04 | 4.34 | 0.72 | 0.67 | 0.5 | 0.36 |
| *Hymenophyllum caudiculatum* | 0.01 | 3.95 | 0.5 | 0.72 | 0.47 | 0.38 |
| *Hymenophyllum dentatum* | 0.01 | 2.83 | 0.2 | 0.7 | 0.43 | 0.38 |
| *Hymenophyllum fucoides* | 0.02 | 6.49 | 0.48 | 0.61 | 0.44 | 0.32 |
| *Hymenophyllum hirsutum* | 0.03 | 6.49 | 0.49 | 0.57 | 0.44 | 0.29 |
| *Hymenophyllum kuhnii* | 0.05 | 6.62 | 0.78 | 0.62 | 0.46 | 0.32 |
| *Hymenophyllum peltatum* | 0.01 | 2.59 | 0.38 | 0.68 | 0.39 | 0.36 |
| *Hymenophyllum plicatum* | 0.01 | 2.41 | 0.18 | 0.73 | 0.46 | 0.39 |
| *Hymenophyllum polyanthos* | 0.02 | 5.88 | 0.46 | 0.59 | 0.48 | 0.31 |
| *Hymenophyllum sanguinolentum* | 0 | 2.19 | 0.25 | 0.36 | 0.33 | 0.13 |

**Table S4**(continued)

| *Hymenophyllum splendidum* | 0.04 | 7.59 | 0.69 | 0.71 | 0.36 | 0.34 |
| --- | --- | --- | --- | --- | --- | --- |
| *Hymenophyllum tunbrigense* | 0.06 | 4.71 | 0.82 | 0.96 | 0.59 | 0.46 |
| *Polyphlebium borbonicum* | 0.05 | 4.34 | 0.71 | 0.57 | 0.47 | 0.31 |
| *Trichomanes bucinatum* | 0.05 | 11.14 | 0.74 | 0.7 | 0.2 | 0.16 |
| *Trichomanes capillaceum* | 0.03 | 6.2 | 0.54 | 0.48 | 0.38 | 0.25 |
| *Trichomanes diaphanum* | 0.02 | 6.26 | 0.46 | 0.58 | 0.42 | 0.3 |
| *Trichomanes polypodioides* | 0.02 | 5.76 | 0.49 | 0.57 | 0.44 | 0.31 |
| *Trichomanes pyxidiferum* | 0.02 | 4.28 | 0.51 | 0.56 | 0.42 | 0.29 |
| *Trichomanes radicans* | 0.03 | 4.91 | 0.52 | 0.58 | 0.45 | 0.32 |
| *Trichomanes rigidum* | 0.02 | 5.86 | 0.5 | 0.56 | 0.45 | 0.29 |
| *Isoetes australis* | 0.03 | 3.38 | 0.39 | 0.59 | 0.38 | 0.36 |
| *Craterostigma hirsutum* | 0.04 | 2.82 | 0.73 | 0.69 | 0.67 | 0.48 |
| *Craterostigma lanceolatum* | 0.05 | 7.13 | 0.88 | 0.76 | 0.71 | 0.42 |
| *Craterostigma plantagineum* | 0.06 | 4.03 | 0.8 | 0.68 | 0.59 | 0.29 |
| *Craterostigma pumilum* | 0.03 | 1.16 | 0.66 | 0.76 | 1.2 | 0.31 |
| *Craterostigma wilmsii* | 0.06 | 1.84 | 0.73 | 1.25 | 0.88 | 0.66 |
| *Lindernia brevidens* | 0.03 | 3.05 | 0.58 | 0.44 | 0.19 | 0.21 |
| *Lindernia intrepidus* | 0.12 | 4.98 | 0.9 | 0.27 | 0.27 | 0.25 |
| *Lindernia monroi* | 0.06 | 1.38 | 0.82 | 1.37 | 0.69 | 0.61 |
| *Lindernia purpurea* | 0.04 | 11.28 | 1.05 | 0.53 | 0.3 | 0.07 |
| *Linderniella pulchella* | 0.05 | 1.87 | 0.77 | 1.15 | 0.61 | 0.55 |
| *Linderniella wilmsii* | 0.05 | 1.25 | 0.69 | 0.97 | 0.41 | 0.57 |
| *Myrothamnus flabellifolius* | 0.06 | 2.97 | 0.82 | 1.16 | 0.67 | 0.56 |
| *Myrothamnus moschatus* | 0.01 | 1.47 | 0.54 | 0.57 | 0.24 | 0.12 |
| *Eragrostiella bifaria* | 0.03 | 2.8 | 0.45 | 0.41 | 0.21 | 0.22 |
| *Eragrostiella brachyphylla* | 0.01 | 5.39 | 0.45 | 0.26 | 0.09 | 0.11 |
| *Eragrostiella nardoides* | 0.02 | 2.87 | 0.48 | 0.48 | 0.49 | 0.28 |
| *Eragrostis nindensis* | 0.08 | 2.47 | 0.73 | 1.09 | 0.71 | 0.56 |
| *Eragrostis paradoxa* | 0.04 | 1.54 | 0.88 | 0.87 | 0.33 | 0.48 |
| *Micrachne patentiflora* | 0.04 | 2.35 | 0.95 | 1.27 | 0.47 | 0.52 |
| *Micraira adamsii* | 0.07 | 1.61 | 0.26 | 1.01 | 0.49 | 0.54 |
| *Micraira lazaridis* | 0.01 | 1.54 | 0.34 | 1.2 | 1.2 | 0.41 |
| *Micraira multinervia* | 0.07 | 0.82 | 0.26 | 1.11 | 0.43 | 0.55 |
| *Micraira spinifera* | 0.07 | 1.19 | 0.26 | 0.99 | 0.63 | 0.55 |
| *Micraira subulifolia* | 0.01 | 1.18 | 0.44 | 0.58 | 0.19 | 0.23 |
| *Micraira tenuis* | 0.07 | 1.15 | 0.27 | 0.96 | 0.32 | 0.51 |
| *Micraira viscidula* | 0.07 | 1.49 | 0.27 | 1 | 0.53 | 0.55 |
| *Microchloa caffra* | 0.06 | 3.02 | 0.72 | 1.02 | 0.64 | 0.56 |
| *Microchloa indica* | 0.04 | 5.87 | 0.51 | 0.61 | 0.43 | 0.34 |

**Table S4**(continued)

| *Microchloa kunthii* | 0.05 | 3.45 | 0.68 | 0.79 | 0.52 | 0.43 |
| --- | --- | --- | --- | --- | --- | --- |
| *Oropetium aristatum* | 0.04 | 7.71 | 0.58 | 0.74 | 0.56 | 0.37 |
| *Oropetium capense* | 0.08 | 2.41 | 0.73 | 1.02 | 0.61 | 0.54 |
| *Oropetium roxburghianum* | 0.02 | 2.83 | 0.38 | 0.47 | 0.34 | 0.22 |
| *Oropetium thomaeum* | 0.02 | 3.84 | 0.51 | 0.43 | 0.22 | 0.19 |
| *Sporobolus atrovirens* | 0.05 | 6.9 | 0.67 | 0.45 | 0.26 | 0.27 |
| *Sporobolus elongatus* | 0.03 | 2.69 | 0.56 | 0.92 | 0.33 | 0.46 |
| *Sporobolus festivus* | 0.05 | 5.77 | 0.68 | 0.67 | 0.44 | 0.36 |
| *Sporobolus fimbriatus* | 0.07 | 2.53 | 0.74 | 0.95 | 0.58 | 0.52 |
| *Sporobolus pellucidus* | 0.05 | 2.35 | 0.7 | 0.74 | 0.61 | 0.42 |
| *Sporobolus ruspolianus* | 0.05 | 0.71 | 0.4 | 2.34 | 1.7 | 0.54 |
| *Sporobolus stapfianus* | 0.06 | 3.23 | 0.77 | 1.01 | 0.68 | 0.58 |
| *Styppeiochloa hitchcockii* | 0.01 | 0.84 | 0.56 | 0.42 | 0.31 | 0.09 |
| *Tripogon capillatus* | 0.03 | 2.42 | 0.44 | 0.33 | 0.32 | 0.16 |
| *Tripogon curvatus* | 0.04 | 1.1 | 0.78 | 0.09 | 0.2 | 0.24 |
| *Tripogon filiformis* | 0.03 | 3.11 | 0.51 | 0.3 | 0.24 | 0.12 |
| *Tripogon jacquemontii* | 0.01 | 4 | 0.29 | 0.28 | 0.29 | 0.31 |
| *Tripogon lisboae* | 0.02 | 3.81 | 0.33 | 0.28 | 0.31 | 0.16 |
| *Tripogon major* | 0.03 | 6.69 | 0.56 | 1.18 | 0.42 | 0.67 |
| *Tripogonella loliiformis* | 0.04 | 1.81 | 0.55 | 0.88 | 0.35 | 0.3 |
| *Tripogonella minima* | 0.06 | 6 | 0.64 | 0.85 | 0.52 | 0.44 |
| *Tripogonella spicata* | 0.03 | 4.39 | 0.43 | 0.57 | 0.33 | 0.26 |
| *Ctenopteris heterophylla* | 0 | 1.26 | 0.22 | 0.32 | 0.3 | 0.18 |
| *Goniophlebium furfuraceum* | 0.01 | 2.83 | 0.45 | 0.46 | 0.34 | 0.14 |
| *Loxogramme abyssinica* | 0.05 | 5.86 | 0.71 | 0.62 | 0.49 | 0.36 |
| *Loxogramme lanceolata* | 0.02 | 1.28 | 0.63 | 1.21 | 0.24 | 0.2 |
| *Melpomene flabelliformis* | 0.02 | 6.25 | 0.49 | 0.62 | 0.45 | 0.31 |
| *Melpomene peruviana* | 0.01 | 5.08 | 0.38 | 0.56 | 0.36 | 0.28 |
| *Microgramma piloselloides* | 0.03 | 9.96 | 0.39 | 0.53 | 0.69 | 0.36 |
| *Pecluma eurybasis* | 0.02 | 7.92 | 0.29 | 0.57 | 0.39 | 0.28 |
| *Platycerium stemaria* | 0.06 | 11 | 0.58 | 0.64 | 0.47 | 0.31 |
| *Pleopeltis angusta* | 0.03 | 4.08 | 0.69 | 0.54 | 0.43 | 0.29 |
| *Pleopeltis crassinervata* | 0.04 | 6.66 | 0.67 | 0.5 | 0.3 | 0.21 |
| *Pleopeltis hirsutissima* | 0.01 | 3.36 | 0.53 | 0.6 | 0.41 | 0.31 |
| *Pleopeltis macrocarpa* | 0.03 | 4.34 | 0.55 | 0.65 | 0.47 | 0.35 |
| *Pleopeltis mexicana* | 0.04 | 6.45 | 0.71 | 0.48 | 0.36 | 0.27 |
| *Pleopeltis minima* | 0.02 | 3.48 | 0.46 | 0.46 | 0.36 | 0.24 |
| *Pleopeltis plebeia* | 0.04 | 6.27 | 0.69 | 0.43 | 0.3 | 0.24 |
| *Pleopeltis pleopeltifolia* | 0.01 | 3.88 | 0.49 | 0.54 | 0.4 | 0.27 |

**Table S4**(continued)

| *Pleopeltis polypodioides* | 0.04 | 3.79 | 0.61 | 0.41 | 0.41 | 0.22 |
| --- | --- | --- | --- | --- | --- | --- |
| *Polypodium cambricum* | 0.07 | 3.75 | 0.88 | 1.33 | 0.88 | 0.72 |
| *Polypodium interjectum* | 0.08 | 5.54 | 1.05 | 1.3 | 0.67 | 0.52 |
| *Polypodium remotum* | 0.02 | 8.45 | 0.31 | 0.52 | 0.46 | 0.33 |
| *Polypodium virginianum* | 0.02 | 1.64 | 0.79 | 0.82 | 0.33 | 0.22 |
| *Polypodium vulgare* | 0.04 | 3.19 | 1.06 | 0.8 | 0.38 | 0.3 |
| *Actiniopteris australis* | 0.02 | 1.28 | 0.63 | 1.21 | 0.24 | 0.2 |
| *Actiniopteris dimorpha* | 0.06 | 2.74 | 0.73 | 0.82 | 0.53 | 0.43 |
| *Actiniopteris radiata* | 0.05 | 3.35 | 0.71 | 0.75 | 0.49 | 0.39 |
| *Actiniopteris semiflabellata* | 0.06 | 3.04 | 0.7 | 0.68 | 0.62 | 0.36 |
| *Adiantum hispidulum* | 0.03 | 2.8 | 0.51 | 0.95 | 0.47 | 0.4 |
| *Adiantum incisum* | 0.05 | 4.67 | 0.72 | 0.82 | 0.55 | 0.41 |
| *Adiantum latifolium* | 0.03 | 5.92 | 0.44 | 0.56 | 0.44 | 0.3 |
| *Adiantum raddianum* | 0.01 | 4.61 | 0.46 | 0.53 | 0.41 | 0.29 |
| *Aleuritopteris albomarginata* | 0.04 | 3.86 | 0.5 | 0.45 | 0.3 | 0.21 |
| *Aleuritopteris farinosa* | 0.05 | 3.95 | 0.76 | 0.7 | 0.6 | 0.42 |
| *Allosorus coriaceus* | 0.07 | 1.44 | 0.68 | 1.14 | 1.39 | 0.36 |
| *Allosorus pteridioides* | 0.1 | 4.66 | 1.09 | 2.01 | 1.24 | 1.04 |
| *Argyrochosma fendleri* | 0.02 | 1.73 | 0.73 | 0.34 | 0.17 | 0.21 |
| *Astrolepis cochisensis* | 0.07 | 1.21 | 0.98 | 0.72 | 0.35 | 0.34 |
| *Astrolepis integerrima* | 0.07 | 2.37 | 0.9 | 0.69 | 0.33 | 0.34 |
| *Astrolepis sinuata* | 0.06 | 2.56 | 0.88 | 0.71 | 0.42 | 0.4 |
| *Bommeria hispida* | 0.06 | 2.04 | 1.02 | 0.79 | 0.43 | 0.45 |
| *Cheilanthes bonariensis* | 0.05 | 4.33 | 0.73 | 0.7 | 0.48 | 0.41 |
| *Cheilanthes buchtienii* | 0.02 | 4.15 | 0.29 | 0.43 | 0.3 | 0.24 |
| *Cheilanthes capensis* | 0.06 | 1.3 | 0.6 | 1.11 | 0.64 | 0.56 |
| *Cheilanthes catanensis* | 0.06 | 2.38 | 0.77 | 1.43 | 1.1 | 0.86 |
| *Cheilanthes depauperata* | 0.06 | 1.23 | 0.65 | 1.4 | 0.75 | 0.73 |
| *Cheilanthes dinteri* | 0.11 | 5.01 | 0.94 | 0.74 | 0.33 | 0.25 |
| *Cheilanthes distans* | 0.03 | 2.2 | 0.48 | 0.71 | 0.31 | 0.4 |
| *Cheilanthes eckloniana* | 0.06 | 2.18 | 0.62 | 1.01 | 0.6 | 0.59 |
| *Cheilanthes fragillima* | 0.06 | 1.16 | 0.24 | 0.87 | 0.48 | 0.44 |
| *Cheilanthes glauca* | 0.01 | 1.77 | 0.34 | 0.86 | 0.6 | 0.51 |
| *Cheilanthes gracillima* | 0.04 | 1.08 | 0.59 | 0.51 | 0.2 | 0.3 |
| *Cheilanthes hirta* | 0.05 | 1.55 | 0.63 | 0.8 | 0.53 | 0.38 |
| *Cheilanthes inaequalis* | 0.06 | 4.44 | 0.8 | 0.87 | 0.67 | 0.5 |
| *Cheilanthes lasiophylla* | 0.06 | 2.3 | 0.53 | 0.53 | 0.37 | 0.31 |
| *Cheilanthes lendigera* | 0.04 | 5.44 | 0.74 | 0.7 | 0.46 | 0.42 |
| *Cheilanthes marginata* | 0.03 | 5.87 | 0.44 | 0.51 | 0.37 | 0.28 |

**Table S4**(continued)

| *Cheilanthes marlothii* | 0.09 | 3.97 | 0.92 | 0.28 | 0.25 | 0.21 |
| --- | --- | --- | --- | --- | --- | --- |
| *Cheilanthes multifida* | 0.03 | 1.24 | 0.8 | 0.52 | 0.42 | 0.23 |
| *Cheilanthes myriophylla* | 0.03 | 5.88 | 0.52 | 0.64 | 0.47 | 0.37 |
| *Cheilanthes nitidula* | 0.04 | 3.58 | 0.57 | 0.29 | 0.2 | 0.17 |
| *Cheilanthes notholaenoides* | 0.05 | 5.43 | 0.69 | 0.55 | 0.35 | 0.32 |
| *Cheilanthes parryi* | 0.06 | 0.98 | 0.87 | 0.42 | 0.25 | 0.18 |
| *Cheilanthes parviloba* | 0.05 | 1 | 0.49 | 1.09 | 0.58 | 0.5 |
| *Cheilanthes pringlei* | 0.07 | 1.58 | 1.08 | 0.84 | 0.51 | 0.55 |
| *Cheilanthes quadripinnata* | 0.05 | 2.27 | 0.65 | 0.83 | 0.55 | 0.47 |
| *Cheilanthes sieberi* | 0.04 | 2.13 | 0.53 | 0.64 | 0.35 | 0.38 |
| *Cheilanthes tenuifolia* | 0.04 | 4.44 | 0.54 | 0.43 | 0.35 | 0.22 |
| *Cheilanthes tomentosa* | 0.04 | 1.04 | 0.82 | 0.58 | 0.21 | 0.18 |
| *Cheilanthes viridis* | 0.06 | 3.26 | 0.72 | 0.84 | 0.57 | 0.45 |
| *Cheilanthes wrightii* | 0.07 | 1.35 | 1.09 | 0.8 | 0.45 | 0.44 |
| *Cosentinia vellea* | 0.07 | 3.3 | 0.87 | 1.52 | 1.12 | 0.94 |
| *Doryopteris collina* | 0.02 | 5.05 | 0.44 | 0.68 | 0.48 | 0.34 |
| *Doryopteris concolor* | 0.02 | 3.88 | 0.46 | 0.46 | 0.37 | 0.25 |
| *Doryopteris kitchingii* | 0.01 | 0.47 | 0.59 | 0.35 | 0.34 | 0.06 |
| *Doryopteris pedata* | 0.01 | 2.13 | 0.62 | 0.61 | 1.02 | 0.44 |
| *Doryopteris triphylla* | 0.01 | 2.15 | 0.42 | 0.58 | 0.39 | 0.27 |
| *Doryopteris varians* | 0.02 | 4.13 | 0.48 | 0.56 | 0.44 | 0.34 |
| *Haplopteris volkensii* | 0.05 | 5.06 | 0.84 | 0.61 | 0.53 | 0.32 |
| *Hemionitis palmata* | 0.04 | 3.32 | 0.56 | 0.63 | 0.4 | 0.29 |
| *Hemionitis tomentosa* | 0.02 | 4.54 | 0.41 | 0.48 | 0.4 | 0.26 |
| *Myriopteris rufa* | 0.05 | 1.91 | 0.96 | 0.67 | 0.32 | 0.34 |
| *Negripteris scioana* | 0.1 | 1.29 | 0.68 | 0.17 | 0.89 | 0.1 |
| *Notholaena dipinnata* | 0.08 | 4.87 | 0.8 | 0.45 | 0.31 | 0.32 |
| *Notholaena lanuginosa* | 0.06 | 2.02 | 0.82 | 1.51 | 1.07 | 0.95 |
| *Notholaena muelleri* | 0.02 | 1.65 | 0.5 | 0.5 | 0.18 | 0.22 |
| *Onychium divaricatum* | 0.07 | 1.38 | 0.71 | 0.68 | 0.86 | 0.38 |
| *Paragymnopteris marantae* | 0.04 | 2.03 | 0.54 | 0.53 | 0.38 | 0.23 |
| *Pellaea andromedifolia* | 0.02 | 0.9 | 0.53 | 0.47 | 0.44 | 0.21 |
| *Pellaea atropurpurea* | 0.02 | 1.23 | 0.8 | 0.68 | 0.32 | 0.23 |
| *Pellaea boivinii* | 0.02 | 1.51 | 0.54 | 0.59 | 0.38 | 0.21 |
| *Pellaea brachyptera* | 0.03 | 0.66 | 0.6 | 0.47 | 0.22 | 0.22 |
| *Pellaea bridgesii* | 0.03 | 0.55 | 0.71 | 0.33 | 0.32 | 0.17 |
| *Pellaea calomelanos* | 0.06 | 3.06 | 0.76 | 0.9 | 0.61 | 0.47 |
| *Pellaea dura* | 0.06 | 7.37 | 0.76 | 0.45 | 0.43 | 0.23 |
| *Pellaea falcata* | 0.02 | 2.27 | 0.47 | 0.84 | 0.32 | 0.42 |

**Table S4**(continued)

| *Pellaea glabella* | 0.02 | 1.32 | 0.8 | 0.81 | 0.38 | 0.25 |
| --- | --- | --- | --- | --- | --- | --- |
| *Pellaea longipilosa* | 0.05 | 2.23 | 0.8 | 1.12 | 0.68 | 0.71 |
| *Pellaea mucronata* | 0.02 | 0.78 | 0.59 | 0.41 | 0.4 | 0.19 |
| *Pellaea ovata* | 0.04 | 4.15 | 0.63 | 0.56 | 0.4 | 0.3 |
| *Pellaea pectiniformis* | 0.05 | 2.94 | 0.73 | 0.99 | 0.67 | 0.55 |
| *Pellaea rotundifolia* | 0 | 2.13 | 0.24 | 0.29 | 0.25 | 0.14 |
| *Pellaea sagittata* | 0.03 | 6.08 | 0.55 | 0.56 | 0.38 | 0.32 |
| *Pellaea ternifolia* | 0.03 | 4.83 | 0.56 | 0.62 | 0.49 | 0.37 |
| *Pellaea truncata* | 0.06 | 1.24 | 1.01 | 0.6 | 0.29 | 0.34 |
| *Pellaea wrightiana* | 0.06 | 1.5 | 1 | 0.7 | 0.39 | 0.4 |
| *Pentagramma triangularis* | 0.04 | 1.31 | 0.63 | 0.5 | 0.32 | 0.25 |
| *Vittaria guineensis* | 0.07 | 11.46 | 0.65 | 0.71 | 0.56 | 0.33 |
| *Vittaria isoetifolia* | 0.04 | 1.76 | 0.59 | 0.82 | 0.49 | 0.43 |
| *Schizaea pusilla* | 0.03 | 1.32 | 0.66 | 0.49 | 0.24 | 0.17 |
| *Selaginella arizonica* | 0.07 | 1.39 | 1.04 | 0.62 | 0.3 | 0.3 |
| *Selaginella bryopteris* | 0.09 | 2.72 | 0.77 | 1.2 | 1.06 | 0.58 |
| *Selaginella caffrorum* | 0.06 | 2.48 | 0.6 | 0.92 | 0.57 | 0.55 |
| *Selaginella convoluta* | 0.04 | 6.88 | 0.42 | 0.72 | 0.51 | 0.34 |
| *Selaginella densa* | 0.02 | 1.44 | 0.56 | 0.59 | 0.26 | 0.21 |
| *Selaginella digitata* | 0.04 | 1.87 | 0.56 | 0.94 | 0.32 | 0.19 |
| *Selaginella dregei* | 0.05 | 2.28 | 0.74 | 1.11 | 0.66 | 0.56 |
| *Selaginella echinata* | 0.02 | 1.32 | 0.56 | 0.62 | 0.3 | 0.13 |
| *Selaginella eremophila* | 0.03 | 1.09 | 0.75 | 0.4 | 0.42 | 0.25 |
| *Selaginella helicoclada* | 0.05 | 1.61 | 0.62 | 1 | 0.3 | 0.15 |
| *Selaginella helvetica* | 0.05 | 4.6 | 0.98 | 0.52 | 0.36 | 0.27 |
| *Selaginella imbricata* | 0.11 | 1.78 | 0.72 | 1.03 | 1.44 | 0.63 |
| *Selaginella lepidophylla* | 0.07 | 3.16 | 0.82 | 0.8 | 0.5 | 0.44 |
| *Selaginella nivea* | 0.03 | 2.41 | 0.53 | 0.96 | 0.16 | 0.14 |
| *Selaginella njamnjamensis* | 0.02 | 6.56 | 0.51 | 0.45 | 0.21 | 0.21 |
| *Selaginella peruviana* | 0.05 | 2.5 | 0.78 | 0.62 | 0.36 | 0.33 |
| *Selaginella phillipsiana* | 0.06 | 1.39 | 0.69 | 0.5 | 0.63 | 0.22 |
| *Selaginella pilifera* | 0.07 | 3.01 | 0.82 | 0.74 | 0.35 | 0.33 |
| *Selaginella rupincola* | 0.07 | 3.73 | 0.94 | 0.82 | 0.54 | 0.52 |
| *Selaginella sartorii* | 0.07 | 5.37 | 0.85 | 0.79 | 0.52 | 0.5 |
| *Selaginella sellowii* | 0.02 | 4.34 | 0.34 | 0.51 | 0.35 | 0.25 |
| *Selaginella tamariscina* | 0.06 | 4.08 | 0.81 | 0.31 | 0.24 | 0.17 |
| *Selaginella trisulcata* | 0.02 | 7.74 | 0.29 | 0.5 | 0.41 | 0.29 |
| *Selaginella yemensis* | 0.08 | 1.37 | 0.67 | 0.88 | 1.34 | 0.29 |
| *Arthropteris orientalis* | 0.05 | 5.85 | 0.73 | 0.79 | 0.53 | 0.43 |

**Table S4**(continued)

| *Acanthochlamys bracteata* | 0.02 | 1.71 | 0.48 | 0.33 | 0.16 | 0.1 |
| --- | --- | --- | --- | --- | --- | --- |
| *Barbacenia blackii* | 0 | 12.28 | 0.41 | 0.21 | 0.24 | 0.2 |
| *Barbacenia fanniae* | 0.01 | 2.73 | 0.32 | 0.72 | 0.32 | 0.5 |
| *Barbacenia flava* | 0.01 | 10.76 | 0.43 | 0.41 | 0.37 | 0.29 |
| *Barbacenia fragrans* | 0 | 3.8 | 0.53 | 0.16 | 0.2 | 0.26 |
| *Barbacenia gentianoides* | 0.01 | 13.99 | 0.4 | 0.12 | 0.13 | 0.12 |
| *Barbacenia gounelleana* | 0.01 | 1.9 | 0.48 | 0.05 | 0.14 | 0.19 |
| *Barbacenia graminifolia* | 0.01 | 13.2 | 0.41 | 0.26 | 0.23 | 0.17 |
| *Barbacenia longiflora* | 0.01 | 14.14 | 0.41 | 0.13 | 0.11 | 0.1 |
| *Barbacenia longiscapa* | 0 | 14.08 | 0.41 | 0.06 | 0.09 | 0.11 |
| *Barbacenia macrantha* | 0.01 | 13.47 | 0.41 | 0.15 | 0.18 | 0.12 |
| *Barbacenia purpurea* | 0.01 | 5.14 | 0.41 | 0.34 | 0.37 | 0.25 |
| *Barbacenia riedeliana* | 0.01 | 13.59 | 0.42 | 0.17 | 0.15 | 0.12 |
| *Barbacenia seubertiana* | 0.01 | 0.96 | 0.31 | 0.36 | 0.33 | 0.33 |
| *Barbacenia spectabilis* | 0 | 6.2 | 0.27 | 0.13 | 0.56 | 0.07 |
| *Barbacenia tomentosa* | 0.01 | 9.88 | 0.4 | 0.57 | 0.52 | 0.38 |
| *Barbaceniopsis boliviensis* | 0.02 | 3.58 | 0.21 | 0.42 | 0.25 | 0.14 |
| *Barbaceniopsis humahuaquensis* | 0.02 | 1.56 | 0.46 | 0.35 | 0.29 | 0.12 |
| *Vellozia albiflora* | 0.01 | 9.37 | 0.41 | 0.49 | 0.45 | 0.32 |
| *Vellozia andina* | 0.02 | 3.38 | 0.17 | 0.14 | 0.06 | 0.27 |
| *Vellozia angustifolia* | 0.01 | 10.64 | 0.47 | 0.51 | 0.66 | 0.43 |
| *Vellozia candida* | 0.01 | 7.09 | 0.34 | 0.39 | 0.45 | 0.25 |
| *Vellozia caput-ardeae* | 0 | 14.11 | 0.41 | 0.06 | 0.1 | 0.1 |
| *Vellozia caruncularis* | 0.01 | 11.18 | 0.43 | 0.45 | 0.4 | 0.31 |
| *Vellozia ciliata* | 0.01 | 13.89 | 0.44 | 0.56 | 0.44 | 0.33 |
| *Vellozia compacta* | 0.01 | 10.89 | 0.43 | 0.48 | 0.45 | 0.33 |
| *Vellozia declinans* | 0.01 | 12.53 | 0.46 | 0.24 | 0.38 | 0.19 |
| *Vellozia epidendroides* | 0.01 | 12.79 | 0.41 | 0.23 | 0.23 | 0.15 |
| *Vellozia flavicans* | 0.02 | 10.71 | 0.53 | 0.8 | 0.86 | 0.61 |
| *Vellozia glochidea* | 0.02 | 9.39 | 0.57 | 0.76 | 0.96 | 0.61 |
| *Vellozia hatschbachii* | 0 | 14.21 | 0.41 | 0.06 | 0.1 | 0.09 |
| *Vellozia hirsuta* | 0.02 | 12.86 | 0.45 | 0.75 | 0.54 | 0.41 |
| *Vellozia nanuzae* | 0.01 | 12.24 | 0.39 | 0.43 | 0.37 | 0.29 |
| *Vellozia nivea* | 0.01 | 12.01 | 0.43 | 0.21 | 0.25 | 0.17 |
| *Vellozia plicata* | 0.04 | 6.73 | 0.34 | 0.69 | 0.48 | 0.27 |
| *Vellozia pulchra* | 0.02 | 9.87 | 0.35 | 0.3 | 0.33 | 0.15 |
| *Vellozia resinosa* | 0.01 | 13.16 | 0.41 | 0.27 | 0.25 | 0.19 |
| *Vellozia sellowii* | 0.01 | 11.48 | 0.41 | 0.57 | 0.52 | 0.39 |
| *Vellozia semirii* | 0.01 | 14.11 | 0.4 | 0.11 | 0.11 | 0.1 |

**Table S4** (continued)

| *Vellozia squalida* | 0 | 8.28 | 0.46 | 0.48 | 0.64 | 0.36 |
| --- | --- | --- | --- | --- | --- | --- |
| *Vellozia streptophylla* | 0.01 | 13.91 | 0.44 | 0.12 | 0.13 | 0.08 |
| *Vellozia subscabra* | 0.01 | 8.36 | 0.48 | 0.4 | 0.36 | 0.33 |
| *Vellozia taxifolia* | 0.01 | 14.27 | 0.39 | 0.13 | 0.11 | 0.1 |
| *Vellozia tubiflora* | 0.02 | 9.16 | 0.49 | 0.69 | 0.6 | 0.43 |
| *Vellozia variabilis* | 0.02 | 9.67 | 0.49 | 0.57 | 0.61 | 0.42 |
| *Vellozia variegata* | 0.01 | 6.48 | 0.34 | 0.57 | 0.55 | 0.34 |
| *Vellozia verruculosa* | 0.01 | 8.03 | 0.37 | 0.65 | 0.62 | 0.43 |
| *Xerophyta dasylirioides* | 0.02 | 0.64 | 0.59 | 0.37 | 0.34 | 0.08 |
| *Xerophyta eglandulosa* | 0.02 | 0.34 | 0.66 | 0.14 | 0.35 | 0.05 |
| *Xerophyta elegans* | 0.04 | 2.68 | 0.57 | 0.81 | 0.35 | 0.51 |
| *Xerophyta equisetoides* | 0.05 | 2.21 | 0.9 | 1.29 | 0.87 | 0.86 |
| *Xerophyta humilis* | 0.07 | 3.23 | 0.9 | 0.98 | 0.64 | 0.5 |
| *Xerophyta nandrasanae* | 0.02 | 0.36 | 0.62 | 0.51 | 0.34 | 0.04 |
| *Xerophyta pectinata* | 0.02 | 1.09 | 0.57 | 0.54 | 0.29 | 0.1 |
| *Xerophyta pinifolia* | 0.02 | 2.44 | 0.49 | 0.78 | 0.04 | 0.16 |
| *Xerophyta retinervis* | 0.07 | 1.49 | 0.7 | 1.12 | 0.78 | 0.66 |
| *Xerophyta rippsteinii* | 0.04 | 1.16 | 0.67 | 0.74 | 1.24 | 0.3 |
| *Xerophyta scabrida* | 0.04 | 4.24 | 0.63 | 1.04 | 0.6 | 0.54 |
| *Xerophyta schlechteri* | 0.07 | 1.1 | 0.76 | 1.35 | 1.03 | 0.74 |
| *Xerophyta schnizleinia* | 0.07 | 2 | 0.66 | 0.66 | 0.62 | 0.35 |
| *Xerophyta spekei* | 0.05 | 3.74 | 0.64 | 0.58 | 0.36 | 0.37 |
| *Xerophyta splendens* | 0.02 | 2.11 | 0.69 | 1.49 | 0.75 | 0.62 |
| *Xerophyta squarrosa* | 0.07 | 1.12 | 0.76 | 1.34 | 1.03 | 0.73 |
| *Xerophyta villosa* | 0.07 | 1.47 | 0.77 | 1.29 | 0.95 | 0.72 |
| *Xerophyta viscosa* | 0.04 | 4.72 | 0.63 | 0.8 | 0.66 | 0.74 |
| *Woodsia ilvensis* | 0.02 | 1.75 | 0.79 | 0.85 | 0.34 | 0.24 |

Table S5. List of R packages used in this study

| R package | Reference |
| --- | --- |
| ade4 | Dray, S., Dufour, A. (2007). The ade4 Package: Implementing the Duality Diagram for Ecologists. Journal of Statistical Software, 22(4), 1-20. doi:10.18637/jss.v022.i04. |
| dismo | Hijmans, R.J., Phillips, S., Leathwick, J., Elith, J. (2022). dismo: Species Distribution Modeling. R package version 1.3-9. Available at https://CRAN.R-project.org/package=dismo. |
| dplyr | Wickham, H., François, R., Henry, L., Müller, K., & Vaughan, D. (2023). dplyr: A Grammar of Data Manipulation. R package version 1.1.4. Available at https://CRAN.R-project.org/package=dplyr. |
| egg | Auguie, B. (2019). egg: Extensions for 'ggplot2': Custom Geom, Custom Themes, Plot Alignment, Labelled Panels, Symmetric Scales, and Fixed Panel Size. R package version 0.4.5. Available at https://CRAN.R-project.org/package=egg. |
| ggplot2 | Wickham, H. 2016. ggplot2: Elegant Graphics for Data Analysis. Springer-Verlag New York. |
| ggpubr | Kassambara, A. (2022). ggpubr: 'ggplot2' Based Publication Ready Plots. R package version 0.5.0. Available at https://CRAN.R-project.org/package=ggpubr. |
| htmlwidgets | Vaidyanathan, R., Xie, Y., Allaire, J., Cheng, J., Sievert, C., Russell, K. (2023). htmlwidgets: HTML Widgets for R. R package version 1.6.4. Available at https://CRAN.R-project.org/package=htmlwidgets. |
| humboldt | Brown, J. (2023). humboldt: Analysis of Species in Environmental Space. R package version 1.0.0.0420121. |
| leaflet | Cheng, J., Karambelkar, B., Xie, Y. (2022). leaflet: Create Interactive Web Maps with the JavaScript 'Leaflet' Library. R package version 2.1.1. . Available at https://CRAN.R-project.org/package=leaflet. |
| maps | Becker, O., Minka A., Deckmyn, A. (2021). maps: Draw Geographical Maps. R package version 3.4.0. Available at https://CRAN.R-project.org/package=maps. |
| maptools | Bivand, R., Lewin-Koh, N. (2022). maptools: Tools for Handling Spatial Objects. R package version 1.1-6. Available at https://CRAN.R-project.org/package=maptools. |
| phylobase | Hackathon, R. et al. (2020). phylobase: Base Package for Phylogenetic Structures and Comparative Data. R package version 0.8.10. Available at https://CRAN.R-project.org/package=phylobase. |
| phyloregion | Daru, B.H., Karunarathne, P., & Schliep, K. (2020), phyloregion: R package for biogeographic regionalization and macroecology. Methods in Ecology and Evolution, 11: 1483-1491. doi:10.1111/2041-210X.13478 |

**Table S5**( continued)

| phylosignal | Keck, F., Rimet, F., Bouchez A., & Franc, A. (2016). phylosignal: an R package to measure, test, and explore the phylogenetic signal. Ecology and Evolution, 6(9), 2774-2780. doi:10.1002/ece3.2051 |
| --- | --- |
| raster | Hijmans, R. (2023). raster: Geographic Data Analysis and Modeling.R package version 3.6-20. Available at https://CRAN.R-project.org/package=raster. |
| RColorBrewer | Neuwirth, E. (2022). RColorBrewer: ColorBrewer Palettes. R package version 1.1-3. Available at https://CRAN.R-project.org/package=RColorBrewer. |
| SDMtune | Vignali, S., Barras, A. G., Arlettaz, R., Braunisch, V. (2022). SDMtune: An R package to tune and evaluate species distribution models. Ecology and Evololution, 10(20), 11488–11506. https://doi.org/10.1002/ece3.6786 |
| V.PhyloMaker | Jin, Y., & Qian H. (2019). V.PhyloMaker: an R package that can generate very large phylogenies for vascular plants. Ecography 42: 1353–1359. doi: 10.1111/ecog.04434 |
| vegan | Oksanen, J., Simpson, G., Blanchet, F., Kindt, R., Legendre, P., Minchin, P., … Borman, T. (2022). vegan: Community Ecology Package. R package version 2.6-2. https://github.com/vegandevs/vegan |
| viridis | Garnier, S., Ross, N., Rudis, R., Camargo, A.P., Sciaini, M., & Scherer, C. (2024). Viridis (Lite) - Colorblind-Friendly Color Maps for R. viridis package version 0.6.5. |

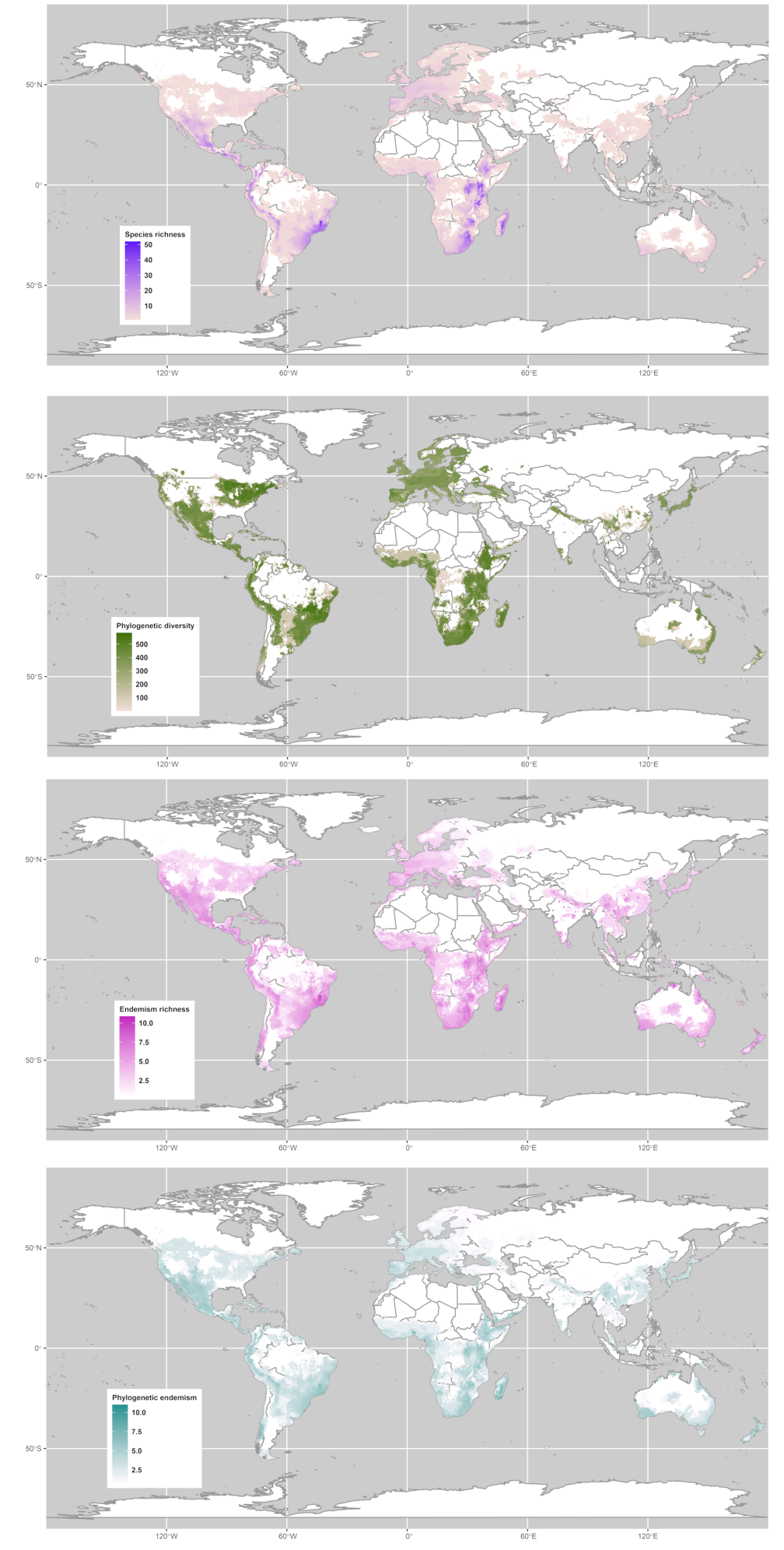

**Fig S1** Centers of diversity for DT plants according to species richness and phylogenetic diversity.

In, we found a high diversity of species in locations in Meso- and Central America (Central American Cordillera, ranging from Guatemalan Sierras Madre and de los Cuchumatanes to the Costa Rican Cordilleras Central and de Talamanca), South America (in both campos rupestres of Cadeia do Espinhaço and lowland inselbergs of Sugarloaf Land), Eastern Africa (East African Rift – Eastern Highlands – Drakensberg) and in the Malagasy Central High Plateau.

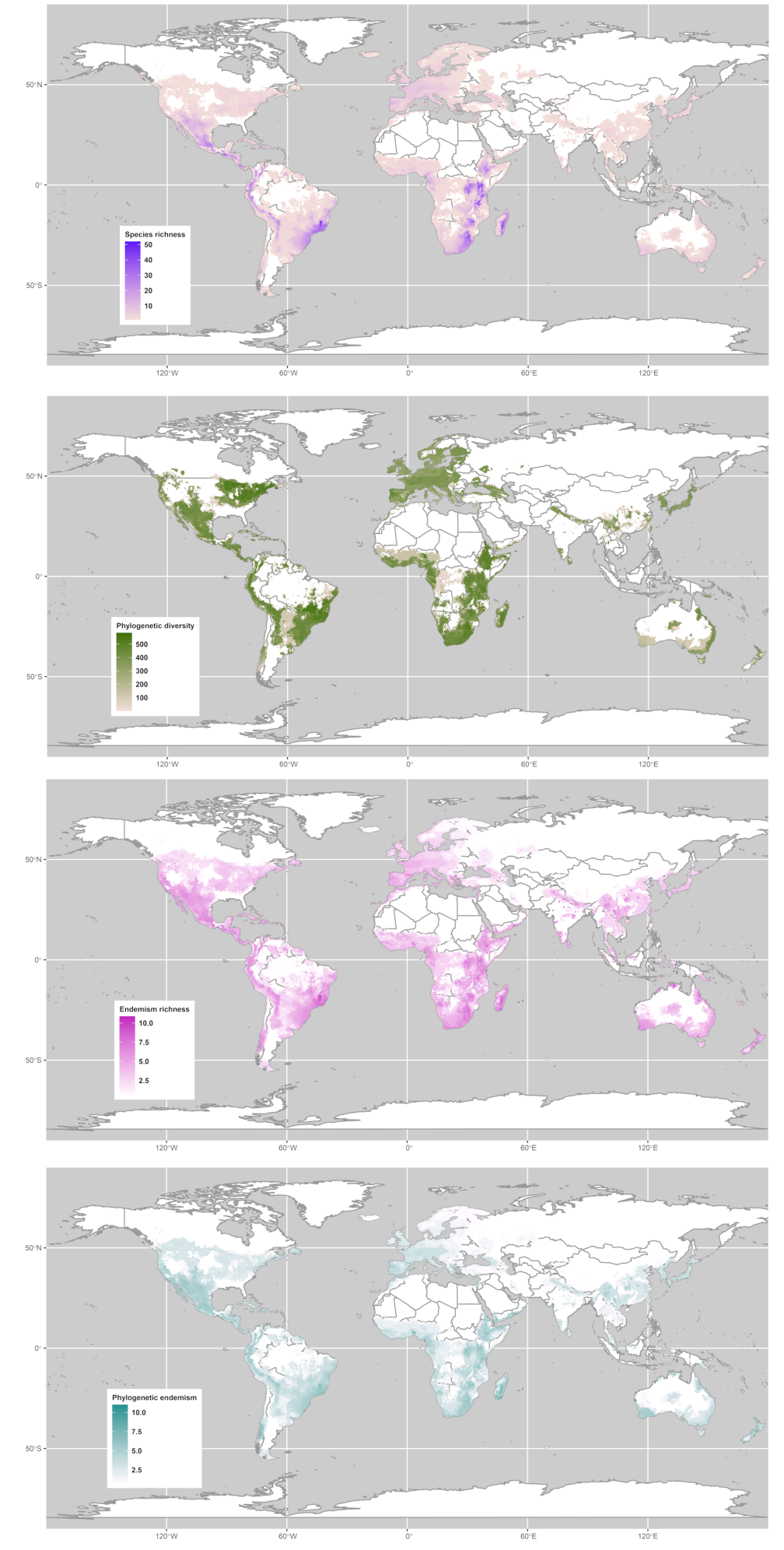

**Fig S2** Centers of endemism for DT plants according to endemism richness, and phylogenetic endemism. We found a high endemism of species in the Meso- and Central America (Mexican Sierras Madre Occidental, Madre Oriental, Madre del Sur, and Mixteca, in Mexico; extending to the Sierra Madre from Chiapas to Nicaragua and in Costa Rica ranging from Cordilleras Central, Tilarán, and Guanacaste), Caribbean islands (Cuba, Jamaica, and Hispaniola), in Center-west Sourth America (Tucumán – Bolivian province of Bolivia’s Cordillera Oriental and the Jujuy province in northwestern Argentina), east South America (Cadeia do Espinhaço – Sugarloaf Land to this continent), in Europe (the Provence region in southeastern France, continental areas of the Ionian Region in northeastern Greece, and the Belasica Range region in Greece-Bulgaria), East Africa (East African Rift, including Somalian Ogo Mountains and Ethiopian Highlands), in southern Africa (southern and western areas of the Great Escarpment, including Lesotho’s Drakensberg and Namibian Khomas Hochland), Madagascar (from Central High Plateau to the Anozy region), Mascarene Islands (Mauritius and Réunion), center-southern Arabian Mountains (ranging from the Dhofar Governorate of Oman and Mahra Governorate of Yemen), Indian Western Ghats (Nilgiri Hills), Yunnan province in China, North Australia (Northern Territory and the Wet Tropics of Queensland), and New Zealand (northeast of the Richmond Range).
